# The identity of human tissue-emigrant CD8^+^ T cells

**DOI:** 10.1101/2020.08.11.236372

**Authors:** Marcus Buggert, Laura A. Vella, Son Nguyen, Vincent Wu, Takuya Sekine, André Perez-Potti, Colby R. Maldini, Sasikanth Manne, Samuel Darko, Amy Ransier, Leticia Kuri-Cervantes, Alberto Sada Japp, Irene Bukh Brody, Martin A. Ivarsson, Laura Hertwig, Jack P. Antel, Matthew E. Johnson, Afam Okoye, Louis Picker, Golnaz Vahedi, Ernesto Sparrelid, Sian Llewellyn-Lacey, Emma Gostick, Niklas Björkström, Amit Bar-Or, Yoav Dori, Ali Naji, David H. Canaday, Terri M. Laufer, Andrew D. Wells, David A. Price, Ian Frank, Daniel C. Douek, E. John Wherry, Maxim G. Itkin, Michael R. Betts

## Abstract

Lymphocyte migration is essential for human adaptive immune surveillance. However, our current understanding of this process is rudimentary, because most human studies to date have been restricted to immunological analyses of blood and various tissues. To address this issue, we used an integrated approach to characterize tissue-emigrant immune cells in thoracic duct lymph (TDL). In humans and non-human primates, lymphocytes were by far the most abundant immune lineage population in efferent lymph, and a vast majority of these lymphocytes were T cells. Cytolytic CD8^+^ T cell subsets were clonotypically discrete and selectively confined to the intravascular circulation, persisting for months after inhibition of S1P-dependent tissue egress by FTY-720. In contrast, non-cytolytic CD8^+^ T cell subsets with stem-like epigenetic and transcriptional signatures predominated in tissues and TDL. Collectively, these data provide an atlas of the migratory immune system and define the nature of tissue-emigrant CD8^+^ T cells that recirculate via TDL.

## INTRODUCTION

The pioneering work of Gowans and colleagues taught us that lymphocytes egress continuously from peripheral tissues and lymph nodes (LNs) and return to the intravascular circulation in a unidirectional manner via the lymphatic system (Gowans, 1957, 1959; Gowans and Knight, 1964). This process of lymphocyte migration now underpins the concept of immune surveillance. The thoracic duct is the major anatomical structure that carries terminal efferent lymph to the blood stream, draining the entire subdiaphragmatic compartment and the upper left part of the body (Phang et al., 2014). A vast majority of immune cells therefore egress from lymphoid tissues (LTs) and non-lymphoid tissues (NLTs) into the thoracic duct (Kubik, 1973). Although it has been estimated that up to 3 × 10^10^ immune cells migrate daily via this route (Pabst, 1988), current knowledge is based primarily on animal studies, and the nature of lymphocyte subsets that egress from human tissues has remained obscure (Buggert et al., 2018; Fox et al., 1984; Girardet and Benninghoff, 1977; Lemaire et al., 1998; Vella et al., 2019; Voillet et al., 2018).

A series of elegant studies in rats and sheep between the 1960s and the 1980s reported that thoracic duct lymph (TDL) was composed mainly of T cells (Gowans, 1957; Mackay et al., 1988; Mackay et al., 1990; Maddox et al., 1985). Most of the cells in these models exhibited a naive phenotype and entered the efferent lymphatic system from LNs via high endothelial venules (Hall and Morris, 1965; Mackay et al., 1990; Mackay et al., 1992; Smith et al., 1970). Memory T cells were subsequently categorized into two major subsets, namely central memory T (T_CM_) cells, which proliferative vigorously in response to activation and express the LT-homing receptors CCR7 and CD62L, and effector memory T (T_EM_) cells, which are fully equipped with effector capabilities and traffic to NLTs (Sallusto et al., 1999). However, this binary classification was based on studies of intravascular lymphocytes, and more recent work has suggested greater complexity, including the existence of a T_EM_-like subset that expresses intermediate levels of CX3CR1 and patrols NLTs (Gerlach et al., 2016).

The early studies in rats by Gowans and colleagues suggested that the number of lymphocytes migrating via the thoracic duct on a daily basis was sufficient to replace the entire intravascular pool multiple times, and a later study in humans estimated this exchange rate at 48 times per day (Schick et al., 1975). However, these estimates were based on various labelling methods designed to track lymphocytes from the thoracic duct or the dilution of isotopes among intravascular lymphocytes, which do not account for the possible existence of lymphocyte populations that remain in the blood and do not recirculate via LTs or NLTs. The concept that all human T cells recirculate has also been challenged by the identification of tissue-resident memory T (T_RM_) cells (Gebhardt et al., 2009; Masopust et al., 2010; Masopust et al., 2001; Wakim et al., 2008), which form highly stable populations in solid tissues (reviewed in (Buggert et al., 2019; Szabo et al., 2019)). T_RM_ cells dominate the total pool of memory CD8^+^ T cells in NLTs (Steinert et al., 2015) and generate immediate effector responses after secondary challenge (Gebhardt et al., 2009). The protection afforded by T_RM_ cells is thought to involve via the perforin-mediated transfer of serine proteases, cytolytic activity (Masopust et al., 2001), because many murine T_RM_ cells constitutively express granzyme B (GzmB) (Masopust et al., 2006). However, this paradigm does not necessarily apply to all human T_RM_ cells (Bartolome-Casado et al., 2019; Buggert et al., 2018; Pallett et al., 2017), which instead exhibit a unique transcriptional profile compared to circulating cells (Kumar et al., 2017) and are thought to work partly as innate-like sensors (Schenkel et al., 2013). As such, it remains an open question of whether human CD8^+^ T cells with cytolytic molecule expression are constantly surveying tissues and recirculate or merely are restricted to the vasculature at steady-state.

To address these knowledge gaps, we characterized the tissue-emigrant immune system in humans and non-human primates. In contrast to prevalent models of lymphocyte trafficking in LTs and NLTs, our data show that cytolytic memory CD8^+^ T cells are confined to the intravascular circulation at steady-state, whereas stem-like memory CD8^+^ T cells survey tissues and recirculate via TDL.

## RESULTS

### Immunological atlas of human TDL

Current knowledge of the thoracic duct immune system derives primarily from older studies with limited human replicates (Fox et al., 1984; Girardet and Benninghoff, 1977; Lemaire et al., 1998). These studies also lacked the technology required to profile the functional, phenotypic, and transcriptional properties of immune cell subpopulations. We therefore set up a research protocol to acquire matched blood and TDL from a large cohort of individuals (n = 210) with clinical indications for undergoing thoracic duct cannulation. A total of 52 of these individuals between the ages of 3 months and 88 years were used for the purposes of this study (Table S1).

We first defined the major hematopoietic cell subsets in paired samples of blood and TDL (Figure 1A and Figure S1A). The majority of immune lineage cells (CD45^+^) in TDL were T cells (75%) or B cells (20%) (Figure 1B). Lower frequencies of innate lymphoid cells (ILCs) and higher frequencies of granulocytes, monocytes, dendritic cells (DCs), and hematopoietic stem cells (HSCs) were present in blood versus TDL (Figures 1B and S1B). Natural killer (NK) cells were present at similar frequencies in each compartment (Figures 1B and S1B). A direct correlation was observed between the frequency of neutrophils and the relative frequency of red blood cells in TDL (Figure S1C). Blood contamination therefore likely explained the occasional detection of neutrophils in TDL.

**Figure 1.**
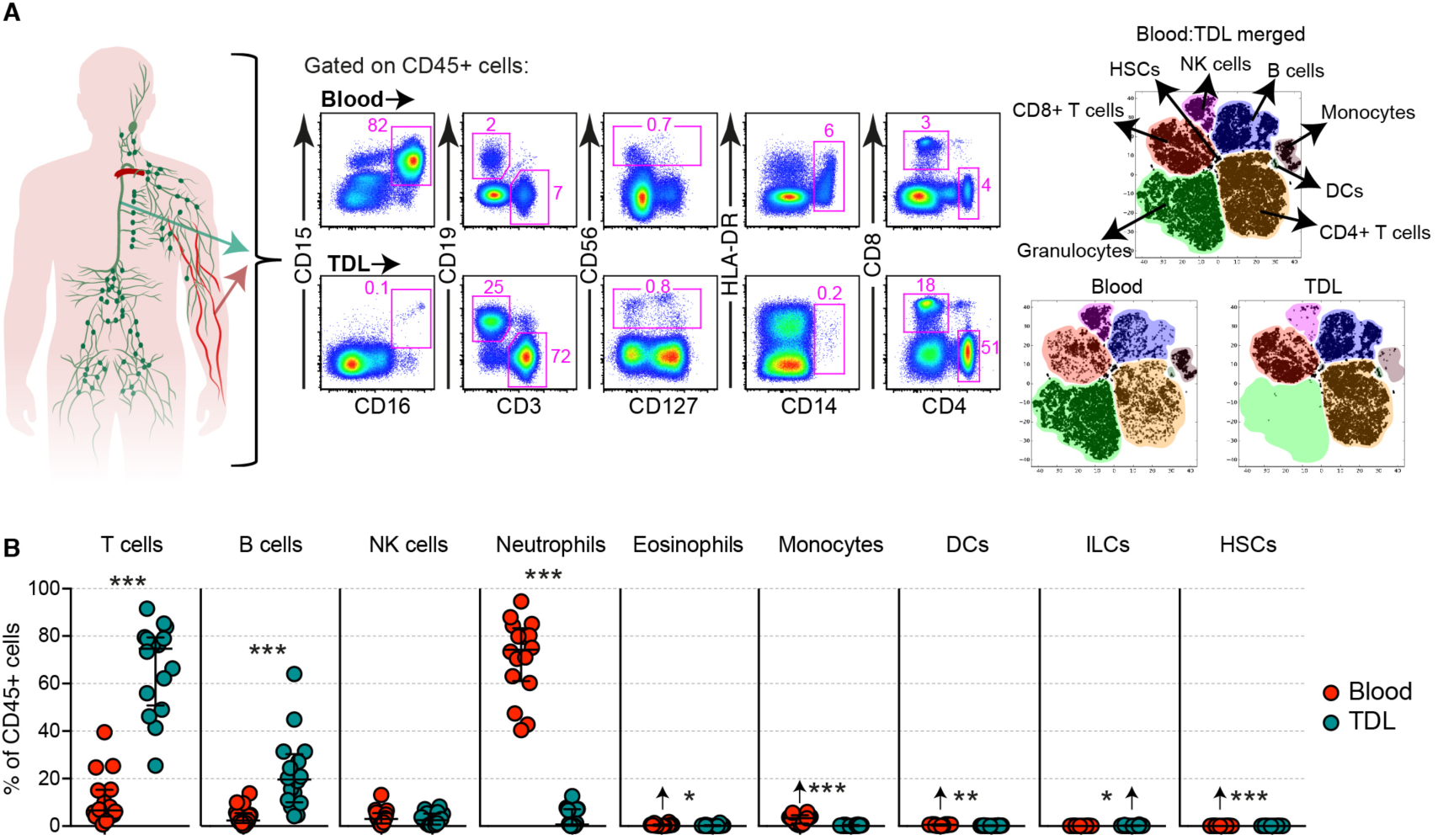
Immunological atlas of human TDL. (**A**) Representative flow cytometry plots with graphic (left) and merged tSNE plots (right) showing the differential immune lineage content of blood and TDL. (**B**) Quantification of immune cell subsets in blood and TDL. Arrows indicate higher average values. *p < 0.05, **p < 0.01, ***p < 0.001.

### Highly differentiated CD8^+^ T cells are uncommon in TDL

To extend these findings, we compared the general memory characteristics of CD8^+^ T cells in blood versus TDL based on the expression of CCR7, CD27, and CD45RA (Figure S2A). This analysis revealed that T_EM_ and terminally differentiated effector memory T (T_EMRA_) cells predominated in blood, whereas naive and T_CM_ cells predominated in TDL (Figure 2A). A similar pattern was observed in rhesus macaques based on the expression of CD28 and CD95 (Figure 2B) or CCR7 and CD95 (Figure S2B).

**Figure 2.**
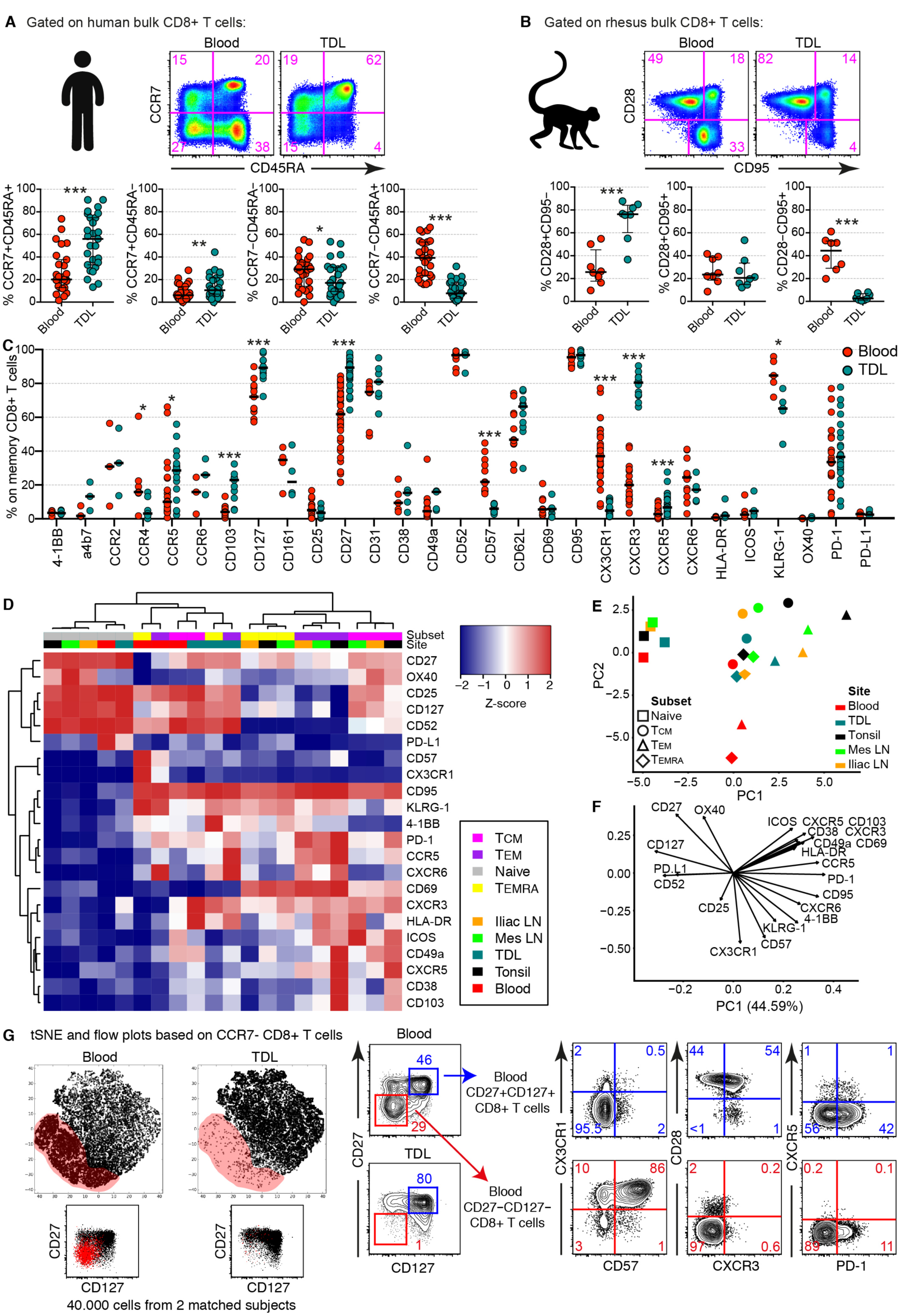
Highly differentiated CD8^+^ T cells are uncommon in TDL. (**A**) Representative flow cytometry plots (top) and summary graphs (bottom) showing the frequencies of naive and memory CD8^+^ T cell subsets in matched samples of human blood and TDL. Subsets were defined as naive (CCR7^+^CD45RA^+^), T_CM_ (CCR7^+^CD45RA^−^), T_EM_ (CCR7^−^CD45RA^−^), or T_EMRA_ (CCR7^−^CD45RA^+^). (**B**) Representative flow cytometry plots (top) and summary graphs (bottom) showing the frequencies of naive and memory CD8^+^ T cell subsets in matched samples of rhesus macaque blood and TDL. Subsets were defined as naïve (CD28^+^CD95^−^), T_CM_ (CD28^+^CD95+), or T_EM_ (CD28^−^CD95^+^). (**C**) Expression frequencies of various markers on the surface of human memory CD8^+^ T cells in matched samples of blood and TDL. (**D**) Heatmap showing the expression intensity of various markers on the surface of human memory CD8^+^ T cell subsets in blood, TDL, tonsils, iliac LNs, and mesenteric LNs. Flow cytometry data are z-score-transformed in each row. (**E**) PCA plot using the dataset in (D) to show the segregation of human naive and memory CD8^+^ T cell subsets across anatomical locations. (**F**) PCA plot showing key markers associated with the segregation observed in (E). (**G**) Merged tSNE plots (left) and flow cytometry plots (right) showing the relative absence of the CD27^−^CD127− subpopulation (red) among CCR7^−^ memory CD8^+^ T cells in human TDL. *p < 0.05, **p < 0.01, ***p < 0.001.

In more detailed flow cytometric analyses, we found that memory CD8^+^ T cells more commonly expressed certain integrins (CD103), chemokine receptors (CCR5, CXCR3, and CXCR5), and early differentiation markers (CD27 and CD127) in TDL versus blood and more commonly expressed late differentiation and effector markers (CD57, CX3CR1, and KLRG-1) in blood versus TDL (Figure 2C). Hierarchical clustering using these, and other markers revealed that each memory CD8^+^ T cell subset exhibited unique phenotypic characteristics in various lymphoid organs, including tonsils, iliac LNs, and mesenteric LNs (Figure 2D). These clusters segregated by CD69 expression, likely indicating residency in LTs (Buggert et al., 2018). In contrast, T_EM_ and T_EMRA_ cells in blood formed a discrete cluster, whereas T_CM_ cells in blood clustered with all memory subsets in TDL (Figure 2D). Principal component analysis (PCA) confirmed that T_EM_ and T_EMRA_ cells in blood segregated away from other memory clusters (Figure 2E), driven largely by the expression CD25, CD57, CX3CR1, and KLRG1 (Figure 2F). Merged t-distributed stochastic neighbor embedding (tSNE) analysis further showed that CCR7^−^ memory CD8^+^ T cells in blood often lacked CD27 and CD127 and commonly expressed markers associated with late differentiation, whereas CCR7^−^ memory CD8^+^ T cells in TDL mostly expressed CD27 and CD127 (Figure 2G).

Collectively, these analyses demonstrate that naive and early memory CD8^+^ T cells are selectively enriched in TDL and that late memory CD8^+^ T cells, defined according to standard markers, are more differentiated in blood versus TDL.

### Effector memory CD8^+^ T cells exhibit stem-like signatures in TDL

To determine the transcriptional basis of these phenotypic differences, we performed an extensive RNA-sequencing (RNA-seq) analysis of naive and memory CD8^+^ T cell subsets in blood, TDL, and mesenteric LNs (Figure 3A). A tSNE representation of these transcriptomes revealed that T_CM_ cells in blood clustered in close proximity with all memory subsets in TDL (Figure 3A). In contrast, T_EM_ and T_EMRA_ cells in blood clustered away from all other memory subsets, including T_EM_ and T_EMRA_ cells in TDL and mesenteric LNs (Figure 3A). We also identified a core signature of genes that were differentially expressed between T_EM_ and T_EMRA_ cells in blood versus TDL (fold change > 2; p < 0.05) (Table S2 and S3). Genes encoding cytolytic and effector molecules (*Gzmb*, *Gzmh*, *Cx3cr1*, *Prss23*, and *Spon2*) were upregulated among T_EM_ and T_EMRA_ cells in blood, whereas trafficking (*Ccr2*, *Ccr4*, *Ccr9*, *Itga1*, *Itga4*, *Itgb7*, and *S1pr4*) and self-renewal genes (*Tcf7*, *Il7r*, *Cd27*, *Cd28*, and *Nell2*) were upregulated among T_EM_ and T_EMRA_ cells in TDL (Figure 3B). Gene set enrichment analysis (GSEA) confirmed that signatures associated with cytolytic activity, degranulation, and differentiation were enriched among T_EM_ and T_EMRA_ cells in blood and further showed that signatures associated with cell cycle transition and telomere maintenance were enriched among T_EM_ and T_EMRA_ cells in TDL (Figure 3C).

**Figure 3.**
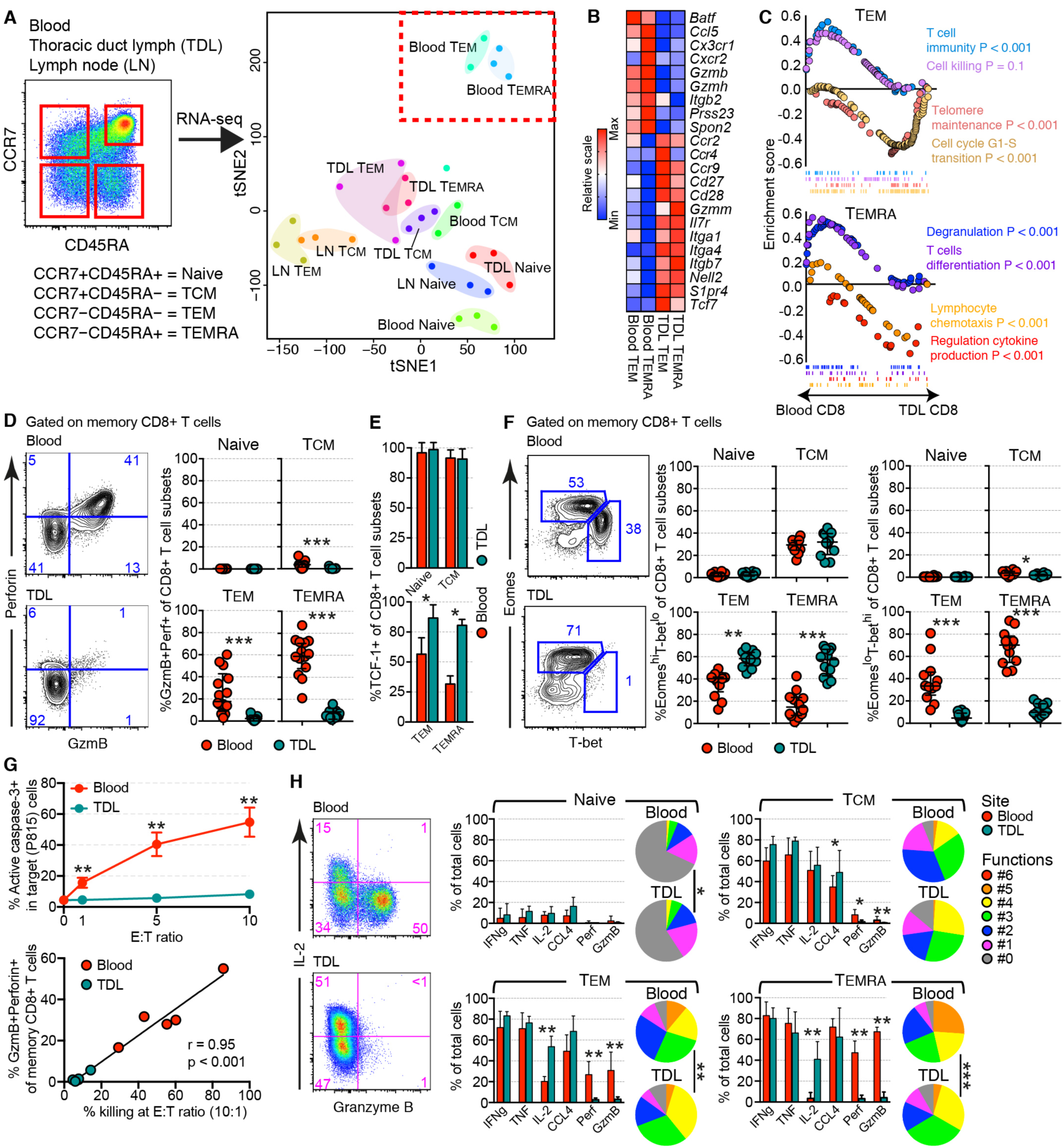
Effector memory CD8^+^ cells exhibit stem-like signatures in TDL. (**A**) Representative flow cytometry plot (left) and tSNE plot (right) showing the clustering of transcriptomes from naive and memory CD8^+^ T cell subsets in blood, TDL, and mesenteric LNs. (**B**) Heatmap showing the expression levels of selected genes among CD8^+^ T_EM_ and T_EMRA_ cells in blood and TDL. (**C**) GSEA showing the enrichment of genes associated with cytolytic activity among CD8^+^ T_EM_ and T_EMRA_ cells in blood and the enrichment of genes associated with stemness among CD8^+^ T_EM_ and T_EMRA_ cells in TDL. (**D**) Flow cytometric quantification of granzyme B and perforin among naive and memory CD8^+^ T cell subsets in blood and TDL. (**E**) Flow cytometric quantification of TCF-1 among naive and memory CD8^+^ T cell subsets in blood and TDL (n = 5). (**F**) Flow cytometric quantification of Eomes and T-bet among naive and memory CD8^+^ T cell subsets in blood and TDL. (**G**) Top: cytolytic activity of CD8^+^ T cells isolated from blood versus TDL (n = 5). Redirected killing was quantified at different effector-to-target (E:T) ratios against sensitized mastocytoma cells (P815). Bottom: correlation between cytolytic activity and the coexpression frequency of granzyme B and perforin among memory CD8^+^ T cells in blood and TDL. (**H**) Flow cytometric quantification of chemokine/cytokine production and the upregulation granzyme B and perforin among naive and memory CD8^+^ T cell subsets in blood and TDL after stimulation with PMA and ionomycin (n = 6). GzmB: granzyme B. *p < 0.05, **p < 0.01, ***p < 0.001. Functional profiles in (H) were compared using the permutation test in Simplified Presentation of Incredibly Complex Evaluations (SPICE).

Flow cytometric analyses confirmed that memory CD8^+^ T cells in TDL lacked the cytolytic proteins GzmB and perforin in humans (Figure 3D) and rhesus macaques (Figure S2C). In contrast, TCF-1, a transcription factor expressed by memory T cells with stem cell-like properties (Jeannet et al., 2010; Utzschneider et al., 2016; Zhou et al., 2010), was detected at lower frequencies among T_EM_ and T_EMRA_ cells in blood versus TDL (Figure 3E). A discrepant pattern was also observed for the T-box binding transcription factors Eomes and T-bet (Figure 3F). Specifically, Eomes^hi^T-bet^lo^ T_EM_ and T_EMRA_ cells predominated in TDL, whereas EomesloT-bethi T_EM_ and T_EMRA_ cells predominated in blood and were virtually absent in TDL (Figure 3F). Memory CD8^+^ T cells in rhesus macaques similarly expressed T-bet at much higher frequencies in blood versus TDL (Figure S2D).

In line with these results, cytolytic activity was largely confined to intravascular CD8^+^ T cells and correlated directly with expression levels of GzmB and perforin (Figure 3G). Moreover, CD8^+^ T_EM_ and T_EMRA_ cells isolated from blood upregulated GzmB and perforin after stimulation with phorbol myristate acetate (PMA) and ionomycin, unlike CD8^+^ T_EM_ and T_EMRA_ cells isolated from TDL, which instead upregulated interleukin (IL)-2 at levels equivalent to those observed among CD8^+^ T_CM_ cells isolated from blood and TDL (Figure 3H).

Collectively, these data further indicate that tissue-emigrant CCR7^−^ memory CD8^+^ T cells are less differentiated than intravascular CCR7^−^ memory CD8^+^ T cells.

### Effector memory CD8^+^ T cells are epigenetically distinct in blood and TDL

The data presented thus far suggested that cytolytic CD8^+^ T_EM_ and T_EMRA_ cells were confined to the intravascular compartment. However, it remained possible that dynamic changes in gene and protein expression allowed these cells to access tissue sites and recirculate via TDL. We therefore used the Assay of Transposase Accessible Chromatin Sequencing (ATAC-seq) to profile the open chromatin landscape of CD8^+^ T_EM_ and T_EMRA_ cells in blood, TDL, and mesenteric LNs (Figure 4A). Such epigenetic signatures are relatively stable and track prior imprints of differentiation. Global chromatin accessibility was compared among distinct cell subsets via PCA. This analysis revealed that T_EM_ and T_EMRA_ cells in TDL colocalized separately from T_EM_ and T_EMRA_ cells in blood and mesenteric LNs (Figure 4A). Certain transcription factor family motifs, most notably T-bet (TBOX) and RUNX, were enriched among T_EM_ and T_EMRA_ cells in blood versus TDL (Figure 4B). Likewise, specific open chromatin regions (OCRs) and motif families (RUNX, bZIP, and bHLH) next to effector genes (*Gzmh*, *Gzmb*, *Tbx21*, and *Prf1*) were more accessible among T_EM_ and T_EMRA_ cells in blood versus TDL (Figure 4C). In additional experiments, we incubated peripheral blood mononuclear cells (PBMCs) and TDL mononuclear cells (TDLMCs) with plasma or TDL fluid to exclude the possibility that soluble factors could regulate the expression of cytolytic molecules, exemplified by GzmB. No upregulation of GzmB was observed in either compartment under either of these conditions (Figure S2E).

**Figure 4.**
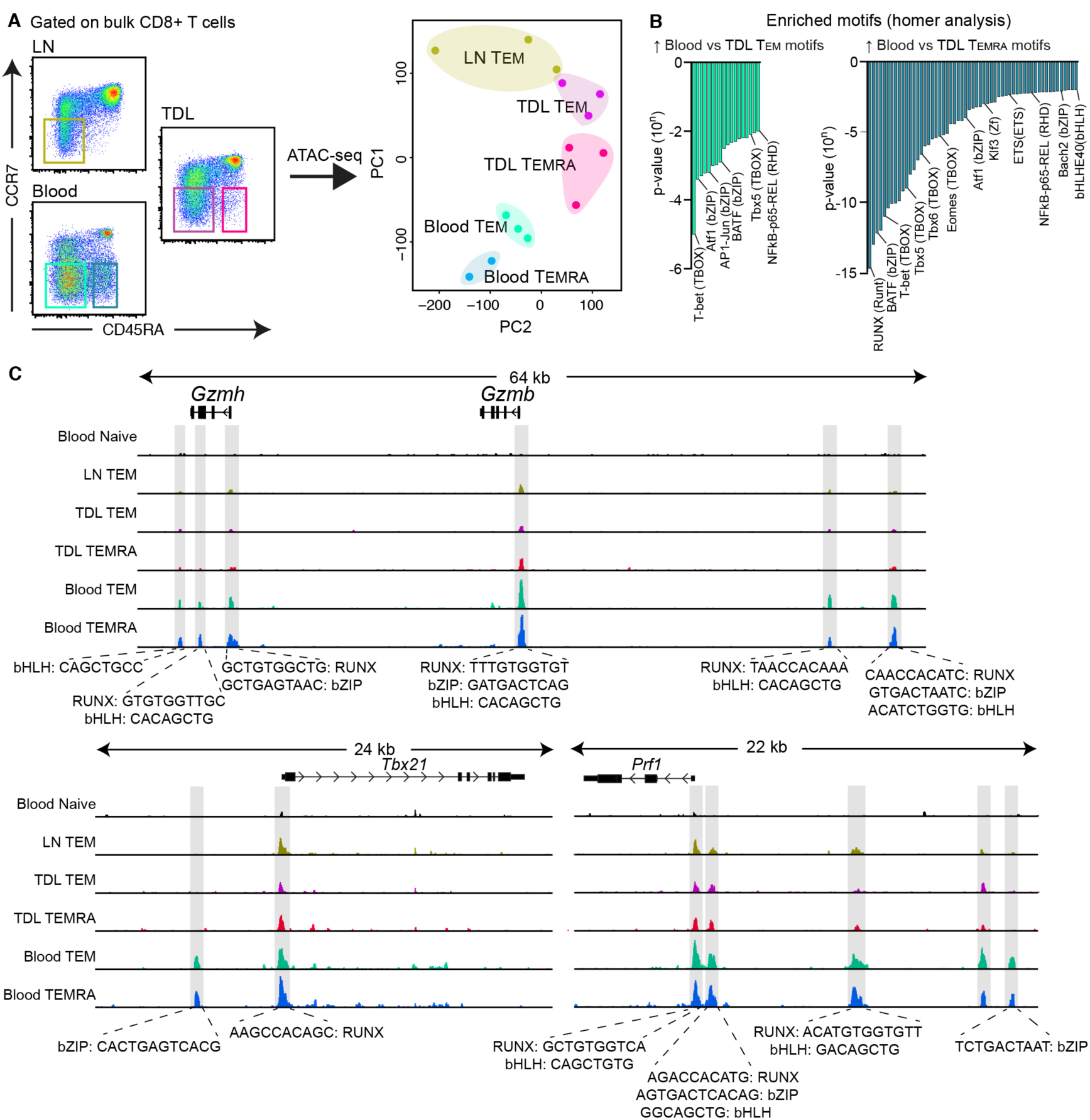
Effector memory CD8^+^ T cells are epigenetically distinct in blood and TDL. (**A**) Flow cytometric gating strategy for cell sorting (left) and PCA plot based on the corresponding ATAC-seq data (right) showing global chromatin accessiblity clusters for CD8^+^ T_EM_ and T_EMRA_ cells in blood, TDL, and mesenteric LNs. (**B**) Transcription factor motifs enriched among CD8^+^ T_EM_ (left) and T_EMRA_ cells (right) in blood versus TDL. (**C**) ATAC-seq tracks of the *Gzmh*, *Gzmb*, *Tbx21*, and *Prf1* loci among naive and memory CD8^+^ T cell subsets in blood, TDL, and mesenteric LNs.

Collectively, these results suggest that effector memory CD8^+^ T cells might be derived from separate maturational lineages in blood and TDL.

### Cytolytic and non-cytolytic CD8^+^ T cells are clonotypically divergent

To seek further evidence of divergent maturational pathways, we used an unbiased molecular approach to sequence T cell receptor (TCR) β gene (*TRB*) rearrangements (TCR-seq) expressed among memory CD8^+^ T cell subsets in blood and TDL. Pilot experiments revealed the presence of cytolytic precursors and cytolytic effectors in the T_EM_ and T_EMRA_ subsets (Figure S3A), consistent with previous work (Patil et al., 2018). We therefore used CX3CR1 as a surrogate marker to identify cytolytic effector CD8^+^ T cells (Figure 5A; (Bottcher et al., 2015). As insufficient numbers of CX3CR1^+^ cells were present in TDL (Figures 5A, S3B, and S3C), we compared CCR7^−^ memory CX3CR1^+^ cells in blood with CCR7^+^ memory CX3CR1^−^ (*i.e.*, predominantly T_CM_) and CCR7^−^ memory CX3CR1^−^ (*i.e.*, predominantly T_EM_) cells in blood and TDL (Figure 5B). Repertoire diversity was substantially lower among CCR7^−^ memory CX3CR1^+^ cells in blood versus all other memory subsets in blood and TDL (Figure 5C). A near-perfect correlation was observed between the frequencies of T_CM_ clonotypes in blood and TDL, indicating that T_CM_ cells in blood were highly representative of T_CM_ cells that egressed from tissues and recirculated via TDL (Figures 5D and 5E). In contrast, limited clonotypic correlation was observed between the T_EM_ subsets in blood and TDL, and the cytolytic effector repertoires were largely dissimilar, based on frequencies, from all other memory subsets in blood and TDL (Figures 5D and 5E).

**Figure 5.**
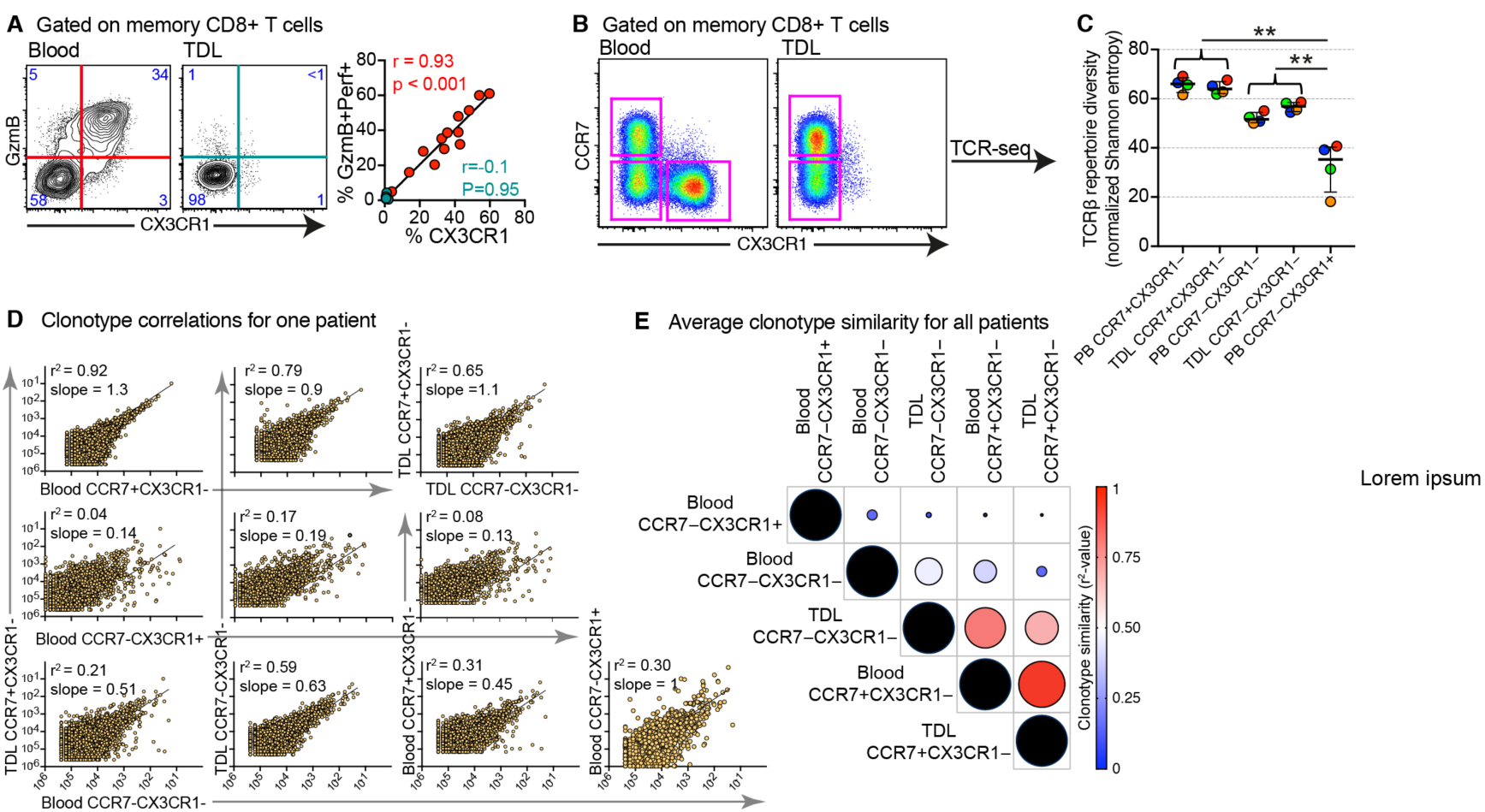
Cytolytic and non-cytolytic CD8^+^ T cells are clonotypically divergent. (**A**) Representative flow cytometry plots (left) and scatter graph (right) showing the expression frequencies of granzyme B and perforin versus CX3CR1 among memory CD8^+^ T cells. (**B**) Flow cytometric gating strategy for sorting memory CD8^+^ T cell subsets based on the expression of CCR7 and CX3CR1. (**C**) TCRβrepertoire diversity calculated for each memory CD8^+^ T cell subset using normalized Shannon entropy. (**D**) Representative clonotype frequency correlations among memory CD8^+^ T cell subsets in blood and TDL. (**E**) Bubble plot showing pairwise comparisons of clonotype similarity among CD8^+^ T cell subsets in blood and TDL (n = 4). GzmB: granzyme B. **p < 0.01.

### Cytolytic CD8^+^ T cells are selectively retained in the intravascular circulation

To determine if cytolytic CD8^+^ T cells were present in tissues, we collected unpaired human LTs (tonsils and mesenteric LNs) and NLTs (colon, decidua, and liver). Cytolytic CD8^+^ T cells were common in blood and rare in all other anatomical locations (Figure 6A). Non-resident CD8^+^ T cells (CD69−) were predominantly CCR7^+^ in LTs and CCR7^−^ in NLTs (Figure 6B), consistent with previous work ((Sallusto et al., 1999). Most non-resident CD8^+^ T_EM_ cells in the decidua also lacked CX3CR1 (Figure 6C; (Gerlach et al., 2016). In addition, we collected matched samples of blood entering the liver via the portal vein and leaving the liver via the central hepatic veins. Similar frequencies of non-resident cytolytic CD8^+^ T cells (CD57^+^) were present before and after organ transit (Figure S5D). These results were confirmed using matched samples of arterial and venous blood from other donors (Figure S5E).

**Figure 6.**
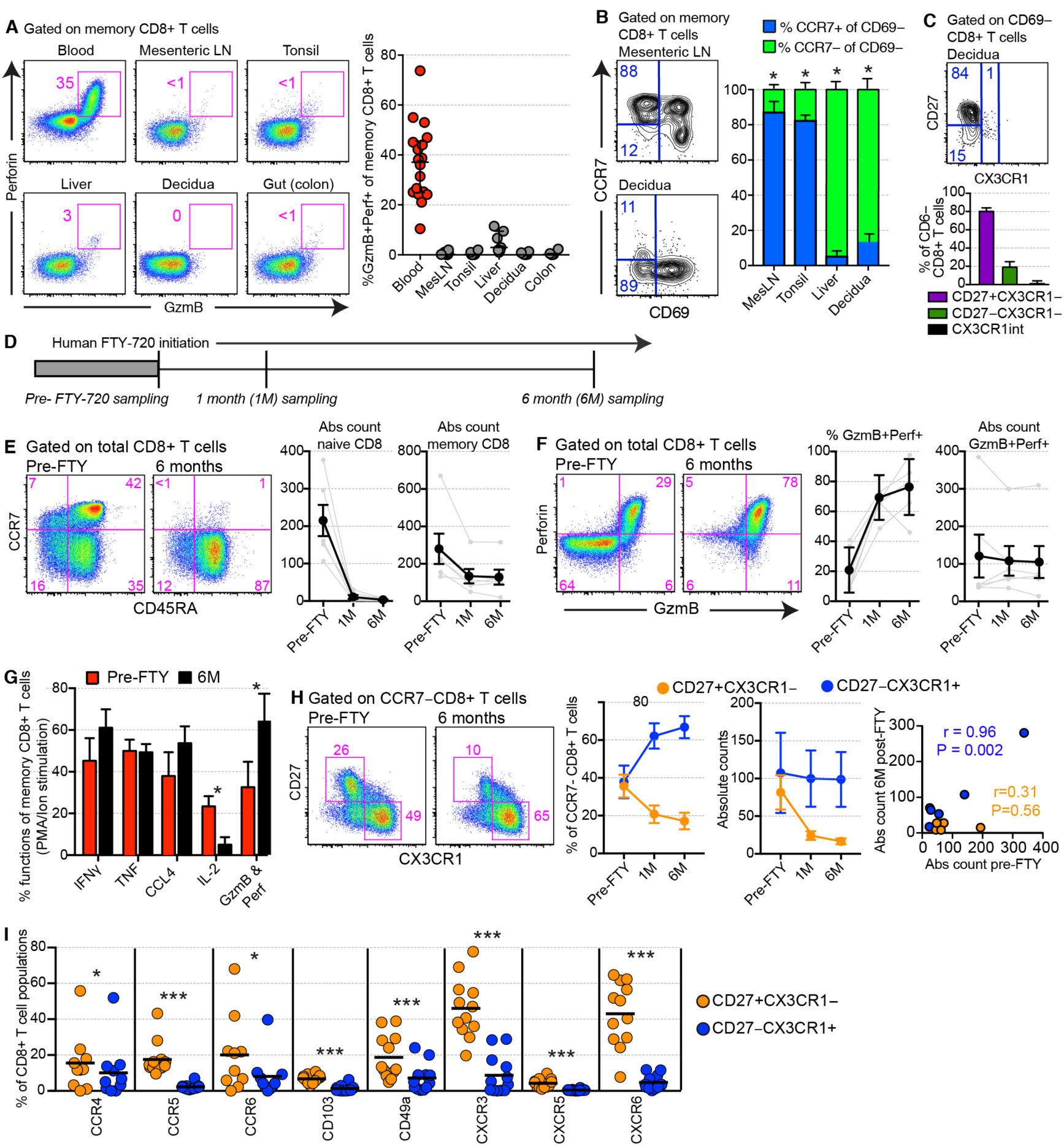
Cytolytic CD8^+^ T cells are selectively retained in the intravascular circulation. (**A**) Representative flow cytometry plots (left) and summary graphs (right) showing the coexpression frequencies granzyme B and perforin among memory CD8^+^ T cells in blood, LTs, and NLTs. All comparisons versus blood were highly significant. (**B**) Representative flow cytometry plots (left) and summary graphs (right) showing the expression frequencies of CCR7 and CD69 among memory CD8^+^ T cells in LTs and NLTs (n = 15). (**C**) Representative flow cytometry plots (top) and summary graphs (bottom) showing the expression frequencies of CD27 and CX3CR1 among non-resident CD8^+^ T_EM_ cells (CD69^−^) in endometrial tissue (n = 4). (**D**) Schematic representation of the fingolimod (FTY-720) study. Blood samples were drawn before,1 month after (1M), and 6 months after (6M) the initiation of FTY-720. (**E**) Representative flow cytometry plots (left) and summary graphs (right) showing the frequencies (left) and absolute numbers (right) of naive and memory CD8^+^ T cells over the course of the study. (**F**) Representative flow cytometry plots (left) and summary graphs (right) showing the frequencies (left) and absolute numbers (right) of cytolytic CD8^+^ T cells (granzyme B^+^perforin^+^) over the course of the study. (**G**) Functional profiles of memory CD8^+^ T cells in response to stimulation with PMA and ionomycin before and 6 months after the initiation of FTY-720 (n = 6). (**H**) Representative flow cytometry plots (left) and summary graphs (middle) showing the persistence of intravascular cytolytic CD8^+^ T cells (CCR7^−^CD27^−^CX3CR1^+^) after the initiation of FTY-720 (n = 6). Right: correlation between the absolute numbers of intravascular cytolytic CD8^+^ T cells (CCR7^−^CD27^−^CX3CR1^+^) before and 6 months after the initiation of FTY-720. (**I**) Expression frequencies of various trafficking receptors among cytolytic (CCR7^−^CD27^−^CX3CR1^+^) and non-cytolytic effector memory CD8^+^ T cells (CCR7^−^CD27^+^CX3CR1^−^). GzmB: granzyme B; NS: non-significant. *p < 0.05, **p < 0.01, ***p < 0.001.

To investigate the migratory capabilities of cytolytic CD8^+^ T cells more directly, we obtained longitudinal samples of venous blood from multiple sclerosis patients undergoing treatment with fingolimod (FTY-720), a drug that prevents sphingosine-1-phosphate (S1P)-mediated lymphocyte egress from LTs and NLTs (Figure 6D). The absolute numbers and frequencies of naive CD4^+^ (Figures S4A and S4B) and CD8^+^ T cells (Figures 6E, S4C, and S4D) fell within the first month and remained low with continuous treatment over a period of 6 months. A similar but less dramatic pattern was observed for memory CD4^+^ (Figures S4A and S4B) and CD8^+^ T cells (Figures 6E, S4C, and S4D). The residual memory CD4^+^ T cell populations exhibited a predominant T_EM_ phenotype (Figures S4A and S4B), and the residual CD8^+^ T cell populations exhibited a predominant T_EMRA_ phenotype (Figures 6E, S4C, and S4D). Moreover, approximately 80% of the residual memory CD8^+^ T cells expressed GzmB and perforin after 6 months, and the absolute counts of memory CD8^+^ T cells that expressed both GzmB and perforin were not significantly lower after 6 months (Figure 6F). These changes were reflected in the activation-induced functional profiles of memory CD8^+^ T cells, which showed lower frequency of IL-2-producers and higher frequency of GzmB and perforin-producers at 6 months of FTY-720 treatment compared to matched samples obtained before initiation of treatment (Figure 6G). In line with these findings, the absolute numbers of CCR7^−^CD27^−^CX3CR1^+^ memory CD8^+^ T cells remained stable over time, whereas the absolute numbers and frequencies of CCR7^−^CD27^+^CX3CR1^−^ memory CD8^+^ T cells declined over time, leading to a proportionate increase in the frequencies of CCR7^−^CD27^−^CX3CR1^+^ memory CD8^+^ T cells (Figure 6H). Various chemokine receptors and other homing molecules were also expressed less frequently among CCR7^−^CD27^−^CX3CR1^+^ memory CD8^+^ T cells versus CCR7^−^CD27^+^CX3CR1^−^ memory CD8^+^ T cells (Figure 6I).

Collectively, these data suggest that cytolytic CD8^+^ T cells are selectively retained in the intravascular circulation and rarely migrate through LTs or NLTs at steady-state.

### Cytolytic and non-cytolytic effector memory CD8^+^ T cells are transcriptionally and epigenetically distinct in the intravascular circulation

We next wanted to identify an intravascular signature of effector memory CD8^+^ T cells with a recirculating non-cytolytic phenotype (CCR7^−^CD27^+^CX3CR1^−^) and vascular restricted cytolytic phenotype (CCR7^−^CD27^−^CX3CR1^+^), by initially employing RNA-seq analysis (Figure 7A). Core signatures of differentially expressed genes (fold change > 2; p < 0.05) were identified among the CCR7^−^CD27^+^CX3CR1^−^ (633 genes) and CCR7^−^CD27^−^CX3CR1^+^ subsets (545 genes), including transcripts encoding various costimulatory and effector molecules, integrins, trafficking receptors, and transcription factors (Figures 7A and 7B and Table S4). Many of the integrins and trafficking receptors were differentially expressed at the protein level (Figures 6K and S5A). Gene ontology (GO) analysis revealed that CCR7^−^CD27^+^CX3CR1^−^ cells were enriched for signatures associated with adhesion, migration, and proliferation, whereas CCR7^−^CD27^−^CX3CR1^+^ cells were enriched for signatures associated with granule localization, epigenetic regulation, and lymphocyte differentiation (Figure 7C). The gene signature of CCR7^−^CD27^+^CX3CR1^−^ cells also overlapped with the gene signature of memory precursors (IL-7R^hi^) in the CD8^+^ lineage, determined using GSEA (Figure 7D). Accordingly, expression levels of IL-7R and TCF-1 readily distinguished the CCR7^−^CD27^+^CX3CR1^−^ and CCR7^−^CD27^−^CX3CR1^+^ populations (Figure 7E), and CCR7^−^CD27^+^CX3CR1^−^ cells proliferated *in vitro* more vigorously than CCR7^−^CD27^−^CX3CR1^+^ cells (Figure 7F).

**Figure 7.**
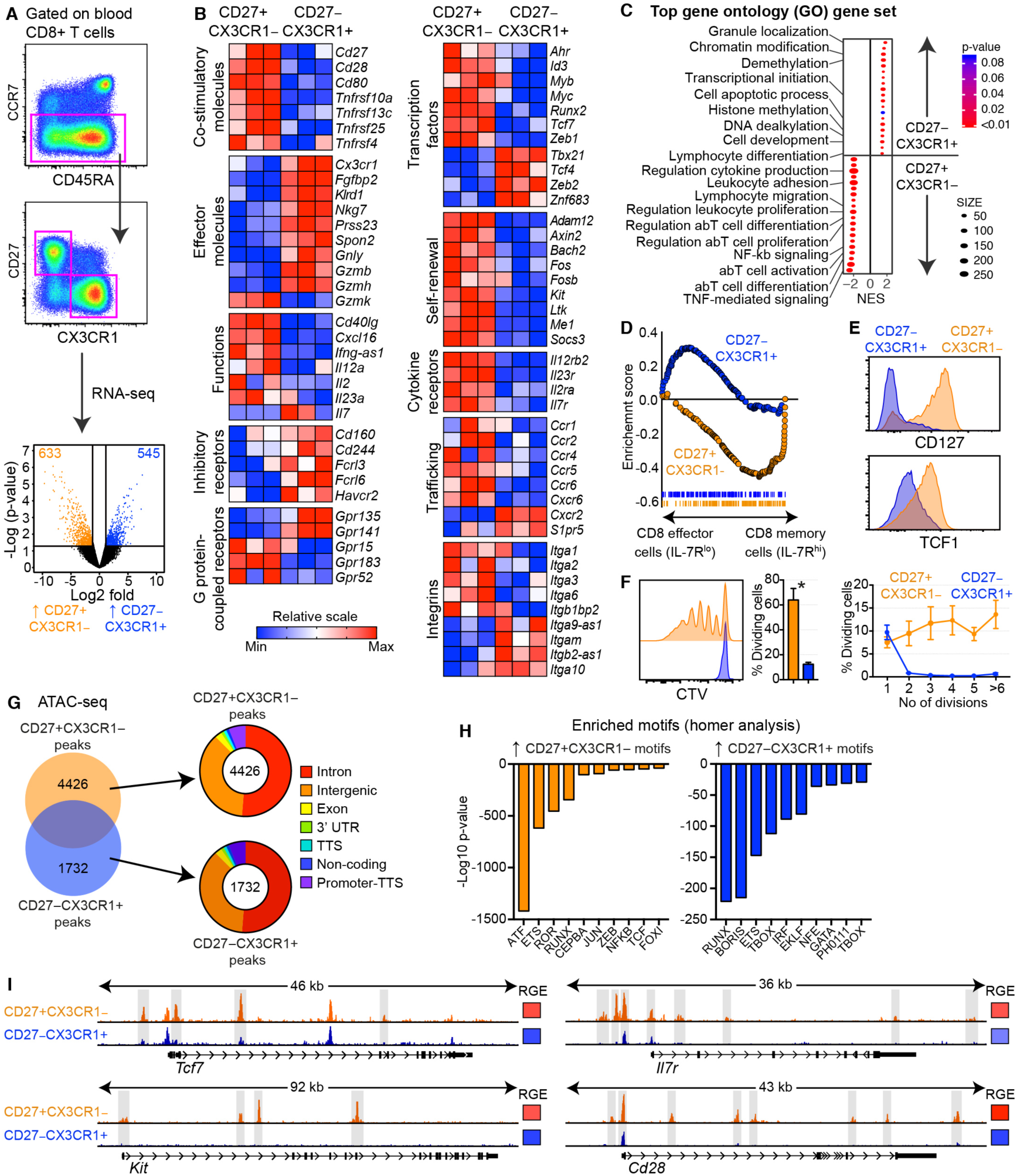
Cytolytic and non-cytolytic effector memory CD8^+^ T cells are transcriptionally and epigenetically distinct in the intravascular circulation. (**A**) Flow cytometric gating strategy for cell sorting (top) and volcano plot (bottom) showing the number of differentially expressed genes (fold change > 2; p < 0.05) among CCR7^−^CD27^+^CX3CR1^−^ versus CCR7^−^CD27^−^CX3CR1^+^ memory CD8^+^ T cells. (**B**) Heatmaps showing the differential expression of gene modules between CCR7^−^CD27^+^CX3CR1^−^ and CCR7^−^CD27^−^CX3CR1^+^ memory CD8^+^ T cells. Data are z-normalized in each panel. (**C**) GSEA comparing signatures from each population versus the GO database (Broad Institute). (**D**) Enrichment analysis of memory (IL-7R^hi^) and effector signatures (IL-7R^lo^) among CCR7^−^CD27^+^CX3CR1^−^ and CCR7^−^CD27^−^CX3CR1^+^ memory CD8^+^ T cells. (**E**) Representative flow cytometry histograms for each population showing expression levels of CD127 and TCF-1. (**F**) Flow cytometry plot (left) and summary graphs (middle and right) from a representative CellTrace Violet (CTV) dilution experiment comparing CCR7^−^CD27^+^CX3CR1^−^ versus CCR7^−^CD27^−^CX3CR1^+^ memory CD8^+^ T cells. (**G**) Genomic distribution of OCRs comparing peaks from CCR7^−^CD27^+^CX3CR1^−^ versus CCR7^−^CD27^−^CX3CR1^+^ memory CD8^+^ T cells. (**H**) Bar graphs showing the enrichment of specific transcription factor binding motifs in each population. (**I**) ATAC-seq tracks demonstrating the enrichment of OCRs adjacent to *Tcf7*, *Kit*, *Il7r*, and *Cd28* among CCR7^−^CD27^+^CX3CR1^−^ memory CD8^+^ T cells. Heatmaps on the right show relative gene expression (RGE) from the RNA-seq analysis (blue = min; red = max). RNA-seq: n = 3 ATAC-seq: n = 3. *p < 0.05.

To determine if these transcriptional differences were associated with distinct transcription factor motifs and OCRs, we used ATAC-seq to map the corresponding epigenetic landscapes. CCR7^−^CD27^+^CX3CR1^−^ cells contained more OCRs (n = 4,426) than CCR7^−^CD27^−^CX3CR1^+^ cells (n = 1,732) (Tables S5 and S6). Most of these OCRs were located in introns or intergenic regions (Figure 7G). Transcription factor motif analysis revealed that OCRs unique to the CCR7^−^CD27^+^CX3CR1^−^ subset contained binding sites for the ATF family, whereas OCRs unique to the CCR7^−^CD27^−^CX3CR1^+^ subset contained binding sites for the RUNX, BORIS, and TBOX families (Figure 7H). Specific regions next to genes associated with self-renewal (*Cd28*, *Il7r*, *Kit*, and *Tcf7*) and effector functions (*Gzmh*, *Gzmb*, *Prf1*, and *Cx3cr1*) were also differentially accessible between the CCR7^−^CD27^+^CX3CR1^−^ and CCR7^−^CD27^−^CX3CR1^+^ subsets (Figures 7I and S5B).

Collectively, these results indicate that intravascular CCR7^−^CD27^+^CX3CR1^−^ memory CD8^+^ T cells, like their non-cytolytic T_EM_ and T_EMRA_ counterparts in TDL, are transcriptionally and epigenetically distinct from intravascular CCR7^−^CD27^−^CX3CR1^+^ memory CD8^+^ T cells.

### Virus-specific CD8^+^ T cells rarely express cytolytic molecules in TDL

To confirm these findings in the context of antigen specificity, we obtained paired samples of blood and TDL from individuals on continuous antiretroviral therapy for chronic HIV infection (n = 11). In matched samples, CMV-specific CD8^+^ T cells were proportionately more common in blood versus TDL (median ratio = 2.8), whereas HIV-specific CD8^+^ T cells, which displayed a transitional memory phenotype (CCR7^−^CD27^+^CD45RO^+^) in both compartments (Figure S6A(Buggert et al., 2014), were proportionately less common in blood versus TDL (median ratio = 0.79) (Figure 8A). The expression of cytolytic molecules was confined almost exclusively to intravascular virus-specific CD8^+^ T cells (Figure 8B). Accordingly, most CMV-specific CD8^+^ T cells in blood displayed a CCR7^−^CD27^−^ phenotype and expressed GzmB, whereas most CMV-specific CD8^+^ T cells in TDL displayed a CCR7^−^CD27^+^ phenotype and lacked GzmB (Figure 8C). Moreover, a direct correlation was observed between the frequency of intrasvascular virus-specific CD8^+^ T cells that expressed both GzmB and perforin and the relative frequency of virus-specific CD8^+^ T cells in blood versus TDL (Figure 8C). In line with these findings, relatively few EomesloT-bethi virus-specific CD8^+^ T cells were present in TDL (Figure 8D).

**Figure 8.**
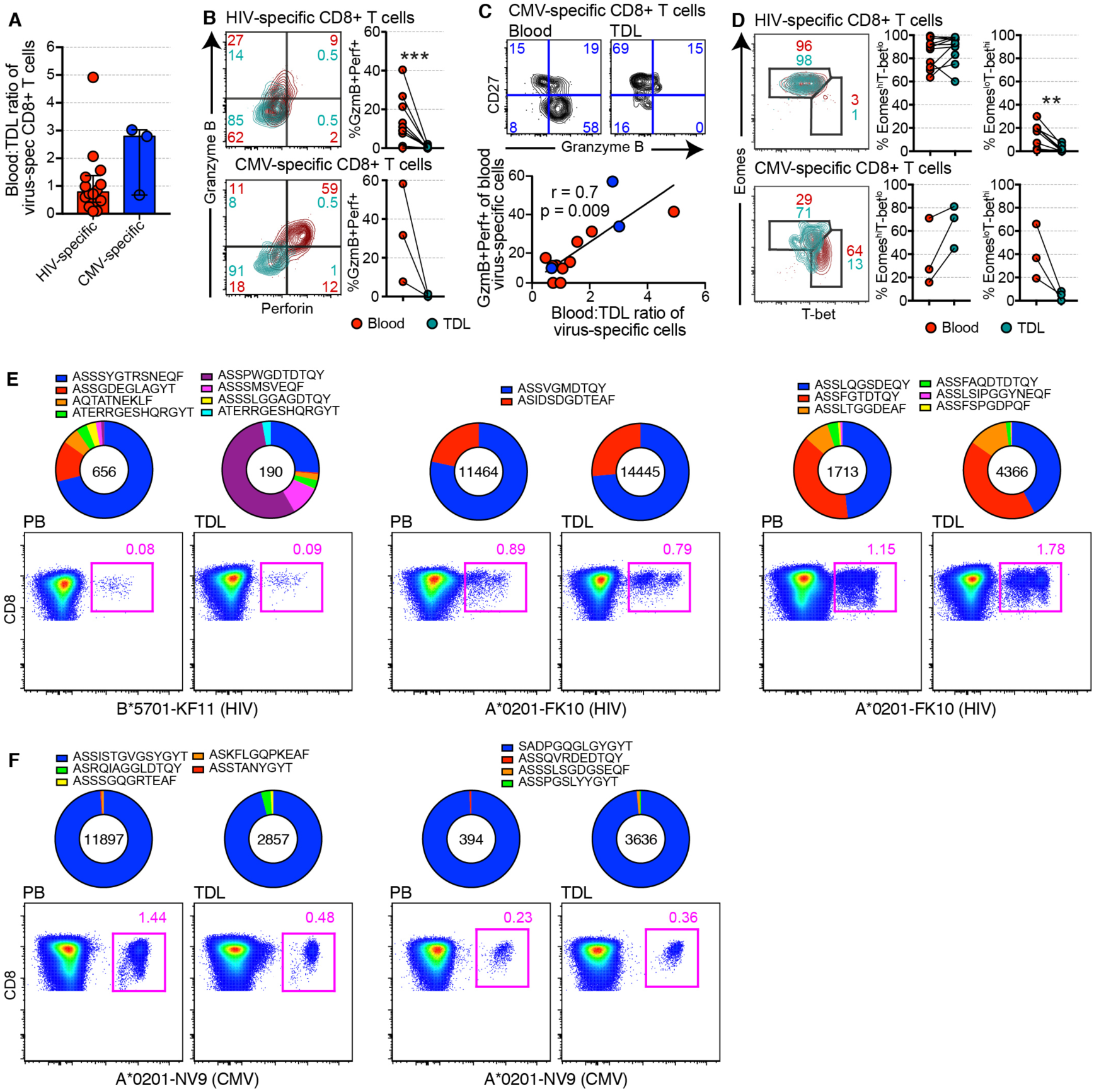
Virus-specific CD8^+^ T cells rarely express cytolytic molecules in TDL. (**A**) MHC class I tetramer-based quantification of virus-specific CD8^+^ T cells in blood and TDL (shown as ratios). (**B**) Representative flow cytometry plots (left) and summary graphs (right) showing the expression frequencies of granzyme B and perforin among virus-specific CD8^+^ T cells in blood and TDL. (**C**) Top: representative flow cytometry plots showing the expression of CD27 and granzyme B among CMV-specific CD8^+^ T cells in blood and TDL. Bottom: correlation between the coexpression frequency of granzyme B and perforin and the frequency of virus-specific CD8^+^ T cells in blood versus TDL. (**D**) Representative flow cytometry plots (left) and summary graphs (right) showing the expression frequencies of Eomes and T-bet among virus-specific CD8^+^ T cells in blood and TDL. (**E**, **F**) Clonotype distribution among virus-specific CD8^+^ T cells in blood and TDL. The number in each circle indicates the total number of TCRs. Population frequencies are shown in the corresponding flow cytometry plots. *p < 0.05, **p < 0.01, ***p < 0.001.

To determine the clonal origins of these anatomically and functionally distinct virus-specific CD8^+^ T cell populations, we sequenced the corresponding *TRB* gene rearrangements in blood and TDL. As expected, the CMV-specific repertoires were heavily skewed in favor of a single clonotype, whereas the HIV-specific repertoires incorporated higher frequencies of subdominant clonotypes (Figures 8E and 8F). A very high degree of clonotypic overlap was also observed between matched specificities in blood and TDL (Figure S6B).

Collectively, these findings demonstrate that functionally and phenotypically distinct virus-specific CD8^+^ T cells with shared clonal ancestries recirculate in blood and TDL.

## DISCUSSION

Elegant work over several decades has enhanced our understanding of anatomically localized and intravascular subsets of human memory CD8^+^ T cells (Kumar et al., 2017; Sallusto et al., 1999; Sathaliyawala et al., 2013). In contrast, relatively little information has emerged about the nomadic CD8^+^ T cell populations that recirculate continuously and survey peripheral tissue sites. In this body of work, we characterized the functional, phenotypic, and transcriptional properties of tissue-emigrant CD8^+^ T cells in humans and rhesus macaques. Our key finding was that not all memory CD8^+^ T cell subsets migrated through LTs or NLTs. Effector memory CD8^+^ T cell subsets that expressed high levels of cytolytic molecules were rarely detected in tissues or TDL at steady-state. Instead, these cells were selectively retained in the intravascular circulation, persisting for months after inhibition of S1P-dependent tissue egress by FTY-720. In addition, we identified previously unrecognized progenitor-like subsets of CD8^+^ T cells within the classically defined T_EM_ and T_EMRA_ compartments that migrated through tissues and TDL.

Since the pioneering work of Gowans and colleagues (Gowans, 1957, 1959; Gowans and Knight, 1964), our understanding of human lymphocyte recirculation and tissue egress has been primarily shaped via studies in animal models (Hall and Morris, 1965; Mackay et al., 1996; Mackay et al., 1988; Mackay et al., 1990; Mackay et al., 1992; Miller and Sprent, 1971; Smith et al., 1970; Sprent, 1973). However, these studies lacked the technology to examine migratory lymphocyte populations in detail, and the reported findings cannot necessarily be extended to humans or non-human primates (Beura et al., 2016). Only a limited number of studies have been conducted on lymphocyte trafficking via efferent lymph in humans (Buggert et al., 2018; Fox et al., 1984; Girardet and Benninghoff, 1977; Klicznik et al., 2019; Lemaire et al., 1998; Vella et al., 2019; Voillet et al., 2018). In contrast to these efforts, we used state-of-the-art technology to map the entire immune system in TDL. A vast majority of immune cells in efferent lymph were T cells (75%). Other lineages commonly found in the intravascular circulation, such as monocytes, neutrophils, and myeloid-derived DCs, were rarely detected in clean samples of efferent lymph, concordant with their distinct origins and differentiation pathways (Furze and Rankin, 2008; Zhao et al., 2018), whereas NK cells were present at largely equivalent frequencies in blood and TDL (Fox et al., 1984).

In contrast to landmark ovine studies, which reported that >90% of all T cells in efferent lymph displayed a naive phenotype (Mackay et al., 1988; Mackay et al., 1990), we found that naive T cell frequencies ranged from <20% to >90% in human and non-human TDL. Classically defined CD8^+^ T_EM_ and T_EMRA_ cells were also present in efferent lymph, but unlike the corresponding intravascular subsets, these cells were not highly differentiated and rarely expressed cytolytic molecules. Intralymphatic CD8^+^ T_EM_ and T_EMRA_ cells instead exhibited progenitor-like characteristics, including the ability to proliferate in response to activation and produce IL-2. These cells were functionally and transcriptionally equivalent to intravascular T_CM_ cells and lacked the tissue residency marker CD69. Accordingly, genes and pathways associated cell cycle transition, chemotaxis, self-renewal, telomere maintenance, and trafficking were upregulated among intralymphatic versus intravascular CD8^+^ T_EM_ and T_EMRA_ cells, whereas genes and pathways associated with cytolytic activity and effector functionality were downregulated among intralymphatic versus intravascular CD8^+^ T_EM_ and T_EMRA_ cells. Moreover, intralymphatic CD8^+^ T_EM_ and T_EMRA_ cells exhibited a stem-like Eomes^hi^T-bet^lo^TCF-1^hi^ transcriptional profile (Im et al., 2016; Siddiqui et al., 2019), unlike intravascular CD8^+^ T_EM_ and T_EMRA_ cells (McLane et al., 2013). Of note, murine studies have identified a migratory CCR7^−^CX3CR1^int^ population in blood, efferent lymph, and tissues (Gerlach et al., 2016). A vast majority of human CCR7^−^ memory CD8^+^ T cells in efferent lymph and tissues nonetheless displayed a CD27^+^CX3CR1^−^ phenotype, conceivably indicating adaptations to recurrent pathogen exposure over a long period of time.

Memory CD8^+^ T cells rarely expressed GzmB and perforin in TDL and almost never expressed GzmB and perforin in LTs and NLTs. A similar pattern was observed for persistent virus-specific CD8^+^ T cells in TDL. These findings suggested that cytolytic CD8^+^ T cells were largely confined to the intravascular circulation, both under physiological conditions and during chronic infection with CMV or HIV. Several observations further argued against the possibility that cytolytic molecules were transiently downregulated as memory CD8^+^ T cells entered the peripheral tissues, a process that would require granule autophagy, extensive transcriptional changes, and epigenetic remodeling at effector gene loci, with reversal of these adaptations after transit and continual plasticity to maintain a state of phenotypic oscillation in response to microenvironmental signals. First, epigenetic analyses confirmed the presence of multiple OCRs adjacent to effector genes, such as *Gzmb*, *Gzmh*, *Prf1*, and *Tbx21*, among intravascular but not among intralymphatic CD8^+^ T_EM_ and T_EMRA_ cells, suggesting a stable phenotype rather than a transitory state. Second, mixing experiments showed that neither efferent lymph nor plasma altered the expression of GzmB expression among CD8^+^ T_EM_ and T_EMRA_ cells isolated from blood or TDL. Third, the clonotype frequency of cytolytic CCR7^−^CX3CR1^+^ memory CD8^+^ T cells were distinct from their non-cytolytic T_CM_ (CCR7^+^CX3CR1^−^) and T_EM_ counterparts (CCR7^−^CX3CR1^−^) in blood and TDL. Fourth, the absolute numbers of T_EM_ and T_EMRA_ cells remained stable in the intravascular circulation after inhibition of S1P-dependent tissue egress via the administration of FTY-720. Cytolytic subsets of memory CD8^+^ T cells therefore appeared to be selectively retained in the intravascular space, consistent with the findings of a previous study (Gerlach et al., 2016).

The general absence of cytolytic CD8^+^ T cells in tissues hinted at the existence of an evolutionarily conserved mechanism that might operate to minimize collateral tissue damage during normal homeostasis. In this scenario, cytolytic CD8^+^ T cells would reside in the bloodstream under physiological conditions, acting as a reservoir that could be mobilized rapidly in response to infection, inflammation, or injury anywhere in the body, akin to neutrophils. Previous studies have shown that CD8^+^ T cells loaded with perforin and serine proteases infiltrate tissue sites of pathology in many autoimmune diseases, cancers, and chronic infections (Boschetti et al., 2016; Duhen et al., 2018; Nguyen et al., 2019; Reuter et al., 2017; Strioga et al., 2011). However, most of these anatomically discrete cell populations were locally induced or constitutively resident, in contrast to non-resident cytolytic CD8^+^ T cells (CD69^−^CX3CR1^+^) derived from the intravascular circulation (Buggert et al., 2018; Duhen et al., 2018). Many intracellular pathogens infect lymphocytes, other blood cells, or vascular endothelial cells (Friedman et al., 1981), as reviewed previously (Valbuena and Walker, 2006). Our data therefore suggested another possible scenario, namely a particular role for cytolytic immune surveillance in the intrasvascular compartment. In line with this concept, CMV-specific CD8^+^ T cells often circulate at high frequencies in the vasculature (Gordon et al., 2017), likely as a consequence of viral persistence in endothelial cells (Jarvis and Nelson, 2007), and typically display a cytolytic phenotype characterized by high expression levels of GzmB, perforin, and CX3CR1 (Appay et al., 2000; Buggert et al., 2014). Fractalkine, the ligand for CX3CR1, is expressed on the membrane of activated vascular endothelial cells, where it plays a critical role in leukocyte recruitment (Schulz et al., 2007), rolling (Imai et al., 1997), and adhesion (Fong et al., 1998). These collective properties could facilitate effective CD8^+^ T cell-mediated immune surveillance under the shear flow conditions that characterize the vascular environment. Of note, intravascular confinement does not necessarily preclude access to tissues. Cytolytic CD8^+^ T cells are abundant in highly perfused organs, such as bone marrow and spleen (Buggert et al., 2018; Sathaliyawala et al., 2013). Moreover, several organs, including the kidneys and liver, contain fenestrated epithelia, which enable intravascular cytolytic CD8^+^ T cells to arrest and engage target cells via cytoplasmic protrusions (Guidotti et al., 2015).

On the basis of our findings, we propose that circulating effector memory CD8^+^ T cells can be divided into two distinct subsets, namely a constitutively intravascular CCR7^−^CD27^−^CX3CR1^+^ subset with cytolytic properties and a tissue-trafficking CCR7^−^CD27^+^CX3CR1^−^ subset with stem-like properties. Importantly, we found that CD27^−^ T_EM_ and T_EMRA_ cells were clonotypically, epigenetically, functionally, and transcriptionally distinct from CD27^+^ T_EM_ and T_EMRA_ cells, which resembled T_CM_ cells in blood and TDL. Accordingly, the intravascular CCR7^−^CD27^+^CX3CR1^−^ subset likely contained the migratory CCR7^−^CD27^+^CX3CR1^−^ subset identified in TDL. Tissue-emigrant CD8^+^ T cells in the intravascular T_EM_/T_EMRA_ compartment might therefore be defined by the expression of CD27.

Intravascular HIV-specific CD8^+^ T cells in individuals with chronic progressive disease typically express a transitional memory phenotype (CCR7^−^CD27^+^CD45RA^−^), which has generally been associated with functional exhaustion and defective maturation (Buggert et al., 2014; Champagne et al., 2001). In the context of our data, this phenotype might instead define tissue-trafficking cells, which could potentially access virally infected targets in LTs and NLTs (Buggert et al., 2018; Nguyen et al., 2019; Reuter et al., 2017). We also found that functionally and phenotypically distinct virus-specific CD8^+^ T cells were clonotypically related in blood and TDL. This particular observation suggested that individual precursors in the naive CD8^+^ T cell pool differentiated into antigen-experienced memory subsets with distinct migratory properties, consistent with previous descriptions of multiple fates at the functional level (Gerlach et al., 2013; Gerlach et al., 2010).

In summary, we have provided a comprehensive atlas of the recirculating immune system in humans and identified a core signature that defines tissue-emigrant CD8^+^ T cells under homeostatic conditions. Our collective dataset might feasibly inform new developments in the field of immunotherapy, which is currently limited by several parameters, including the longevity of cell infusates and access to sites of pathology. In this light, the characterization of intravascular effector memory CD8^+^ T cells with stem-like properties and unrestricted trafficking pathways through peripheral tissues could rationalize attempts to engineer more effective cellular therapies, especially in the context of solid tumors and persistent viruses that establish latent infections, such as HIV.

## EXPERIMENTAL MODEL AND SUBJECT DETAILS

### Human samples

Matched peripheral blood and TDL samples were collected from HIV^−^ individuals undergoing thoracic duct cannulation for idiopathic or traumatic chylopericardium, chylothorax, and/or chylous ascites (n = 52; University of Pennsylvania or Children’s Hospital of Philadelphia). Matched arterial and venous blood samples were obtained from some of these patients (n = 3; University of Pennsylvania). Additional samples were collected from HIV^+^ donors with no clinical indication for thoracic duct cannulation via a research protocol (n = 10; University of Pennsylvania). Donor groups and clinical parameters are summarized in Table S1. Lymphoid and non-lymphoid organs or tissues were obtained as follows: non-enlarged tonsils from patients undergoing tonsillectomy for sleep apnea (n = 7; University of Pennsylvania), macroscopically normal mesenteric LNs from patients undergoing abdominal surgery for various indications (n = 8; Case Western Reserve University), macroscopically normal iliac LNs from kidney transplant donors (n = 6; University of Pennsylvania), liver biopsies from liver transplant donors (n = 6; Karolinska University Hospital), decidua isolated from the first trimester (n = 3; Karolinska University Hospital), and colon sections from patients undergoing abdominal surgery for various indications (n = 3; University of Pennsylvania). Venous blood samples were collected from multiple sclerosis patients before and during treatment with FTY-720 (n = 12; McGill University. Matched central and portal blood samples were obtained from liver transplant donors (n = 3; Hanover Hospital). All participants enrolled in this study provided written informed consent in accordance with protocols approved by the regional ethical research boards and the Declaration of Helsinki.

### Non-human primate samples

Rhesus macaque samples were obtained from the Oregon National Primate Research Center or the Children’s Hospital of Pennsylvania. TDL was obtained via cannulation under anesthesia, and tissues were harvested at necropsy as described previously (Buggert et al., 2018). All procedures were conducted in accordance with federal statutes and regulations, including the Animal Welfare Act.

## METHOD DETAILS

### Cells and tissues

PBMCs were purified from whole blood or leukapheresis products via standard density gradient centrifugation and cryopreserved at −140°C. Lymph node mononuclear cells were isolated via mechanical disruption and cryopreserved at −140°C. Non-lymphoid mononuclear cells were isolated using a combination of mechanical disruption and collagenase treatment. TDL was accessed as described previously (Nadolski and Itkin, 2012). Briefly, a 25-gauge spinal needle was inserted under ultrasound guidance into an inguinal LN on each side of the body, and the oil-based contrast agent ethiodol (Savage Laboratories) was injected under fluoroscopic guidance into each LN. After opacification of the cisterna chyli, access was gained via an anterior transabdominal approach using a 21-gauge or a 22-gauge Chiba needle (Cook Medical Inc.), and a V-18 control guidewire (Boston Scientific) was inserted into the thoracic duct and manipulated cephalad, followed by a 60-cm 2.3F Rapid Transit Microcatheter (Cordis Corp.), which was advanced further into the thoracic duct to aspirate TDL. Aspirated TDL was collected in heparin tubes, and the contrast agent was removed via standard density gradient centrifugation. TDL samples were used directly in flow cytometry experiments or cryopreserved at −140°C.

### Flow cytometry procedures

Cells were stained as described previously (Buggert et al., 2018). Briefly, human cryopreserved mononuclear cells were thawed and rested for at least 1 hr in complete medium (RPMI-1640 supplemented with 10% fetal bovine serum, 1% L-glutamine, and 1% penicillin/streptomycin) in the presence of 10 U/mL DNAse I (Roche). Cells were then washed in phosphate-buffered saline (PBS), prestained for chemokine receptors and adhesion molecules for 10 min at 37°C, labeled with LIVE/DEAD Fixable Aqua (Thermo Fisher Scientific) for 10 min at room temperature, and stained with an optimized panel of directly conjugated monoclonal antibodies for a further 20 min at room temperature to detect additional surface markers. In some experiments, MHC class I tetramers were added for 10 min at room temperature immediately after washing in PBS. Cells were then washed in fluorescence-activated cell sorting (FACS) buffer (PBS containing 0.1% sodium azide and 1% bovine serum albumin) and fixed/permeabilized using a Cytofix/Cytoperm Buffer Kit (BD Biosciences) or a FoxP3 Transcription Factor Buffer Kit (eBioscience). Intracellular markers were detected via the subsequent addition of an optimized panel of directly conjugated antibodies for 1 hour at 37°C. Stained cells were fixed in PBS containing 1% paraformaldehyde (Sigma-Aldrich) and stored at 4°C. Similar procedures were used to characterize activated cells stimulated for 5 hr with PMA (5 ng/mL; Sigma-Aldrich) and ionomycin (500 ng/mL; Sigma-Aldrich) and proliferating cells stimulated for 5 days with α-CD3 (1µg/mL; clone UCHT1; Bio-Rad) and α-CD28/CD49d (each at 1 µg/mL; clones L293/L25; BD Biosciences). Non-human primate cells were stained as described previously (Roberts et al., 2016). All samples were acquired within 3 days using an LSRII, an LSR Fortessa, or an LSR Symphony (BD Biosciences). Data were analyzed with FlowJo software version 9.8.8 or higher (Tree Star). The gating strategies are depicted in the relevant figures. All sorting experiments were performed using a FACSAriaII (BD Biosciences).

### Flow cytometry reagents

The following reagents were used to stain human cells: α-TCF-1 PE (clone C63D9) from Cell Signaling Technology; α-CCR5–BV650 (clone 3A9), α-CD3–AF700 (clone UCHT1), α-CD3–APC-R700 (clone UCHT1), α-CD25–PE (clone M-A251), α-CD45RA–BV650 (clone HI100), α-CD45RA–PE (clone HI100), α-CD45RA–PE-CF594 (clone HI100), α-CD45RO–BV650 (clone UCHL1), α-CD45RO–PE-CF594 (clone UCHL1), α-CD49c–BV421 (clone C3 II.1), α-CD95–BB515 (clone DX2), α-CXCR5– AF488 (clone RF8B2), α-CXCR5–AF647 (clone RF8B2), α-GzmB–Alexa700 (clone GB11), α-HLA-DR–BV605 (clone G46-6), α-HLA-DR–BV650 (clone G46-6), α-Ki67– FITC (clone B56), α-OX40–APC (clone ACT35), and α-TNF–PE-Cy7 (clone MAb11) from BD Biosciences; α-4-1BB–BV421 (clone 4B4-1), α-CCR7–APC-Cy7 (clone G043H7), α-CD3–BV711 (clone UCHT1), α-CD8a–BV570 (clone RPA-T8), α-CD8a–BV605 (clone RPA-T8), α-CD8a–BV711 (clone RPA-T8), α-CD8a–BV785 (clone RPAT8), α-CD11b–PE-Cy5 (clone ICRF44), α-CD14–BV510 (clone M5E2), α-CD18–PE-Cy7 (clone 1B4), α-CD19–BV510 (clone HIB19), α-CD27–AF700 (clone O323), α-CD27–BV650 (clone O323), α-CD27–BV785 (clone O323), α-CD29–AF700 (clone TS2/16), α-CD38–APC (clone HIT2), α-CD38–BV711 (clone HIT2), α-CD49a–PE-Cy7 (clone TS2/7), α-CD49b–APC (clone P1E6-C5), α-CD49f–BV650 (clone GoH3), α-CD52–FITC (clone HI186), α-CD69–BV421 (clone FN50), α-CD69–PE-Cy5 (clone FN50), α-CD103–BV605 (clone Ber-ACT8), α-CD103–PE-Cy7 (clone Ber-ACT8), α-CX3CR1–PE (clone 2A9-1), α-CX3CR1–APC (clone 2A9-1), α-CXCR2–FITC (clone 5E8), α-CXCR3–BV711 (clone G025H7), α-CXCR6–BV421 (clone K041E5), α-PD-1– BV421 (clone EH12.2H7), α-PD-1–PE (clone EH12.2H7), α-PD-L1–BV785 (clone 10F.9G2), α-perforin–BV421 (clone B-D48), α-perforin–PE-Cy7 (clone B-D48), and α-T-bet–PE-Dazzle 594 (clone 4B10) from BioLegend; α-Eomes–AF647 (clone WD1928), α-ICOS–PE-EF610 (clone ISA-3), α-perforin–PE-Cy7 (clone dG9), and a20 T-bet–PE (clone 4B10) from eBioscience; α-a4b7–FITC (clone Act-1) from the NIH AIDS Reagent Program; and α-CD4–PE-Cy5.5 (clone S3.5) and α-IFNγ–AF700 (clone B27) from Thermo Fisher Scientific. Dead cells were identified using LIVE/DEAD Fixable Aqua (Thermo Fisher Scientific), and proliferating cells were identified using CTV (Thermo Fisher Scientific).

The following reagents were used to stain non-human primate cells: α-CD3–APC-Cy7 (clone SP34-2), α-CD95–PE-Cy5 (clone DX2), and α-GzmB–AF700 (clone GB11) from BD Biosciences; α-CD28–ECD (clone CD28.2) from Beckman Coulter; α-CD8a– BV570 (clone RPA-T8), α-CD14–BV650 (clone M5E2), α-CD16–BV650 (clone 3G8), and α-CD20–BV650 (clone 2H7) from BioLegend; α-T-bet–PE-Cy7 (clone 4B10) from eBioscience; α-perforin–FITC (clone pf344) from Mabtech AB; and α-CD4–PE-Cy5.5 (clone S3.5) from Thermo Fisher Scientific. Dead cells were identified using LIVE/DEAD Fixable Aqua (Thermo Fisher Scientific).

### Tetramers

MHC class I tetramers conjugated to BV421 or PE were used to detect CD8^+^ T cells with the following specificities: CMV NLVPMVATV (NV9/HLA-A*0201), HIV FLGKIWPSHK (FK10/HLA-A*0201), HIV ILKEPVHGV (IV9/HLA-A*0201), HIV SLYNTVATL (SL9/HLA-A*0201), HIV KYKLKHIVW (KW9/HLA-A*2402), HIV RYPLTFGW (RW8/HLA-A*2402), HIV GPGHKARVL (GL9/HLA-B*0702), HIV HPRVSSEVHI (HI10/HLA-B*0702), HIV SPAIFQSSF (SM9/HLA-B*0702), HIV KRWIILGLNK (KK10/HLA-B*2705), HIV ISPRTLNAW (IW9/B*5701), HIV KAFSPEVIPMF (KF11/HLA-B*5701), and HIV QASQEVKNW (QW9/HLA-B*5701),and HIV TSTLQEQIGW (TW10/HLA-B*5701). All tetramers were generated in house as described previously (Price et al., 2005).

### RNA-seq

RNA-seq was conducted as described previously (Buggert et al., 2018; Vella et al., 2019). Briefly, naive and memory CD8^+^ T cells from mesenteric LNs, blood, and TDL (250 cells per subset) were sorted directly into lysis buffer (Takara) using a FACSAriaII (BD Biosciences) and frozen at −140°C. Libraries were prepared using the SMART-Seq v4 Ultra Low Input RNA Kit (Takara). PCR products were then indexed using a Nextera XT DNA Library Prep Kit (Illumina) and sequenced across 75 base pairs (bp) using a paired-end strategy with a 150-cycle high-output flow cell on a NextSeq 550 (Illumina). Three biological replicates were sequenced per experiment. Fastq files from replicate sequencing runs were concatenated and aligned to hg38 using STAR software version 2.5.2a. Mapped read depth ranged from 8 million to 75 million reads per sample. Aligned files were normalized using DESeq2 (Bioconductor).

Differentially expressed genes were identified among normalized RNA-seq counts from distinct subsets of CD8^+^ T cells using a t-test (p < 0.05) in the R limma package (version 3.28). Genes of interest were then subjected to Ingenuity Pathway Analysis (Qiagen Bioinformatics). Heatmaps, PCA plots, and tSNE plots were created using the Pheatmap (version 1.0.12) and ggplot2 (version 3.1.0) packages in R or Prism version7.0 (GraphPad). Normalized counts were also subjected to GSEA using software from the Broad Institute (http://www.broadinstitute.org/gsea/index.jsp). Gene signatures were compared with immunological and GO signatures from the Broad Institute Database.

### Redirected killing assay

P815 mastocytoma target cells were labeled with LIVE/DEAD Fixable Violet (Thermo Fisher Scientific) and TFL4 (OncoImmun), washed twice in PBS, and incubated for 30 min at room temperature with α-CD3 (1 µg/mL; clone UCHT1; Bio-Rad). CD8^+^ T cells were negatively selected from blood and TDL using a CD8^+^ T Cell Enrichment Kit (stemCell Technologies). Isolated CD8^+^ T cells were rested in complete medium for at least 45 min at 37°C and then incubated with α-CD3-coated P815 cells at different E:T ratios in a 96-well V-bottom plate for 4 hr at 37°C. Cells were then stained with α-active caspase-3–FITC (clone C92-605; BD Biosciences) and α-CD3–PE (clone SK7; BD Biosciences) and acquired using an LSRII (BD Biosciences). Cytolytic activity was calculated by subtracting the frequency of active caspase-3^+^TFL4^+^LIVE/DEAD^−^ P815 cells in target-only wells from the frequency of active caspase-3^+^TFL4^+^LIVE/DEAD^−^ P815 cells in wells containing CD8^+^ T cells.

### ATAC-seq

ATAC-seq was performed as described previously (Buenrostro et al., 2013) with minor modifications (Buggert et al., 2018). Briefly, memory CD8^+^ T cells from blood, mesenteric LNs, and TDL (30,000–60,000 cells per subset) were sorted directly into complete medium using a FACSAriaII (BD Biosciences). Cells were then pelleted, washed with PBS, and treated with lysis buffer (10 mM Tris-HCl pH 7.4, 10 mM NaCl,3 mM MgCl_2_, 0.1% IGEPAL CA-630). Nuclear pellets were resuspended in the presence of Tn5 transposase (Illumina) for 30 minutes at 37°C. Tagmented DNA was purified using a MinElute Reaction Cleanup Kit (Qiagen). Amplified libraries were purified using a QIAQuick PCR Purification Kit (Qiagen) and sequenced across 75 or 100 bp using a paired-end strategy on a NextSeq 550 or a HiSeq4000 (Illumina). One or two biological replicates were sequenced per experiment.

The data processing pipeline is available at GitHub (https://github.com/wherrylab/jogiles_ATAC/blob/master/Giles_Wherry_ATAC_pipeline_hg19_UPennCluster.sh). Fastq files were aligned to hg19 using Bowtie2 (http://www.bowtie-bio.sourceforge.net/bowtie2). Unmapped, unpaired, and mitochondrial reads were removed using SAMtools (http://www.htslib.org). Duplicates were removed using Picard (https://broadinstitute.github.io/picard), and blacklist regions were removed using bedtools subtract (https://bedtools.readthedocs.io). ATAC-seq peaks were called at a false discovery rate of 0.01 using MACS2 (https://pypi.python.org/pypi/MACS2). Normalization and differential peak analyses were performed using DESeq2 (Bioconductor) as described previously (Vella et al., 2019). Motif analyses were carried out using HOMER (http://homer.ucsd.edu/homer). ATAC-seq tracks were visualized using Integrative Genomics Viewer software version2.5.2 (https://software.broadinstitute.org/software/igv).

### TCR-seq

Expressed *TRB* gene rearrangements were sequenced as described previously (Boritz et al., 2016). Briefly, bulk memory or virus-specific CD8^+^ T cells were sorted into cold fetal bovine serum and frozen in RNAzol (Molecular Research Center). TCRβ transcripts were amplified using a template-switch anchored RT-PCR. Libraries were sequenced across 150 bp using a paired-end strategy on a NextSeq 550 (Illumina). TCRβ sequences were annotated using a custom Java script and BLAST (National Center for Biotechnology Information). CDR3βsequences were assigned between the conserved cysteine at the 3’ end of each *TRBV* gene and the conserved phenylalanine at the 5’ end of each *TRBJ* gene. Unique TCRβcombinations were collapsed to determine clonotype counts.

## QUANTIFICATION AND STATISTICAL ANALYSIS

Unmatched groups were compared using an unpaired t-test or the Mann-Whitney U test, and matched groups were compared using a paired t-test or the Wilcoxon signed-rank test. Non-parametric tests were used if the data were not distributed normally according to the Shapiro-Wilk test. Correlations were assessed using the Pearson correlation or the Spearman rank correlation. All analyses were performed using RStudio (https://rstudio.com) or Prism version 7.0 (GraphPad). Functional profiles were compared using the permutation test in SPICE version 6.0 (https://niaid.github.io/spice). Phenotypic relationships within multivariate datasets were visualized using FlowJo software version 10.6.1 (Tree Star).

## DATA AND SOFTWARE AVAILABILITY

The sequence datasets reported in this paper have been deposited in the Gene Expression Omnibus under accession no. XXX.

## ACKNOWLEDGEMENTS

We express our gratitude to all study participants and funding agencies.

## FUNDING

M.B. was supported by the Swedish Research Council, the Karolinska Institutet, the Swedish Society for Medical Research, the Jeansson Stiftelse, the Åke Wibergs Stiftelse, the Swedish Society of Medicine, the Cancerfonden, the Barncancerfonden, the Magnus Bergvalls Stiftelse, the Lars Hiertas Stiftelse, the Swedish Physician against AIDS Foundation, the Jonas Söderquist Stiftelse, and the Clas Groschinskys Minnesfond. Additional support was provided by R01 and R56 grants from the National Institutes of Health (AI076066, AI118694, and AI106481 to M.R.B.) and the Penn Center for AIDS Research (AI045008). D.A.P. is a Welcome Trust Senior Investigator (100326/Z/12/Z).

## AUTHOR CONTRIBUTIONS

M.B. and M.R.B. conceived the project; M.B. L.A.V., S.N., T.S., A.P.P., C.M., A.R., L.K.C., A.S.J., I.B.B., M.I., and L.H. designed and performed experiments; M.B. L.A.V., S.N., V.W., T.S., A.P.P., S.M., S.D., J.P.A., M.E.J., and G.V. analyzed data; M.B., A.O., L.P., E.S., S.L.L., E.G., D.A.P., N.B., A.B.O., Y.D., A.N., D.H.C., I.F., D.C.D., E.J.W., M.G.I., and M.R.B. provided critical resources; M.B., D.A.P., N.B., D.H.C., T.M.L., A.D.W., D.C.D., E.J.W., and M.R.B. supervised experiments; M.B. and M.R.B. drafted the manuscript; M.B., D.A.P., and M.R.B. edited the manuscript.

## COMPETING INTERESTS

The authors declare that they have no competing financial interests, patents, patent applications, or material transfer agreements associated with this study.

## Notes

### Competing Interest Statement

The authors have declared no competing interest.

