## Supplemental information for "The identity of human tissue-emigrant CD8^+^ T cells"

Figure S1

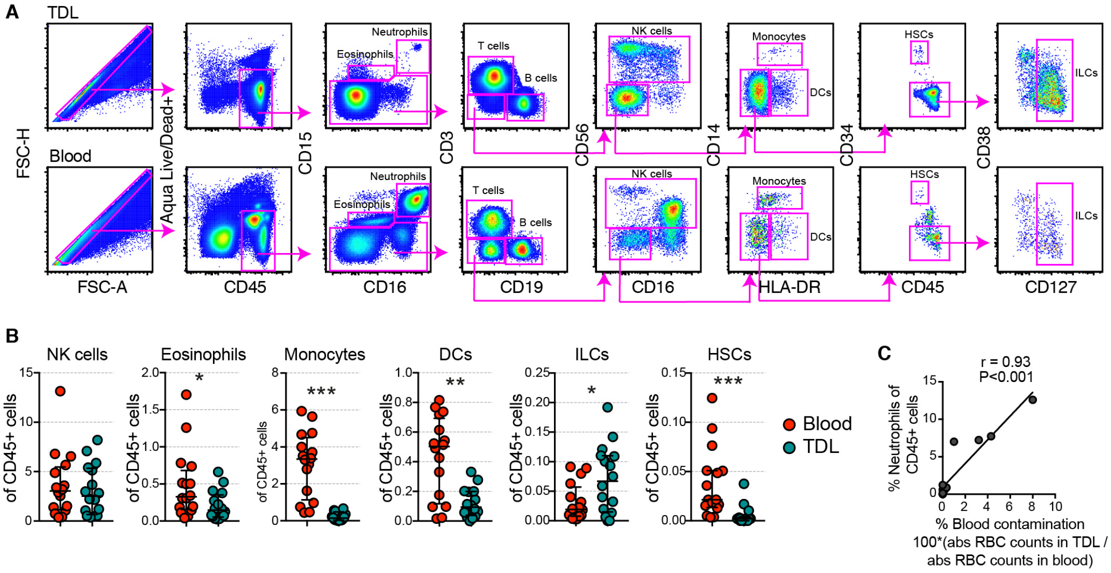

**Figure S1. Flow cytometric analysis of immune cell populations in matched samples of blood and TDL.** (**A**) Flow cytometric gating strategy used to identify immune lineage subsets in blood and TDL. (**B**) Quantification of immune subsets in blood versus TDL using a value-adapted y-axis. (**C**) Correlation between the frequency of neutrophils and the relative frequency of red blood cells (RBCs) in TDL. The absolute count in TDL divided by the absolute count in blood was used to calculate the relative frequency of RBCs. *p < 0.05, **p < 0.01, ***p < 0.001. Related to Figure 1.

Figure S2

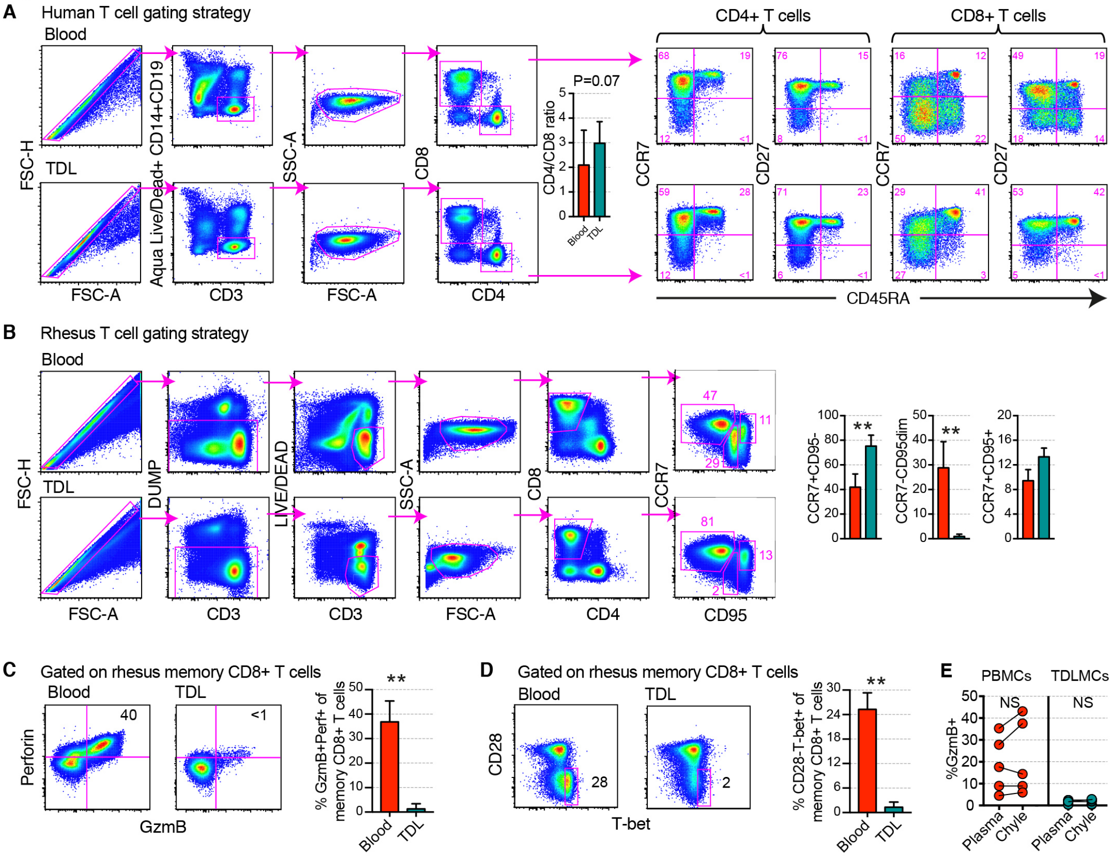

**Figure S2. Flow cytometric analysis of human and rhesus macaque naive and memory CD8$^+$ T cell subsets in blood and TDL.** (**A**) Flow cytometric gating strategy for the detection of human CD4$^+$ and CD8$^+$ T cell subsets in blood and TDL. The bar graph shows the overall CD4 versus CD8 ratio among CD3$^+$ cells in blood and TDL (n = 16). (**B**) Flow cytometric gating strategy for the detection of rhesus macaque CD8$^+$ T cell subsets in blood and TDL. The bar graphs show the overall distribution of naive and memory CD8$^+$ T cell subsets based on the expression of CCR7 and CD95 (n = 8). (**C**) Flow cytometric quantification of granzyme B and perforin among rhesus macaque memory CD8$^+$ T cells (non-CD28$^+$CD95$^-$) subsets in blood and TDL (n = 8). (**D**) Flow cytometric quantification of T-bet among rhesus macaque memory CD8$^+$ T cells (non-CD28$^+$CD95$^-$) subsets in blood and TDL (n = 8). **(E)** Flow cytometric quantification of granzyme B among PBMCs and TDLMCs before and after incubation with matched plasma or chyle (TDL) for 5 days. **$p < 0.01$. Related to Figures 2, 3, and 4.

Figure S3

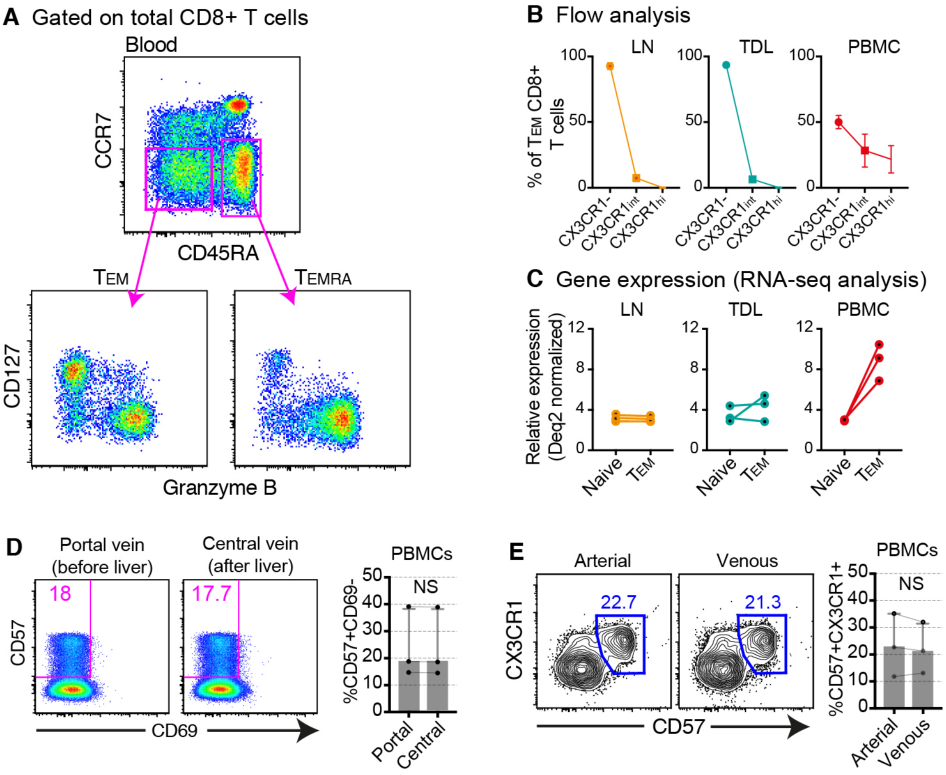

**Figure S3. Flow cytometric and transcriptomic analyses of effector memory $CD8^+$ T cell subsets in blood, TDL, and mesenteric LNs.** (**A**) Representative flow cytometry plots showing the expression pattern of CD127 versus granzyme B among intravascular $CD8^+$ $T_{EM}$ and $T_{EMRA}$ cells. (**B**) Flow cytometric analysis showing percent expression of CX3CR1 among $CD8^+$ $T_{EM}$ cells in blood, TDL, and mesenteric LNs. (**C**) RNA-seq analysis showing relative expression of CX3CR1 among $CD8^+$ $T_{EM}$ cells in blood, TDL, and mesenteric LNs. (**D**) Representative flow cytometry plots (left) and summary graphs (right) showing the frequencies of non-resident cytolytic $CD8^+$ T cells ($CD57^+CD69^-$) in the portal and central hepatic veins (n = 5). (**E**) Representative flow cytometry plots (left) and summary graphs (right) showing the frequencies of non-resident cytolytic $CD8^+$ T cells ($CD57^+CD69^-$) in arterial and venous blood (n = 5). Related to Figure 5.

Figure S4

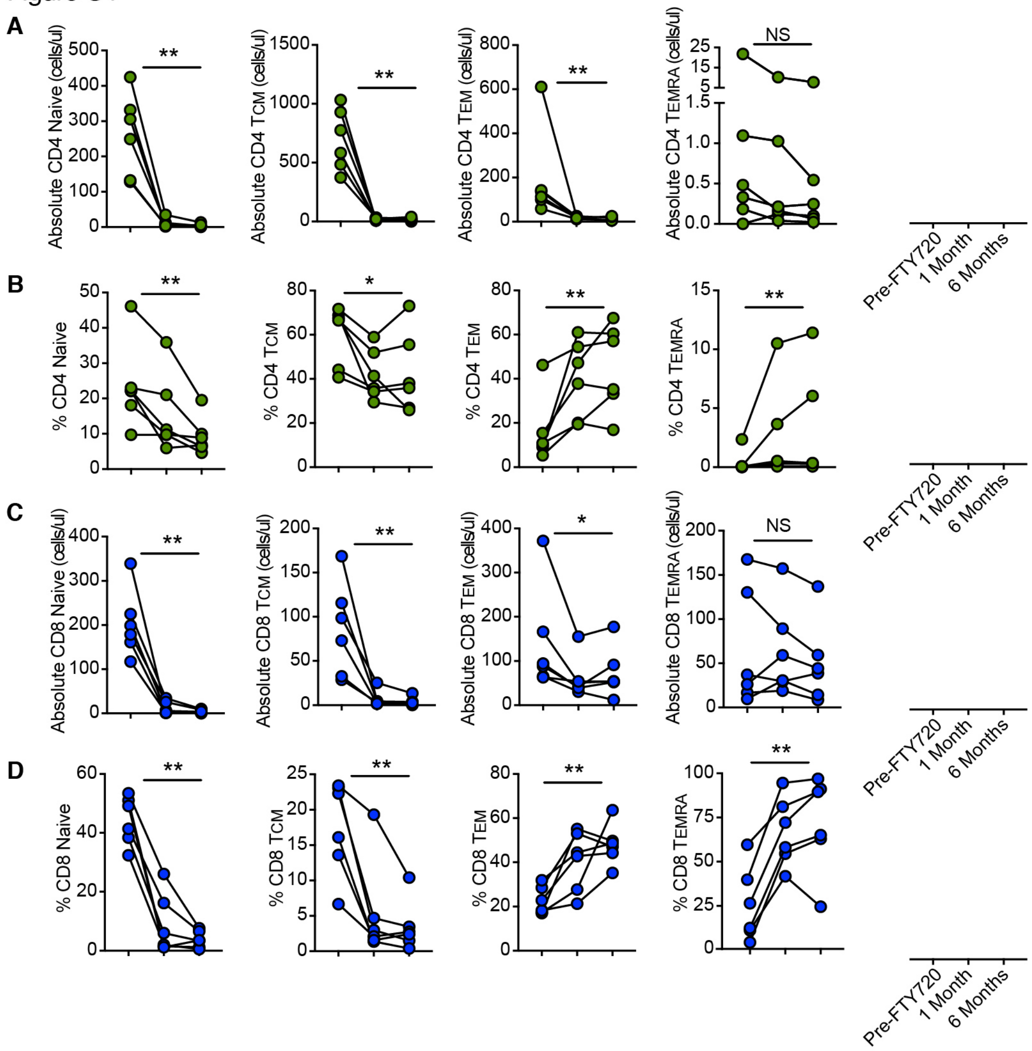

**Figure S4. Global effects of FTY-720 on the intravascular T cell compartment.** Before-after-graphs showing the impact of FTY-720 on naive (left), $T_{CM}$ (middle left), $T_{EM}$ (middle right) and $T_{EMRA}$ cells (right) in the $CD4^+$ and $CD8^+$ lineages. (**A**) Absolute numbers of naive and memory $CD4^+$ T cells. (**B**) Frequencies of naive and memory $CD4^+$ T cells. (**C**) Absolute numbers of naive and memory $CD8^+$ T cells. (**D**) Frequencies of naive and memory $CD8^+$ T cells. *p < 0.05, **p < 0.01. Related to Figure 6.

Figure S5

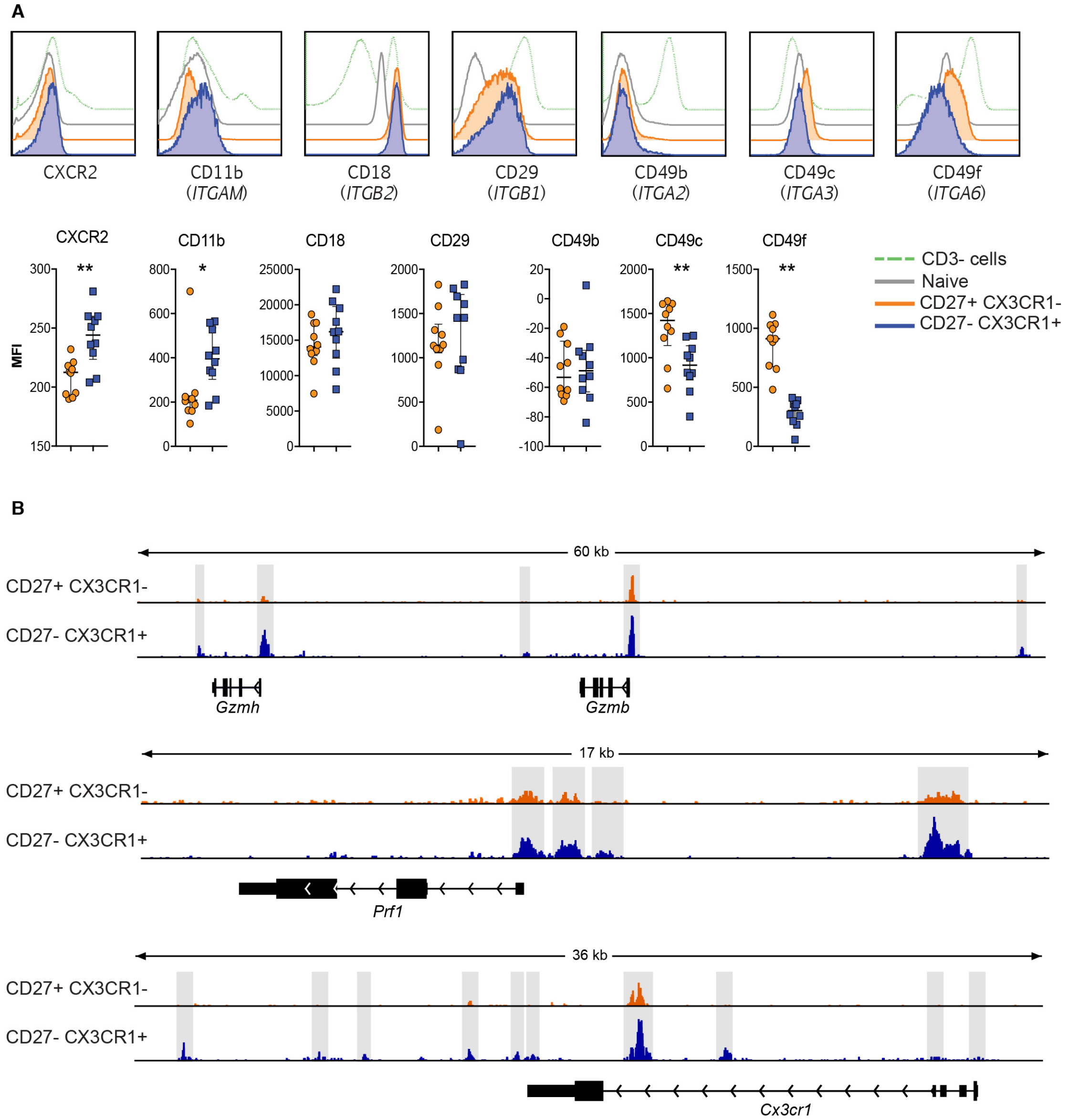

**Figure S5. Flow cytometric and epigenetic analyses of intravascular cytolytic and non-cytolytic effector memory CD8⁺ T cells.** (**A**) Representative flow cytometry histograms (top) and summary graphs (bottom) showing the expression levels of chemokine receptors and integrins that were differentially expressed among CCR7$^-$CD27$^+$CX3CR1$^-$ and CCR7$^-$CD27$^-$CX3CR1$^+$ memory CD8$^+$ T cells in the corresponding RNA-seq experiments. MFI: median fluorescence intensity. (**B**) ATAC-seq tracks for *Gzmh*, *Gzmb*, *Prf1*, and *Cx3cr1* comparing CCR7$^-$CD27$^+$CX3CR1$^-$ versus CCR7$^-$CD27$^-$CX3CR1$^+$ memory CD8$^+$ T cells. *p < 0.05, **p < 0.01. Related to Figure 7.

Figure S6

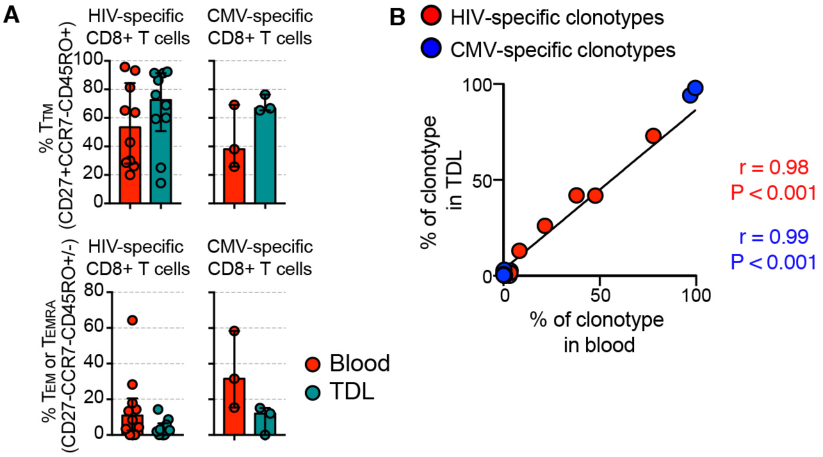

**Figure S6. Anatomical distribution of virus-specific CD8$^+$ T cells and clonotypes in blood versus TDL.** (**A**) MHC class I tetramer-based quantification of virus-specific CD8$^+$ $T_{TM}$ (top) and $T_{EM}$/$T_{EMRA}$ cells (bottom) in blood and TDL. (**B**) Correlation between the frequency of virus-specific CD8$^+$ T cell clonotypes in blood and the frequency of virus-specific CD8$^+$ T cell clonotypes in TDL. Each dot represents a matched specificity with a minimum of 200 sorted cells per compartment. Related to Figure 8.

**Table S1.** Donor groups and clinical parameters.

| HIV status | Cohort size | Samples | Age (years) | Sex | Clinical details |
|---|---|---|---|---|---|
| HIV$^-$ | 52 | Matched blood and TDL | Median: 31y<br>IQR: 10–61y | Male:<br>n = 31<br><br>Female:<br>n = 21 | Idiopathic or traumatic chylopericardium, chylothorax, and/or chylous ascites |
| HIV$^+$ | 11 | Matched blood and TDL | Median: 39y<br>IQR: 28–47y | Male:<br>n = 11<br><br>Female:<br>n = 0 | ART+ ART-<br>n = 8 n = 3<br>VL: <50 73,430<br>CD4: 936 416 |

**Table S2. List of top 500 differently expressed genes btw Tem blood and TDL**

| Gene | logFC | P.Value |
|---|---|---|
| SLC17A2 | -0.0977409 | 5.80E-07 |
| 403312 | 0.18825617 | 8.75E-06 |
| IFI6 | 3.28566446 | 9.82E-06 |
| SORCS3 | -0.0779164 | 2.28E-05 |
| NRG1 | -0.1070396 | 3.06E-05 |
| COQ3 | -1.7933505 | 3.98E-05 |
| REG4 | -3.9658268 | 4.59E-05 |
| BST2 | 1.3521969 | 4.88E-05 |
| SCUBE1 | -0.519561 | 6.04E-05 |
| PJA2 | -1.1383138 | 7.00E-05 |
| LINC01058 | 0.34336625 | 7.14E-05 |
| FCRLA | -0.0237408 | 9.12E-05 |
| IFI44L | 7.26977997 | 9.23E-05 |
| NSL1 | 0.94158534 | 0.0001093 |
| IFI44 | 2.78787504 | 0.00011652 |
| DRAP1 | 0.80594148 | 0.00012421 |
| BATF | 1.41853505 | 0.00012551 |
| 100506241 | 0.10102024 | 0.00013017 |
| IFITM1 | 2.19222006 | 0.00014681 |
| IFIT1 | 6.49614544 | 0.00014777 |
| 101928539 | -0.5299032 | 0.00017292 |
| ODF3B | 1.27362521 | 0.00017385 |
| CERS5 | -1.2418634 | 0.00017843 |
| APOF | -0.3882323 | 0.00017919 |
| CX3CR1 | 4.53378031 | 0.00017946 |
| MAX | -1.1080599 | 0.00017949 |
| ISG20 | 1.54598556 | 0.000187 |
| URAHP | -0.0343804 | 0.00019802 |
| MIR646HG | 0.28494857 | 0.00022416 |
| MTRNR2L10 | 0.08115269 | 0.00023007 |
| PCDH11Y | -0.0483361 | 0.0002576 |
| CEP68 | -2.3554273 | 0.00028107 |
| IL4I1 | 1.28255746 | 0.0002924 |
| MAT2A | -1.7322751 | 0.00033288 |
| NMU | -0.4421263 | 0.00035108 |
| C8G | -0.3017391 | 0.00036629 |
| PDCD6 | 0.80371794 | 0.0004139 |
| ISG15 | 2.28540109 | 0.0004375 |
| TRQ-CTG1-5 | -0.073734 | 0.0004459 |
| RBM24 | 0.47149744 | 0.00044819 |
| TMEM14B | 0.71316035 | 0.00048251 |
| CARS2 | -1.288773 | 0.00048257 |
| PLGRKT | 0.96918608 | 0.00050282 |
| NRIR | 1.42474903 | 0.00054467 |
| ADAMTSL4-A | 0.58841298 | 0.00056809 |
| TBC1D4 | -1.9750488 | 0.00057062 |

| | | |
|---|---|---|
| CNNM4 | -1.4263525 | 0.00058088 |
| LPAL2 | -0.4080653 | 0.00059591 |
| RHOBTB2 | 1.42702864 | 0.00063534 |
| PARP3 | -2.9750653 | 0.0006402 |
| NOD2 | 1.66626754 | 0.00069794 |
| SMIM23 | -0.1951544 | 0.00072964 |
| TMED6 | 0.72670071 | 0.0007366 |
| C10orf35 | -2.0365701 | 0.00074084 |
| VSX2 | 0.06507291 | 0.00076758 |
| PNPLA2 | -1.5303518 | 0.00078337 |
| LINC01448 | -0.0709116 | 0.00082583 |
| LINC00094 | -2.7967386 | 0.00084859 |
| MYSM1 | 1.36981113 | 0.00096363 |
| NUP93 | -1.4212354 | 0.00098367 |
| STMN2 | 0.11988287 | 0.00099648 |
| KIAA0232 | -1.3292924 | 0.00099827 |
| DLAT | -2.1698955 | 0.00102516 |
| EGFL6 | -0.3364684 | 0.00102798 |
| LSG1 | -1.348099 | 0.00105786 |
| EIF3B | -1.4620063 | 0.00106122 |
| LSP1 | -0.9093712 | 0.001072 |
| FMO3 | -0.1445572 | 0.00110698 |
| RNU1-1 | 0.333678 | 0.00111498 |
| UNK | -1.3196099 | 0.00114134 |
| RECQL | -1.3132853 | 0.00115728 |
| TPD52L3 | -0.2205678 | 0.00120447 |
| PYCR2 | -1.5282526 | 0.00121117 |
| LINC00824 | -0.299325 | 0.0012138 |
| CCNL2 | -1.2873939 | 0.00123782 |
| GZMH | 3.20713829 | 0.00132738 |
| HIC1 | -0.4265331 | 0.0013311 |
| SLC28A1 | -0.1060302 | 0.00133246 |
| TUBG1 | -2.2643507 | 0.00133624 |
| TMEM30A | -2.4517189 | 0.00138593 |
| INMT | -0.091294 | 0.00139162 |
| DGKA | -1.1875156 | 0.00139636 |
| TRBV20-1 | 2.44748561 | 0.00141008 |
| TOB2P1 | 0.0693015 | 0.00142147 |
| C7orf73 | 0.5298448 | 0.00142453 |
| CMPK2 | 3.64878753 | 0.00143617 |
| MX1 | 2.13527388 | 0.00151534 |
| 101928438 | 0.3918616 | 0.00154874 |
| HELT | -0.1277908 | 0.00158462 |
| ARHGAP5 | -1.6581942 | 0.00160987 |
| CYSLTR1 | -4.2162129 | 0.00164581 |
| PA2G4P4 | -0.6613471 | 0.00164916 |
| WDR19 | -1.880125 | 0.00165644 |
| 105374314 | -0.2668137 | 0.00166149 |

| | | |
|---|---|---|
| LINC01559 | 0.04024479 | 0.00166989 |
| STXBP2 | -1.3469591 | 0.00169977 |
| GTF2IP4 | -0.3667406 | 0.00173099 |
| RNF146 | 1.73527121 | 0.00174264 |
| SCYL1 | -2.4468571 | 0.00174421 |
| MT-ATP6 | 0.87134369 | 0.00181784 |
| NIFK-AS1 | 1.11139987 | 0.00186635 |
| NUCB1 | -1.0606674 | 0.00188933 |
| CEP95 | -1.2702139 | 0.00189228 |
| AES | -1.1851222 | 0.00199341 |
| SARS2 | -2.1067308 | 0.00202503 |
| 100506023 | 1.42663167 | 0.00203759 |
| IKBKG | -1.077878 | 0.00205666 |
| SCARNA2 | 1.18306397 | 0.00211082 |
| CEP162 | -1.4116782 | 0.00217399 |
| LINC00853 | 0.28014414 | 0.0021834 |
| LDHA | 0.78336807 | 0.00219564 |
| GLIPR2 | 1.0962834 | 0.00219612 |
| NCF1C | 1.2732488 | 0.00220052 |
| ZNF814 | 1.34204463 | 0.00222649 |
| RRAGB | -1.0773512 | 0.00224111 |
| LRP4-AS1 | -0.0896781 | 0.00230006 |
| TBCB | 0.72003068 | 0.00232055 |
| SLC35F4 | -0.209417 | 0.00236497 |
| PRRX2 | 0.42727738 | 0.00246533 |
| 101927417 | -0.1424947 | 0.00246991 |
| NOMO2 | -0.8090528 | 0.00251809 |
| MCCC2 | -1.3682604 | 0.00252335 |
| PKNOX2 | -0.0554589 | 0.00256722 |
| PLEKHB1 | -3.2606703 | 0.0026766 |
| NARS | -0.8291303 | 0.00274375 |
| PSMA7 | 0.71042788 | 0.00279385 |
| TRBV13 | 0.53507009 | 0.00287249 |
| 101927830 | 0.12690291 | 0.00287392 |
| ACSL6 | -2.5424423 | 0.00289916 |
| MOGAT2 | -0.0258091 | 0.00299753 |
| MIER2 | -1.0800369 | 0.00303325 |
| CYBB | 0.04709992 | 0.00304065 |
| SERPINE1 | 0.03453183 | 0.00304227 |
| LRIT2 | 0.04712649 | 0.00307289 |
| NANS | -1.030751 | 0.00309719 |
| HSPA4 | -1.3214742 | 0.0031487 |
| BTBD1 | -1.2769433 | 0.00320113 |
| RPL31P11 | 0.24030359 | 0.0032016 |
| RSRC1 | 1.04865009 | 0.00326668 |
| PITHD1 | -0.7862437 | 0.00327806 |
| ZBED5 | -1.1461964 | 0.00328823 |
| CTHRC1 | 0.20805025 | 0.0032932 |

| | | |
|---|---|---|
| 100507195 | 1.03550245 | 0.00329724 |
| ACACA | -0.2435005 | 0.00330472 |
| CLIP3 | -0.9645448 | 0.00331235 |
| PKM | -1.2969968 | 0.00332428 |
| ADD1 | -0.8231125 | 0.00334554 |
| RECK | -1.8828973 | 0.00338395 |
| PSME2 | 1.02486023 | 0.00338508 |
| CLDN12 | -1.1446221 | 0.00340033 |
| MEIG1 | 0.52684333 | 0.00342952 |
| NFE2L2 | -1.0799043 | 0.00343 |
| ZNF385B | 0.14314927 | 0.00343474 |
| PLSCR1 | 1.63010397 | 0.00343875 |
| SNRPA | -0.6565384 | 0.00346514 |
| CERS4 | -1.5358045 | 0.00353899 |
| RNF213 | 1.2983045 | 0.00360035 |
| SNORD70 | -0.092562 | 0.00361486 |
| CDC42 | 1.0607753 | 0.00361544 |
| ARNT | -1.6351588 | 0.00362645 |
| RBM27 | 1.24859602 | 0.00362736 |
| WDR1 | -0.8867269 | 0.00363064 |
| ZNF248 | -1.4213294 | 0.00363649 |
| CLEC12B | 0.04491896 | 0.0036755 |
| CCR4 | -2.4466703 | 0.00374201 |
| 101929524 | -0.5141638 | 0.00378007 |
| SCNN1G | -0.0409246 | 0.00383528 |
| UBR3 | -1.4786761 | 0.00383538 |
| 100996671 | 0.16764858 | 0.00383572 |
| NFX1 | -1.4126333 | 0.00384952 |
| ACADL | -0.0243979 | 0.00385385 |
| NEK9 | 1.11591594 | 0.0039048 |
| GDI1 | -1.3271555 | 0.00395723 |
| MIR548AC | 0.06061799 | 0.00396302 |
| DCAF15 | -0.9932269 | 0.00402516 |
| MYLIP | -1.4908165 | 0.00404655 |
| DENND5A | 1.47879265 | 0.00407142 |
| HSPBP1 | -0.8925282 | 0.00408793 |
| PFKL | -1.3263348 | 0.00412282 |
| RP2 | -1.4220986 | 0.00412359 |
| KLHL26 | -1.3761592 | 0.00416823 |
| WBP2 | -0.97926 | 0.00423507 |
| DGUOK-AS1 | 0.34654774 | 0.00424263 |
| LINC00836 | -0.1247522 | 0.00430485 |
| TRDJ3 | -0.393315 | 0.00435744 |
| YEATS4 | 0.83761199 | 0.00440718 |
| LINC00113 | 0.03004382 | 0.004434 |
| PTPRM | 2.68522435 | 0.00445097 |
| ACYP2 | 1.04484192 | 0.00446533 |
| HEMGN | -1.6032187 | 0.00452408 |

| | | |
|---|---|---|
| LSM14B | 1.03329167 | 0.00453593 |
| MRPL28 | 0.58896984 | 0.00454291 |
| MCMDC2 | 0.89898708 | 0.0046524 |
| HCFC1R1 | -1.0092417 | 0.00465395 |
| TMEM259 | -1.5706619 | 0.00469788 |
| SIRPB1 | 0.22663432 | 0.00474983 |
| EFCAB12 | -0.2562591 | 0.00476407 |
| ENAH | -0.3902452 | 0.00478866 |
| MT-CO3 | 0.86187039 | 0.00482128 |
| CLCN7 | -2.1085376 | 0.00484097 |
| CCDC93 | -1.294488 | 0.00486707 |
| 100131626 | -0.1026925 | 0.00500433 |
| ZGRF1 | -2.2130626 | 0.00502719 |
| DCXR | 0.82167205 | 0.00506643 |
| MIR155HG | 1.60538108 | 0.00509198 |
| ADNP-AS1 | -1.2965586 | 0.00510386 |
| DUT | 0.66174312 | 0.00510756 |
| ASIP | -0.2420152 | 0.00512964 |
| PSME3 | -0.7861312 | 0.00513984 |
| MADD | -1.3893045 | 0.00514709 |
| MINA | -1.4702493 | 0.00518811 |
| 101927243 | -0.2306738 | 0.00523701 |
| PIM1 | 1.14102525 | 0.00524321 |
| TBK1 | -0.9460642 | 0.00524777 |
| ENTHD1 | -0.4918665 | 0.00525095 |
| CBY1 | -1.6992088 | 0.00525638 |
| LRRC8D | -2.2414783 | 0.00526487 |
| UBE2L6 | 0.9731189 | 0.00530073 |
| SCOC-AS1 | 0.82343686 | 0.00533641 |
| CAP1 | -0.7574189 | 0.00536109 |
| FBXW11 | 1.41279372 | 0.00536352 |
| TTC37 | -0.9689577 | 0.00540922 |
| 101928834 | 0.07708721 | 0.00544085 |
| PRICKLE4 | -0.6397907 | 0.00544139 |
| PTMA | 0.59749198 | 0.00544197 |
| EPN1 | -0.9265937 | 0.00547557 |
| ADAM20 | -0.8201617 | 0.00547792 |
| EXOC6 | -1.4372062 | 0.00552083 |
| FDPS | 0.69231202 | 0.00557714 |
| RWDD2A | -2.2369232 | 0.00559158 |
| LINC00673 | -0.2859936 | 0.00561407 |
| SPON2 | 0.90711565 | 0.00561536 |
| PDE6D | 0.81544657 | 0.00563937 |
| RNU2-2P | 1.13623192 | 0.00564016 |
| EPSTI1 | 2.41565043 | 0.00564106 |
| C5orf56 | 1.5007057 | 0.00566305 |
| ANXA6 | -1.4752542 | 0.00569946 |
| KMT2E | 0.68295316 | 0.00576857 |

| | | |
|---|---|---|
| SP6 | -0.0339893 | 0.00579069 |
| S1PR4 | -0.5887017 | 0.00586096 |
| PPEF1 | -0.3551045 | 0.00587825 |
| TNKS2 | -1.1420174 | 0.00588577 |
| PHGDH | -2.997042 | 0.00590373 |
| SYN1 | -0.1714615 | 0.00592118 |
| PPP2CB | -2.0209992 | 0.00592994 |
| BMS1 | -1.0162101 | 0.00598056 |
| RSAD2 | 3.64159332 | 0.00598119 |
| ZNF441 | -2.4648005 | 0.00600467 |
| MED28 | 0.77997947 | 0.00601741 |
| GAREM1 | -0.0564024 | 0.00606321 |
| MYH7B | 0.83574724 | 0.00613933 |
| BIN2 | -0.8577895 | 0.00614764 |
| ATP8B2 | -1.3375321 | 0.0061629 |
| TMEM106B | -1.0051293 | 0.0061923 |
| ZNF625-ZNF | 0.11578284 | 0.00622975 |
| SP100 | 0.6312428 | 0.00625143 |
| SLFN5 | -1.3287092 | 0.0062568 |
| ATL1 | -0.9678199 | 0.00628559 |
| TMEM256-PI | -0.1912066 | 0.00635537 |
| RNF20 | -1.4572504 | 0.00638673 |
| CAMTA1 | 0.78104931 | 0.00640915 |
| FEN1 | 2.018031 | 0.00650046 |
| TRV-CAC1-6 | -0.0617375 | 0.00650963 |
| 100631378 | -0.0668486 | 0.00652437 |
| FMR1 | 0.92077729 | 0.00654709 |
| C6orf203 | 0.98436398 | 0.00658548 |
| SF3B1 | -1.0514096 | 0.00659651 |
| TAF9B | -1.0990143 | 0.00663897 |
| CNOT11 | -1.0572179 | 0.00668885 |
| CRYBA4 | 0.38654332 | 0.00669485 |
| MTO1 | -1.1603718 | 0.00673091 |
| TRAV8-1 | 1.84563991 | 0.00678125 |
| UBE2J1 | -1.0198904 | 0.0068024 |
| KLHL25 | -1.285691 | 0.00680514 |
| LRRCC1 | -1.5312399 | 0.00685139 |
| THEGL | -0.1220757 | 0.00686095 |
| PI4KAP2 | -0.8196212 | 0.0068653 |
| CTPS1 | -1.3343179 | 0.00689045 |
| 286178 | -0.1954858 | 0.00689266 |
| 101927354 | 0.60087592 | 0.00692278 |
| FADD | -1.0183752 | 0.00693321 |
| RBL2 | -0.9236179 | 0.00704665 |
| PRPF6 | -1.1318103 | 0.00705433 |
| TIMP2 | 0.68727047 | 0.00705985 |
| CLP1 | -1.6693277 | 0.00716941 |
| HNRNPA1L2 | -1.0966478 | 0.00717058 |

| | | |
|---|---|---|
| ZNF615 | -1.6773731 | 0.007186 |
| TMEM231 | -0.7578495 | 0.00718729 |
| RPL11 | 0.80355074 | 0.00719359 |
| LIMD2 | -0.6933652 | 0.00719508 |
| STX3 | -1.8995154 | 0.00720892 |
| EPC1 | 0.78714389 | 0.00723003 |
| RASA2 | -0.9936539 | 0.00724586 |
| GSTM2 | -1.1656561 | 0.00731874 |
| ANAPC16 | 0.44801157 | 0.00731939 |
| KIAA1143 | 0.96016829 | 0.00732395 |
| TBL3 | -1.9781264 | 0.00732487 |
| ARHGAP31 | 0.68991492 | 0.00733802 |
| SMTNL1 | -0.312313 | 0.00736461 |
| MIEF1 | 1.90175058 | 0.0073955 |
| COG1 | -1.4104406 | 0.00740826 |
| VAMP2 | 0.58738558 | 0.00741538 |
| ATP8B4 | 0.38085728 | 0.00742829 |
| CHST15 | -0.4990582 | 0.00743849 |
| CHD6 | 1.01674317 | 0.00755018 |
| PER2 | 0.57131148 | 0.00759145 |
| SAMD9L | 1.24988107 | 0.00761886 |
| DYNLL1 | 1.35984194 | 0.00763494 |
| SP140 | 0.71720025 | 0.0076506 |
| APP | 1.20911637 | 0.00767067 |
| SNRPD1 | 1.00105165 | 0.00767511 |
| 105372343 | -0.0163042 | 0.00767664 |
| MAF1 | -0.8845225 | 0.00769156 |
| MID1IP1 | -1.0261714 | 0.00774319 |
| SGK3 | -0.1857909 | 0.00775439 |
| MYLK | 0.75746519 | 0.0077572 |
| ERICH1 | 0.78381374 | 0.00791411 |
| GTPBP4 | -0.9064316 | 0.00793871 |
| TNRC6C | 0.8350841 | 0.00794383 |
| UGT2B7 | -0.0193474 | 0.00803449 |
| MRPL55 | 0.90802137 | 0.00806113 |
| 440461 | -1.0508116 | 0.00806902 |
| DAW1 | 0.13024316 | 0.00807314 |
| ITIH4-AS1 | 0.25391164 | 0.00811475 |
| C15orf39 | -1.520287 | 0.00814072 |
| ALOX5AP | 1.21924562 | 0.00818371 |
| TBC1D15 | -0.8523015 | 0.00823479 |
| DARS | -0.6665732 | 0.00823974 |
| SMURF1 | 1.28000433 | 0.00824093 |
| METTL4 | -0.9967867 | 0.00824549 |
| EIF4G2 | -0.8254862 | 0.00825503 |
| NGEF | -0.5431348 | 0.00828783 |
| LYPLA1 | -1.0357659 | 0.00830748 |
| SCD5 | 1.14140591 | 0.00831303 |

| | | |
|---|---|---|
| GGT5 | -0.0279276 | 0.00833571 |
| B2M | 0.73945414 | 0.00836373 |
| SOCS5 | -2.2004832 | 0.00838027 |
| RAB8A | -0.724801 | 0.00840174 |
| CIR1 | 0.61481329 | 0.00852355 |
| DIP2B | 1.42386405 | 0.00860937 |
| APOBEC3G | 1.05759258 | 0.00861754 |
| BEST3 | -0.1352957 | 0.00863017 |
| PMM2 | -1.4853158 | 0.00863626 |
| HDLBP | -1.2351121 | 0.00870398 |
| PRMT5-AS1 | 0.34183199 | 0.0087604 |
| ITGA4 | -1.0285647 | 0.00879145 |
| RPL26L1 | 0.75100076 | 0.00881645 |
| LAGE3 | 0.99979833 | 0.00882233 |
| ASB16-AS1 | -1.1429413 | 0.00882705 |
| CIZ1 | -1.347615 | 0.0088485 |
| 101927532 | -0.1014742 | 0.00887195 |
| DNA2 | 1.27566467 | 0.00888278 |
| TRBJ1-1 | 0.45537249 | 0.00896672 |
| ATP5C1 | 0.62221388 | 0.00900777 |
| TMBIM1 | -1.3688751 | 0.009011 |
| RGL3 | -0.203834 | 0.00903118 |
| TNRC6C-AS1 | -1.2282422 | 0.00904855 |
| AFG3L2 | -1.3052863 | 0.00908624 |
| CDIPT | -0.9153313 | 0.00909694 |
| ETV7 | 2.31850913 | 0.00911528 |
| MIR98 | 0.17132507 | 0.00912666 |
| CHCHD7 | 0.50809245 | 0.00913496 |
| LINC01320 | -0.0432528 | 0.00913844 |
| LINC00092 | -0.2839795 | 0.00915244 |
| TRAM1 | -0.5415112 | 0.00920708 |
| CD2 | 0.64656573 | 0.00922233 |
| MX2 | 1.52423311 | 0.00925288 |
| CYP4Z2P | -0.0227532 | 0.00928717 |
| AREG | 1.35626397 | 0.00931074 |
| TRIM52 | -0.9265631 | 0.00931238 |
| ZNF204P | -2.6568993 | 0.00936996 |
| MEMO1 | 0.70399658 | 0.00939557 |
| HOMER3 | -0.4910506 | 0.00942546 |
| LINC01583 | 0.14405012 | 0.00943506 |
| HABP4 | -1.1873 | 0.00953785 |
| ZNF28 | -1.3936058 | 0.0095417 |
| HMGN2P46 | 0.42004812 | 0.0095466 |
| CWC15 | 0.79572108 | 0.00955862 |
| PPFIA2 | -0.0555271 | 0.00961675 |
| UNC13B | 0.60941538 | 0.00962518 |
| NAA20 | 0.82106191 | 0.00965035 |
| FAM64A | -0.2170818 | 0.00969405 |

| | | |
|---|---|---|
| RAD23A | -0.6409211 | 0.00969687 |
| PPP1R16A | -1.5947558 | 0.00973808 |
| TLX2 | 0.2447893 | 0.00974101 |
| NOP2 | -1.2765039 | 0.00974983 |
| ETNK1 | -0.6427626 | 0.00979064 |
| POLE3 | -0.7148683 | 0.00987185 |
| UBE4A | -0.9089961 | 0.00987665 |
| CCDC142 | 1.83522993 | 0.00988888 |
| CCT4 | -0.6397568 | 0.00990195 |
| KIF15 | -0.6395566 | 0.00991674 |
| ZAP70 | -1.1805841 | 0.00992604 |
| C10orf2 | 2.05231379 | 0.00999185 |
| DDX49 | -1.2185196 | 0.01003113 |
| PCSK7 | -1.2848608 | 0.01003777 |
| ZNF766 | -1.2788218 | 0.01006155 |
| FKBP8 | -1.4972257 | 0.0100897 |
| PRR29 | -0.6485482 | 0.01010155 |
| LINC01023 | 0.44909424 | 0.01010807 |
| AIMP1 | 0.65542379 | 0.01017038 |
| RBBP4 | -0.5663827 | 0.01023719 |
| CALR | -1.0452629 | 0.01023861 |
| GFRA3 | -0.0318063 | 0.01030342 |
| TRPM7 | -0.8302103 | 0.01042067 |
| LAMB4 | -0.2714459 | 0.01046587 |
| GCM2 | 0.27838172 | 0.01049474 |
| 100129216 | 0.14327236 | 0.01054874 |
| KHDC1 | -1.3159579 | 0.0105657 |
| SAP30L-AS1 | 1.61844836 | 0.01058437 |
| MT-ND1 | 0.58422906 | 0.01060605 |
| MIATNB | -1.4295568 | 0.01066485 |
| TRDJ2 | -0.3003989 | 0.01066732 |
| NDUFB5 | 0.6594227 | 0.01067445 |
| 101929766 | -0.0530847 | 0.01068957 |
| IVNS1ABP | -1.0869509 | 0.01077871 |
| NEPRO | -0.8879274 | 0.010782 |
| KLHL8 | -1.9978457 | 0.01078416 |
| SLC23A2 | 0.91569292 | 0.01081334 |
| ELF1 | -0.5557855 | 0.01082488 |
| SEC62 | 0.99045537 | 0.01082844 |
| TSPYL1 | 1.01682428 | 0.01083091 |
| PDGFRA | -0.05926 | 0.01084008 |
| IPO7 | 1.67170399 | 0.01084507 |
| OAS3 | 1.70116426 | 0.01092008 |
| ACHE | 0.12626758 | 0.01092058 |
| KARS | -0.8445851 | 0.01093308 |
| MT-ATP8 | 0.9242909 | 0.01097536 |
| MAP2K5 | -1.4654151 | 0.0109821 |
| HPS4 | -1.3671643 | 0.01098658 |

| | | |
|---|---|---|
| SF3A3 | -0.6865718 | 0.01103748 |
| ZNF442 | -1.9627622 | 0.01106116 |
| C6orf229 | -0.476879 | 0.01110104 |
| OR14A16 | -0.0288051 | 0.01123214 |
| PHC1 | 0.58590421 | 0.01128382 |
| CMPK1 | -0.5927347 | 0.01130089 |
| ZNF414 | -0.8518988 | 0.01132147 |
| TACC3 | -1.1900368 | 0.01140675 |
| BOLA3 | 1.05563388 | 0.01143534 |
| CCND2 | 1.03245675 | 0.01148442 |
| ZP1 | 0.44000697 | 0.01149297 |
| GIPR | 0.65994756 | 0.01155633 |
| 101928710 | 1.68797513 | 0.01157584 |
| HSD11B1L | -1.0752249 | 0.01163889 |
| EZR | -0.8460915 | 0.01164479 |
| MT-CYB | 0.61296007 | 0.01168591 |
| PLA2G12A | -0.8247676 | 0.01170861 |
| DCAF12 | -1.5467431 | 0.0117163 |
| IL1RN | 0.14174447 | 0.0117334 |
| RAD50 | -0.8935761 | 0.0117971 |
| P4HB | -0.9992131 | 0.01187762 |
| KIAA1033 | -1.1462309 | 0.01191727 |
| ANAPC1 | 1.19346407 | 0.01192938 |
| ST3GAL4 | 1.10040763 | 0.01195022 |
| FBXL5 | -1.1485534 | 0.01196572 |
| HNRNPH2 | -0.2693786 | 0.0120035 |
| RNF6 | -1.2953098 | 0.01204001 |
| MLXIP | 1.46539675 | 0.01210466 |
| FAM120AOS | -0.7085436 | 0.01215168 |
| ZNF227 | -1.8173368 | 0.01219099 |
| CAPZB | 0.51428571 | 0.01219413 |
| MYL12B | 0.74383313 | 0.01222162 |
| B3GNT2 | -1.0377916 | 0.01222763 |
| TGM5 | 0.22206882 | 0.01226924 |
| FLYWCH1 | 1.66566068 | 0.01227046 |
| IAH1 | 0.68200708 | 0.01227439 |
| GALNT18 | 0.07096211 | 0.0122765 |
| TCTN3 | -0.7972787 | 0.01228978 |
| ZNF451 | -1.0152958 | 0.0123308 |
| RBM8A | 0.85029033 | 0.01234421 |
| RPL7 | 0.33458411 | 0.01235592 |
| PRDX4 | 1.11621942 | 0.01235892 |
| LCN12 | 0.12252254 | 0.01236305 |
| IL17D | -0.3958191 | 0.01241604 |
| GRK5 | -1.7339777 | 0.01242918 |
| MYC | -2.1692651 | 0.01246466 |
| EEPD1 | -1.6404512 | 0.01248154 |
| OASL | 1.26091226 | 0.01249329 |

| | | |
|---|---|---|
| LCT | -0.2189236 | 0.01252532 |
| SEC22A | 0.7330418 | 0.01253574 |
| CD48 | 0.50718256 | 0.01256183 |
| CNTNAP2 | -0.3580093 | 0.01262406 |
| TMEFF2 | -0.0365116 | 0.01266762 |
| XAF1 | 2.26529516 | 0.0127633 |
| MON1B | -1.7146381 | 0.01277637 |
| PDZD11 | 0.89881276 | 0.01279188 |
| IGSF3 | 0.2219598 | 0.01280893 |
| USP39 | -0.7014758 | 0.0129165 |
| NDUFAF3 | 0.72794824 | 0.01293238 |
| ARMC1 | -1.3955034 | 0.0129465 |
| RPL19 | 0.48839637 | 0.01294807 |
| FANCI | -1.4854354 | 0.01294814 |
| CREBZF | -0.8249121 | 0.01299003 |
| TCP11X2 | -0.019734 | 0.01300018 |
| ZNF141 | -1.5477019 | 0.01302798 |
| ZRSR2 | 0.72547852 | 0.01312447 |
| ZNF619 | -1.1530419 | 0.01320671 |
| NT5DC4 | 0.30760474 | 0.01331186 |
| GOLGA6L9 | -0.1139544 | 0.01332844 |
| MSRB2 | 1.02358694 | 0.013335 |

**Table S3. List of top 500 differently expressed genes btw Temra blood and TDL**

| Gene | logFC | P.Value |
|---|---|---|
| LRRC75B | 0.5053442 | 1.55E-07 |
| CCR7 | 2.41259857 | 1.61E-06 |
| PRSS23 | -7.1207014 | 6.35E-06 |
| NELL2 | 1.77387907 | 1.02E-05 |
| TRDD3 | 0.04340317 | 1.48E-05 |
| FCGR2C | -0.386346 | 1.78E-05 |
| GTF2IP4 | 0.55344546 | 2.04E-05 |
| 102724760 | -0.0334293 | 2.92E-05 |
| PASK | 2.07824315 | 4.90E-05 |
| BCL2L15 | -0.6994543 | 5.35E-05 |
| CCR2 | 5.16675422 | 5.36E-05 |
| MX1 | -2.8790413 | 6.89E-05 |
| IFI6 | -2.815211 | 7.06E-05 |
| PDZD4 | -0.4224399 | 8.98E-05 |
| CX3CR1 | -4.7693658 | 0.00010113 |
| IFIT1 | -6.5941509 | 0.00012473 |
| ANKRD44 | -0.8639327 | 0.00013141 |
| DRAP1 | -0.7904937 | 0.00015457 |
| SLC12A5 | -0.2286702 | 0.00016643 |
| GTSE1-AS1 | -0.3663557 | 0.00017111 |
| KDM3A | 1.19614758 | 0.00017603 |
| NME8 | -0.3890447 | 0.00018216 |
| CREBRF | 0.81672654 | 0.00021467 |
| CXCR2 | -0.5587018 | 0.00023363 |
| LRFN2 | -0.5372958 | 0.00024281 |
| ADRA2B | 0.25547115 | 0.00028244 |
| IFITM1 | -2.0511273 | 0.00030283 |
| PPIC | -0.058227 | 0.00035572 |
| PODXL | -0.7966535 | 0.00038 |
| FSTL3 | 0.89746534 | 0.00039269 |
| CD55 | 1.44698215 | 0.00040083 |
| PSME3 | 1.04568353 | 0.00043053 |
| PLSCR1 | -2.031623 | 0.00048512 |
| IFNG | -2.1277705 | 0.00051105 |
| ISG15 | -2.248102 | 0.00051622 |
| SAMD9L | -1.7229716 | 0.00052751 |
| TAF2 | 1.90331299 | 0.00053721 |
| NHSL1 | 0.36484 | 0.00064691 |
| TBC1D4 | 1.94434196 | 0.0006651 |
| S1PR4 | 0.76003488 | 0.00069303 |
| APOBEC3A | -0.0489475 | 0.00078686 |
| LINC00867 | -0.1375593 | 0.00080757 |
| NMUR1 | -0.344687 | 0.00081196 |
| MAMLD1 | 0.36698829 | 0.00083436 |
| 101928718 | 0.29682309 | 0.00088788 |
| 100134317 | -0.0803738 | 0.0009662 |

| | | |
|---|---|---|
| PER2 | -0.7380531 | 0.00097912 |
| GFM1 | 1.56350866 | 0.0010063 |
| ACTR8 | 1.04003158 | 0.00106476 |
| BCKDK | 2.79847735 | 0.00113551 |
| CPNE8 | -1.3125094 | 0.00113797 |
| TMEM259 | 1.85795979 | 0.0011813 |
| ZNF587 | 1.76284517 | 0.00120711 |
| MIR630 | -0.4452918 | 0.00121787 |
| FCGR3A | -5.9445048 | 0.0012502 |
| ANKRD20A1: | -0.6409002 | 0.00126822 |
| 727896 | 0.86510753 | 0.0013019 |
| MIR8085 | 0.06093356 | 0.00131627 |
| TACC3 | 1.57865515 | 0.00135663 |
| PFKL | 1.51669148 | 0.00136898 |
| ZNF615 | 2.06716965 | 0.00139565 |
| PIGV | -1.757117 | 0.00140167 |
| FAR2P2 | -0.0241519 | 0.00141321 |
| MITD1 | -0.8722709 | 0.00142561 |
| MIR1277 | -0.1485273 | 0.00155077 |
| UBE2F | -1.030744 | 0.00155797 |
| TRBV27 | 1.35411962 | 0.00156706 |
| ISG20 | -1.2454364 | 0.00157416 |
| RRAGB | 1.12154956 | 0.00159156 |
| ELOVL1 | 0.91655074 | 0.00162063 |
| PGD | 1.77360935 | 0.00163462 |
| ZNF853 | -1.315294 | 0.00163576 |
| TMEM204 | 1.33268923 | 0.00166579 |
| POLR3H | 1.13634226 | 0.00177861 |
| PIM1 | -1.3066095 | 0.00179463 |
| FGR | -4.1181286 | 0.00180215 |
| FANCE | 2.20540572 | 0.00181798 |
| BST2 | -0.9528353 | 0.00184401 |
| 400927 | -0.4900776 | 0.00184588 |
| PSME2 | -1.1013607 | 0.0018807 |
| 100506124 | -0.7605731 | 0.00192546 |
| 102467081 | 1.33062103 | 0.00195739 |
| XIAP | -1.5569127 | 0.00196774 |
| C9orf85 | -1.1913723 | 0.00202574 |
| SIRT3 | 1.38227997 | 0.00203989 |
| DHRS13 | 1.28918652 | 0.00205407 |
| TST | -1.4556776 | 0.00206451 |
| CMKLR1 | -0.4701927 | 0.00210164 |
| SERINC5 | 2.31268007 | 0.0021786 |
| NOXO1 | 0.31029113 | 0.00218984 |
| CA7 | -0.5700055 | 0.00220685 |
| THEMIS2 | 1.8311059 | 0.00221265 |
| DDX27 | 0.86026989 | 0.00232779 |
| CYB561A3 | 1.99046203 | 0.00232832 |

| | | |
|---|---|---|
| TRBV12-4 | -4.6072249 | 0.0023927 |
| DCXR | -0.9040596 | 0.00241295 |
| CLDND1 | 1.2187506 | 0.00247766 |
| CD320 | 1.34581434 | 0.00251504 |
| PTGDS | -0.2427731 | 0.00253169 |
| MINA | 1.60764612 | 0.00260718 |
| TIMM17B | -0.7129295 | 0.00264823 |
| RAB14 | 0.88921892 | 0.00266063 |
| GPI | 0.80359874 | 0.00275323 |
| MPHOSPH10 | 0.97038813 | 0.00282669 |
| NFE2L2 | 1.10317266 | 0.00289837 |
| TAT-AS1 | -0.4694859 | 0.0029006 |
| USP24 | 0.86053504 | 0.00293991 |
| TTF2 | 1.65182474 | 0.0029795 |
| HIST1H4F | -1.1729204 | 0.0030344 |
| NUP93 | 1.24488944 | 0.00306975 |
| PCDH1 | -0.327701 | 0.00307295 |
| PKM | 1.30884771 | 0.00309436 |
| PPP6C | 0.71070984 | 0.00314321 |
| KIN | -0.7346962 | 0.00320641 |
| GZMH | -2.8861133 | 0.00321742 |
| LSP1 | 0.79831219 | 0.00324696 |
| ZCCHC8 | 0.75146856 | 0.00325024 |
| DHX35 | 2.26204266 | 0.00326465 |
| WASH2P | -0.6344471 | 0.00327099 |
| OSBPL9 | 1.11286568 | 0.00330285 |
| NDFIP2 | -2.3023425 | 0.0033129 |
| H2AFY | 0.96407428 | 0.00333518 |
| IFI44 | -1.9660097 | 0.00334238 |
| PHF19 | 1.26524654 | 0.00338371 |
| MIR103A2 | -0.3596842 | 0.00340821 |
| NSUN5 | 1.54372538 | 0.00341438 |
| ACOT13 | -0.9907157 | 0.00342637 |
| RIMKLB | 1.51553175 | 0.00343628 |
| RBM43 | -1.3360032 | 0.00346414 |
| ODF3B | -0.9259935 | 0.00347482 |
| UBOX5-AS1 | 0.33392732 | 0.00347811 |
| IGF1R | 1.43144498 | 0.0035153 |
| RPA1 | 0.98842971 | 0.00357624 |
| IGSF8 | 1.6414874 | 0.00359234 |
| PXT1 | 0.32498504 | 0.00359684 |
| INO80C | -0.9242785 | 0.00361153 |
| PHF11 | -0.9199833 | 0.00365058 |
| MCM8-AS1 | -0.2191464 | 0.00365902 |
| CMTM3 | 0.81584372 | 0.00367583 |
| TIPARP | 1.64270925 | 0.00371105 |
| C21orf62-AS | -0.1660465 | 0.00373317 |
| 390937 | -0.3973204 | 0.00374378 |

| | | |
|---|---|---|
| IMPDH2 | 0.68431943 | 0.00379985 |
| NPIPB9 | 0.05665313 | 0.00391993 |
| COMMD1 | -1.1684291 | 0.00409952 |
| PHLDA1 | 1.73029589 | 0.00415807 |
| RPLP0 | 0.46030376 | 0.00418219 |
| TM9SF4 | 1.51480082 | 0.0042293 |
| TRDV3 | -1.1700769 | 0.00426133 |
| PTBP2 | 1.72872432 | 0.00426207 |
| BZW2 | 1.33578769 | 0.0042663 |
| TRBV5-4 | 3.23429715 | 0.00428438 |
| ZNF48 | 1.63461813 | 0.0042908 |
| RPL15 | -0.306687 | 0.00432068 |
| NMRAL1 | 0.03462038 | 0.00438155 |
| MAST2 | 1.8479768 | 0.00439275 |
| TESPA1 | 1.15417755 | 0.00441194 |
| CD28 | 1.08871566 | 0.00442187 |
| ACMSD | -0.1038108 | 0.00450854 |
| ATP6AP1 | 1.49815178 | 0.00454379 |
| 105376719 | -0.299603 | 0.00454466 |
| CCL5 | -0.4081894 | 0.00454569 |
| MFSD12 | 1.40945668 | 0.004584 |
| GNAL | -0.8274682 | 0.00460752 |
| C5orf56 | -1.5434521 | 0.00461328 |
| IL7R | 1.00303789 | 0.00463764 |
| PTMA | -0.6106038 | 0.00464231 |
| RNU2-2P | -1.1660888 | 0.00466782 |
| ZNF451 | 1.17142932 | 0.00468141 |
| CD27 | 0.82282016 | 0.00474364 |
| CCDC12 | -0.7676413 | 0.0047497 |
| MYOZ2 | 0.07176057 | 0.0047761 |
| CD3G | -0.9958799 | 0.00479455 |
| TSPYL2 | 1.16762711 | 0.00482593 |
| PM20D2 | -1.7911285 | 0.00491177 |
| ABTB1 | 1.32121811 | 0.00497685 |
| HIST1H2BE | -0.6573847 | 0.00512662 |
| TRBV20-1 | -2.0838686 | 0.00515441 |
| VASH1 | -0.8138645 | 0.00523455 |
| RAB1B | -1.2631425 | 0.00536186 |
| HSPBP1 | 0.85999527 | 0.00538211 |
| CCDC50 | -2.198589 | 0.00543352 |
| CNPY4 | 0.80853696 | 0.00547506 |
| LEXM | -0.6996476 | 0.00550603 |
| ACSL6 | 2.33615977 | 0.00551504 |
| CMAS | 0.96682315 | 0.00551806 |
| 101927866 | 0.06047603 | 0.00552018 |
| UBAC2 | 0.93003873 | 0.00553986 |
| CPSF7 | 0.97543074 | 0.00556837 |
| 375196 | 0.32666291 | 0.00560993 |

| | | |
|---|---|---|
| MAFG | 1.7127256 | 0.00561635 |
| DUSP5 | -0.7041811 | 0.00568225 |
| CNBD2 | -0.4303747 | 0.00576349 |
| C7orf73 | -0.4437267 | 0.00585561 |
| REG4 | 2.39483131 | 0.00593183 |
| RARRES3 | -1.3096297 | 0.00605114 |
| TSG101 | -0.6120116 | 0.00610427 |
| APEH | 1.10449174 | 0.0061678 |
| KPNA5 | -0.8190007 | 0.00619538 |
| UBE4B | -1.318913 | 0.00621873 |
| RTP4 | -1.8031751 | 0.00622494 |
| RECQL | 1.06487926 | 0.00624614 |
| SPON2 | -0.8933679 | 0.00625963 |
| ZNF124 | 1.50124705 | 0.00626471 |
| ZNF821 | 1.54583915 | 0.00627972 |
| B4GALT4-AS | -0.2409321 | 0.00633213 |
| TXNRD1 | 1.18748908 | 0.00637619 |
| NEB | 1.42355526 | 0.00637723 |
| C21orf2 | 1.86971863 | 0.00651788 |
| POP4 | -0.6638464 | 0.00652015 |
| SLC39A13 | 1.20942599 | 0.00653849 |
| EGR1 | -0.4212298 | 0.00654923 |
| FGF9 | 1.51686768 | 0.00658214 |
| COPZ2 | -1.4060894 | 0.00661816 |
| SUV39H1 | 2.41599654 | 0.00670794 |
| SPNS1 | 1.39872032 | 0.00675099 |
| CDK10 | 1.09822908 | 0.00681201 |
| PDE6D | -0.791776 | 0.00693954 |
| NOCT | -1.0846645 | 0.00698855 |
| ZNF296 | -0.8934355 | 0.00708848 |
| IGLV2-11 | -0.4522251 | 0.00734657 |
| 105369174 | 1.19810571 | 0.0074077 |
| PGRMC1 | 1.27987327 | 0.00753635 |
| DDX20 | -1.2154676 | 0.00753907 |
| RPL26L1 | -0.7686799 | 0.00754366 |
| SNORD77 | 0.22820248 | 0.00755285 |
| SNORA31 | 0.16974484 | 0.00756461 |
| POMT1 | 1.84155152 | 0.00764249 |
| DKK3 | 1.91231163 | 0.00766176 |
| N4BP1 | -0.6532793 | 0.00778635 |
| MCRIP2 | 0.61812717 | 0.00781004 |
| 100506985 | 1.22801298 | 0.0078854 |
| TRAV13-1 | -1.6462369 | 0.00788886 |
| PIGM | 1.31612783 | 0.00793692 |
| TRAV8-1 | -1.8017875 | 0.00799238 |
| TBC1D10A | 1.51881216 | 0.00804007 |
| SCD | -0.7366692 | 0.00804159 |
| ABHD3 | 0.78296115 | 0.00804749 |

| | | |
|---|---|---|
| NBPF25P | 0.30363956 | 0.00811775 |
| WDYHV1 | 1.39960569 | 0.00813137 |
| CALU | 1.07829435 | 0.0082233 |
| PLGRKT | -0.690773 | 0.00830659 |
| TMEM14A | -0.6898617 | 0.00834153 |
| R3HDM4 | 1.07893467 | 0.00840093 |
| UBB | -0.8034947 | 0.00841039 |
| DLEU7-AS1 | 0.10584088 | 0.00846611 |
| ENDOV | 1.36971735 | 0.00849925 |
| OGFR-AS1 | 0.0888112 | 0.00861802 |
| TMCC1 | -1.3546819 | 0.00871067 |
| PPP1R15A | 2.44753743 | 0.0087116 |
| 101928710 | -1.7642407 | 0.00871418 |
| MFAP2 | 0.58857831 | 0.00878734 |
| BATF | -0.8803834 | 0.00879331 |
| RELL1 | 1.26905389 | 0.00879961 |
| ZNF619 | 1.22897654 | 0.00882077 |
| CLCF1 | -1.558249 | 0.00883849 |
| SNORD87 | -0.5943635 | 0.00886158 |
| APPL1 | 0.8064921 | 0.00901562 |
| AZIN2 | 1.61266528 | 0.00901932 |
| SMYD5 | 1.65598035 | 0.00912923 |
| TTC19 | 1.0841075 | 0.00916957 |
| HSPA5 | 0.89940604 | 0.0091907 |
| FAM86C2P | 1.00649659 | 0.00919541 |
| LIFR | 0.04076418 | 0.00920382 |
| SAE1 | 0.91628259 | 0.00931411 |
| MAZ | 0.7673601 | 0.00937907 |
| LRRN3 | 4.40935941 | 0.00938598 |
| PTPN21 | -0.2764728 | 0.0095315 |
| ACACA | 0.20998066 | 0.00953794 |
| PCDHGA11 | 0.01952972 | 0.0096387 |
| TARS2 | 1.60592046 | 0.00982406 |
| C6orf229 | -0.4835004 | 0.01016988 |
| MIR6747 | -0.0122926 | 0.01022373 |
| ATP10A | 1.27243674 | 0.01028256 |
| ZNF362 | -1.1603674 | 0.01031639 |
| CMPK2 | -2.8123955 | 0.01032152 |
| TCTEX1D2 | -1.5689906 | 0.01048341 |
| CLEC18A | -0.1582676 | 0.01051746 |
| CUTA | -0.0826439 | 0.01069925 |
| C14orf80 | -1.8013142 | 0.01097675 |
| SUN1 | 1.02549752 | 0.01098654 |
| PLEKHB1 | 2.67837104 | 0.01099273 |
| CCNL1 | 0.50000752 | 0.01106547 |
| GLIPR2 | -0.877775 | 0.0110964 |
| ZNF593 | -1.0797358 | 0.01113207 |
| TRAV22 | 2.13766001 | 0.01119218 |

| | | |
|---|---|---|
| MRPS31 | -0.6422509 | 0.0112393 |
| CRTC3-AS1 | 0.48151902 | 0.01128484 |
| LGALS9 | -0.8256168 | 0.01133633 |
| 102723897 | 0.29640908 | 0.01138789 |
| C1QBP | 0.4938215 | 0.01146653 |
| UQCRH | -1.0108257 | 0.01147578 |
| SESN1 | 1.6971156 | 0.01163044 |
| TNNT3 | 0.94997333 | 0.01167258 |
| COG6 | 1.31246858 | 0.01167561 |
| GOLGA8R | 0.07523398 | 0.01170566 |
| ACD | 0.84812488 | 0.01182199 |
| ZNF419 | 1.61378187 | 0.01184119 |
| TSHZ3 | -0.704966 | 0.01188883 |
| CBR4 | -0.8912795 | 0.01199712 |
| 102724474 | -0.1919745 | 0.01203066 |
| CECR7 | -0.434701 | 0.01215982 |
| ZNF585A | -0.9244779 | 0.01220042 |
| ITGA4 | 0.97658622 | 0.01224051 |
| TMED2 | 0.8453442 | 0.01225419 |
| KIAA0391 | 0.93713965 | 0.01231992 |
| ZNF813 | 2.09204947 | 0.01241857 |
| TRAV16 | -1.6211091 | 0.01242936 |
| NR5A2 | -0.5785236 | 0.01248161 |
| C1orf74 | -1.7538547 | 0.01249847 |
| 100130298 | -1.1359469 | 0.01249933 |
| B2M | -0.6938407 | 0.01256818 |
| DNAAF5 | 1.18728653 | 0.01274472 |
| EBLN3P | 0.98081777 | 0.01277349 |
| PSMA5 | -0.4941256 | 0.01284116 |
| IRF9 | -1.16894 | 0.01286286 |
| 729603 | -0.7621566 | 0.01300863 |
| TRGV3 | -0.9546082 | 0.01301926 |
| UQCC3 | -0.996916 | 0.01302562 |
| FAM213A | 1.19309337 | 0.01303097 |
| PDE3B | 0.95322149 | 0.01307726 |
| YIPF6 | -1.1119755 | 0.013144 |
| USP14 | 0.86001128 | 0.01319049 |
| IFRD2 | 1.47506199 | 0.01337534 |
| CCDC144NL-/ | 0.095502 | 0.01341818 |
| PNPLA2 | 1.05678091 | 0.01355038 |
| SLC4A2 | 1.18686478 | 0.01357051 |
| SLC43A3 | 1.48248486 | 0.0135735 |
| NEK9 | -0.9291263 | 0.01359547 |
| LDAH | 1.9781593 | 0.01360346 |
| FAAH | 1.15415625 | 0.01370264 |
| ZNF581 | 1.13546575 | 0.01371454 |
| RABIF | -0.8666132 | 0.01372104 |
| MIR4750 | 0.01876001 | 0.01379677 |

| | | |
|---|---|---|
| SNORD98 | 0.147321 | 0.01384253 |
| FAM76A | -0.9964 | 0.01396986 |
| TYW5 | -1.2790785 | 0.01400923 |
| XAF1 | -2.2291393 | 0.01406673 |
| IL6ST | 1.40966525 | 0.01408898 |
| STARD5 | 1.52561651 | 0.01419474 |
| CATSPERB | 0.11100448 | 0.01421484 |
| MIR301A | 0.01743905 | 0.01428774 |
| DCAF11 | 1.05662125 | 0.01432072 |
| UQCRHL | -1.0614018 | 0.01434056 |
| ASIC1 | 0.68163858 | 0.01438134 |
| ERVK13-1 | 1.28899292 | 0.01446447 |
| 100240734 | -0.5277602 | 0.01448189 |
| CEP128 | 2.09108193 | 0.01450943 |
| ARMCX4 | 1.84356216 | 0.01458305 |
| WDR4 | 1.82082728 | 0.01458779 |
| 124871 | -0.2652793 | 0.01461566 |
| POLR2G | -0.8058264 | 0.0147065 |
| PEX11G | -0.7933798 | 0.01480051 |
| HLA-H | 1.68510686 | 0.0148044 |
| C7orf49 | 0.9935504 | 0.01483046 |
| OAS3 | -1.6177172 | 0.01484847 |
| BRD2 | -0.1397002 | 0.0148945 |
| RSF1 | -0.7083147 | 0.01492589 |
| SUN2 | 1.24306024 | 0.01511493 |
| SAP30L | -1.0477837 | 0.01511549 |
| CENPH | -1.3785388 | 0.01521738 |
| CBX3P2 | 0.85284386 | 0.01550052 |
| BCAS2 | -0.5048568 | 0.01564322 |
| ITGB7 | 1.04229966 | 0.01572063 |
| CEBPZ | 0.8462657 | 0.01572426 |
| MAN1B1 | 1.49659001 | 0.01589193 |
| SEPW1 | -0.7263633 | 0.01591771 |
| RPL29 | -0.5674685 | 0.01593955 |
| FGFBP3 | 0.80962912 | 0.01596284 |
| MCF2L-AS1 | 1.13003442 | 0.01614288 |
| TCEA1 | 0.51884256 | 0.01617832 |
| ENO2 | 0.89871054 | 0.01618633 |
| TMSB10 | -1.2407234 | 0.01625545 |
| MEN1 | 1.17191638 | 0.01639662 |
| FANCM | -1.1913007 | 0.01642825 |
| ANKRD1 | 0.1821807 | 0.0164286 |
| ZNF837 | 2.15907855 | 0.01650641 |
| FAM122A | -1.0911027 | 0.0165172 |
| TAB2 | 0.85610238 | 0.01653934 |
| ORC5 | 1.12027892 | 0.01657275 |
| IFIH1 | -1.3344777 | 0.01657489 |
| TRGV2 | -1.8650302 | 0.01667573 |

| | | |
|---|---|---|
| PTPRF | -0.5596316 | 0.01669511 |
| THAP9 | 1.54186514 | 0.0167197 |
| TSPOAP1 | -1.6586655 | 0.01679861 |
| ZNF442 | 1.82969343 | 0.0168704 |
| HSCB | 0.70317644 | 0.01689215 |
| TSN | 0.55272897 | 0.01689833 |
| PLCG1-AS1 | -0.8004639 | 0.01691501 |
| TRIM45 | 0.35685953 | 0.01691848 |
| OAS1 | -2.5478532 | 0.0169462 |
| MAST4 | -1.1889737 | 0.01695907 |
| TRPM7 | 0.76447731 | 0.01715004 |
| SLC48A1 | -1.1003743 | 0.01730458 |
| GON4L | -0.742323 | 0.0173476 |
| FMNL3 | -1.5023649 | 0.01740255 |
| YBX3 | 1.10201207 | 0.01745196 |
| ADI1 | 0.81619673 | 0.01747101 |
| GFOD1 | -1.2586101 | 0.01749878 |
| SEC11C | -0.5910301 | 0.01751506 |
| NPTXR | -0.8293644 | 0.01754779 |
| MRPL24 | -0.645058 | 0.01757849 |
| SNRPD1 | -0.8755085 | 0.01764777 |
| TRAV12-1 | 1.69932908 | 0.01764777 |
| KTN1 | -0.7091717 | 0.017657 |
| 102724814 | 0.66469991 | 0.01765756 |
| ZNF408 | 1.53498921 | 0.01766438 |
| ECM2 | 0.61716551 | 0.01767631 |
| GCNT1 | -2.1091598 | 0.01771322 |
| MED16 | 1.13023859 | 0.01774501 |
| YIF1B | 0.86371683 | 0.01780767 |
| SLC7A6 | 0.90480722 | 0.01789058 |
| CLUAP1 | 1.08108775 | 0.01790611 |
| DOK1 | 1.39823482 | 0.01792833 |
| VCP | 1.01877682 | 0.01797792 |
| LINC01481 | -0.8490009 | 0.01799173 |
| CIRBP | -0.4182213 | 0.01802861 |
| MINCR | -1.0896885 | 0.01814636 |
| DHCR24 | -1.1702256 | 0.01820136 |
| PSME1 | -0.828842 | 0.01827473 |
| SATB1-AS1 | 0.7233765 | 0.01828935 |
| 100131655 | 0.58061884 | 0.01830858 |
| 100507195 | -0.8001644 | 0.01847958 |
| VPS51 | 1.16848769 | 0.01849156 |
| SH3D21 | 0.29919623 | 0.01874232 |
| ADAR | -0.6690164 | 0.01877383 |
| PTPRN | 0.26609339 | 0.01878972 |
| PITPNC1 | 0.73490417 | 0.01894121 |
| GPR183 | 0.74873243 | 0.01903782 |
| DPH1 | 1.58219892 | 0.01904379 |

| | | |
|---|---|---|
| ITCH | -0.9552473 | 0.0190678 |
| EPB41L4A | -1.2312746 | 0.01907038 |
| 100505716 | 0.77488018 | 0.01908951 |
| MAP3K6 | -1.4182825 | 0.01922882 |
| MTM1 | 1.11280606 | 0.01924714 |
| STXBP2 | 0.9530022 | 0.01929481 |
| FUNDC2P2 | -0.2304932 | 0.01930534 |
| ZNF467 | 0.671793 | 0.01932104 |
| RARS | 0.63136844 | 0.01937136 |
| NIPA1 | 1.19845835 | 0.01938505 |
| APLF | 1.38493117 | 0.01941832 |
| AGPAT4-IT1 | -0.628129 | 0.01944261 |
| UBE2L6 | -0.793232 | 0.01947994 |
| TRDMT1 | -0.908568 | 0.01949418 |
| MRPL16 | -0.532315 | 0.01954471 |
| TRAF1 | 0.85965913 | 0.01956003 |
| MIR30C1 | 0.00864844 | 0.01963381 |
| PAK2 | 0.81134867 | 0.01965768 |
| COMT | 0.71540097 | 0.019663 |
| HSF2BP | 0.18749796 | 0.01967007 |
| IFIT5 | -1.0768672 | 0.01968924 |
| GPR107 | -0.9421368 | 0.01978329 |
| 101928021 | -0.1714406 | 0.01985598 |
| CHD7 | 1.04714602 | 0.01985652 |
| CDIPT | 0.80357494 | 0.01992221 |
| MXD3 | -1.3241375 | 0.01996017 |
| EZR | 0.77148494 | 0.02000034 |
| GRPEL1 | -0.5106724 | 0.02014585 |
| ZNF763 | -1.5266261 | 0.02014749 |
| CERKL | 0.96345125 | 0.02034851 |
| CRLF3 | 0.82251884 | 0.02036104 |
| SOD2 | -0.8467088 | 0.02045783 |
| ZNF420 | 1.18800201 | 0.02048671 |
| ZFP28 | 1.86546997 | 0.02050988 |
| FAM134B | 1.25322191 | 0.0206084 |
| AFG3L2 | 1.1381836 | 0.0206551 |
| SLC39A8 | 1.22129027 | 0.02067322 |
| DOCK9-AS2 | 0.90201577 | 0.02067963 |
| SAMD9 | -0.8359764 | 0.02068594 |
| GPBP1L1 | 0.60103783 | 0.02070815 |
| THUMPD1 | 0.68335473 | 0.02073201 |
| DDX56 | 1.14778359 | 0.02080039 |
| FBXL8 | 1.01076686 | 0.02087603 |
| SEC24B-AS1 | -1.1283602 | 0.02092781 |
| DBNL | 0.89208331 | 0.0210157 |
| PLEKHS1 | -0.2040479 | 0.02116551 |
| HSP90AB4P | -1.00888 | 0.02125622 |
| 100506922 | -0.8964493 | 0.02135985 |

| | | |
|---|---|---|
| EPSTI1 | -1.9516467 | 0.02143967 |
| TRAV1-1 | 1.07128372 | 0.02146796 |
| HSPB11 | -1.1385401 | 0.02146977 |
| EIF2B5 | 1.27889945 | 0.02157663 |
| MIR423 | -0.0337732 | 0.02166127 |
| CTU2 | 1.92001814 | 0.02173439 |
| RPL21P28 | -0.8742091 | 0.02176172 |
| NPAT | 0.81687948 | 0.02176484 |
| CSDE1 | 0.89041254 | 0.02181616 |
| SCAF8 | -1.1499059 | 0.02184337 |
| PTPN6 | 1.12648498 | 0.02187917 |
| HTRA4 | -0.4364924 | 0.02194638 |
| EPHX2 | 1.40715284 | 0.02201152 |
| AHNAK2 | -0.053015 | 0.02202259 |
| ZBTB8OS | -0.936052 | 0.02203941 |
| DEK | 0.49574192 | 0.02206429 |
| ITGAM | -0.4071545 | 0.02208569 |
| IL4I1 | -0.7366788 | 0.02210201 |
| ZSCAN2 | 0.87935197 | 0.02211352 |
| ZNF584 | 2.13868541 | 0.02211963 |
| CARD16 | -1.3285288 | 0.02220592 |
| CENPE | -2.0673802 | 0.02225994 |

**Table S4. List of top 500 differently expressed genes btw R1+27- and R1-27+**

| Gene | logFC | P.Value |
|---|---|---|
| ADAM12 | -8.1243175 | 8.68E-07 |
| ME1 | -9.5472214 | 1.85E-06 |
| CAPG | -6.5694752 | 1.86E-06 |
| IL23R | -8.1009247 | 2.82E-06 |
| NME8 | 7.51224434 | 2.86E-06 |
| SCRN1 | -7.8381395 | 3.71E-06 |
| IL12RB2 | -6.9624687 | 3.78E-06 |
| AFF3 | -6.6648709 | 3.84E-06 |
| CD40LG | -7.6389594 | 6.38E-06 |
| RBM24 | -6.6038793 | 7.05E-06 |
| TIAM1 | -6.1022818 | 8.32E-06 |
| SLC4A10 | -9.6539569 | 9.17E-06 |
| PROK2 | 6.56973651 | 1.01E-05 |
| RP11-603B24 | -6.1757585 | 1.63E-05 |
| FGFBP2 | 5.25643089 | 1.87E-05 |
| CXCR2 | 5.59460678 | 1.98E-05 |
| RP11-563D1( | -4.7192435 | 2.18E-05 |
| IMPA2 | -5.3770455 | 2.28E-05 |
| LTK | -8.1012045 | 2.31E-05 |
| FAM172BP | -5.0424015 | 2.76E-05 |
| LL22NC03-75 | -7.9222901 | 2.76E-05 |
| EEPD1 | -5.2988992 | 3.42E-05 |
| GZMH | 4.79081815 | 4.79E-05 |
| RP11-693M3 | 5.11324118 | 4.93E-05 |
| RP3-467K16. | 5.04878322 | 5.09E-05 |
| RAB27B | 4.22536889 | 6.14E-05 |
| KIT | -6.525611 | 6.22E-05 |
| GPR183 | -4.0705392 | 6.25E-05 |
| PCDH9 | 4.38832345 | 6.41E-05 |
| TPBG | -6.4660944 | 7.04E-05 |
| RALGPS2 | -6.6919451 | 7.49E-05 |
| TRBV30 | -4.9034679 | 7.91E-05 |
| LAYN | -5.5121803 | 8.00E-05 |
| CCL20 | -8.7215648 | 8.74E-05 |
| GPR141 | 5.89369225 | 9.24E-05 |
| PLD1 | -5.2400003 | 9.78E-05 |
| HYAL2 | -5.2557578 | 9.85E-05 |
| RUNX2 | -5.1678412 | 0.00012268 |
| CXCL16 | -4.5795995 | 0.00013433 |
| RP11-702B1( | 4.59945076 | 0.0001345 |
| KRT1 | -4.4778648 | 0.00013611 |
| ELF3 | 4.51366361 | 0.00013862 |
| WFS1 | -3.7133727 | 0.00013895 |
| MFSD13A | 4.37149018 | 0.00014782 |

| | | |
|---|---|---|
| ZNF415 | -4.1390723 | 0.00014931 |
| PACSIN1 | -4.119432 | 0.00015351 |
| RP11-810P12 | -3.8116576 | 0.00017377 |
| NKX2-5 | 3.44842125 | 0.00019973 |
| CD28 | -5.5722587 | 0.00020323 |
| ZBTB16 | -6.9582339 | 0.00020401 |
| AQP3 | -6.2088876 | 0.00020588 |
| TNFRSF10A | -4.7456146 | 0.00020766 |
| ADGRG1 | 5.29386809 | 0.00020794 |
| CLCN5 | -3.2468096 | 0.00020928 |
| ZC4H2 | -3.4637763 | 0.0002274 |
| RP11-499E18 | 4.18962265 | 0.00023131 |
| SOCS3 | -3.8945518 | 0.00023285 |
| SPINT2 | -5.4714147 | 0.0002334 |
| ARSJ | -4.7816158 | 0.00024052 |
| GZMB | 4.96212901 | 0.00024508 |
| GS1-115G20 | 3.26400963 | 0.00025423 |
| ADTRP | -3.9984645 | 0.00026164 |
| NLRP3 | -4.531967 | 0.00026863 |
| GZMK | -4.749842 | 0.0002807 |
| LY96 | -5.5331808 | 0.00029092 |
| VSIG1 | -4.5510859 | 0.00030559 |
| PRSS23 | 7.31495417 | 0.00031447 |
| RP11-12J10.4 | 3.38037317 | 0.00032291 |
| DMKN | -4.018972 | 0.0003316 |
| CD80 | -3.6233441 | 0.00033319 |
| KRT8P33 | 3.12560207 | 0.00035766 |
| PRSS35 | -6.9359442 | 0.00039286 |
| IFNG-AS1 | -4.937554 | 0.00042525 |
| RP11-843B15 | 3.42071479 | 0.00042677 |
| FBXW9 | -2.9912166 | 0.00042767 |
| RASSF4 | 5.61418728 | 0.00042834 |
| CTB-25B13.5 | -3.9547988 | 0.00045381 |
| DSCC1 | 4.3447614 | 0.00045828 |
| TMEM171 | -5.4094518 | 0.00048719 |
| TLE1 | -5.3503242 | 0.00049892 |
| RP11-288K12 | -3.411368 | 0.00051446 |
| ITGAM | 3.45457773 | 0.00052757 |
| SYDE2 | -4.7564282 | 0.00054672 |
| TET1 | 4.02852054 | 0.00058112 |
| CCR7 | -5.5252266 | 0.00059859 |
| HMGB3P22 | -3.4830156 | 0.0006624 |
| BTBD11 | -2.522421 | 0.0006739 |
| PDZD4 | 3.22284502 | 0.00067565 |
| CMTM8 | -4.5261509 | 0.00068098 |
| RP11-381K20 | 3.03911489 | 0.00068124 |

| | | |
|---|---|---|
| TRAT1 | -3.0230292 | 0.00069085 |
| MFSD2A | -2.776359 | 0.00069669 |
| RP11-84A14. | 2.80918439 | 0.00071455 |
| NET1 | -3.6089808 | 0.00074622 |
| CPNE7 | -3.7450675 | 0.00074786 |
| MEGF6 | -4.1914663 | 0.00075681 |
| RRP12 | -3.4322386 | 0.00077469 |
| NUAK1 | 5.69329114 | 0.00078596 |
| BACH2 | -4.3109871 | 0.00081909 |
| SVIL | -2.5475316 | 0.00084288 |
| APP | -4.9479168 | 0.00090191 |
| ANO9 | -6.2899002 | 0.00090943 |
| RP11-324I22 | 4.53804244 | 0.00092863 |
| CCR6 | -4.684753 | 0.00093186 |
| RP11-480A1( | -2.6530162 | 0.00093328 |
| PCDHGA6 | 3.68014718 | 0.00095944 |
| SPINK2 | -4.7039275 | 0.00096609 |
| RP11-565F1( | -3.2323462 | 0.00096892 |
| AXIN2 | -4.3969129 | 0.00099006 |
| NMUR1 | 3.86486802 | 0.00099487 |
| CMKLR1 | 5.4271651 | 0.00099971 |
| TFCP2L1 | 4.9185596 | 0.0010014 |
| RP11-18H21. | -6.3911538 | 0.00102753 |
| RP11-404F1( | -3.5163184 | 0.00104555 |
| DKK3 | -5.9064193 | 0.00104831 |
| CTC-366B18. | -4.4735382 | 0.00105627 |
| RP11-730B2. | -2.7198237 | 0.00106789 |
| RP11-474B1( | 3.14662405 | 0.00108304 |
| RP11-627G2. | 4.75162392 | 0.00110315 |
| AC002331.1 | -3.2277339 | 0.00113378 |
| RAMP1 | 5.68445748 | 0.00113643 |
| FCGR3A | 4.36747399 | 0.00114158 |
| ID3 | -6.5855813 | 0.00116182 |
| GRASP | -3.351809 | 0.0011741 |
| DDR2 | 3.13640456 | 0.00117989 |
| RP11-83A24. | -3.7815522 | 0.00118121 |
| CAMK4 | -2.3772183 | 0.00119524 |
| DPP4 | -6.9149683 | 0.0012011 |
| CEBPD | -5.857237 | 0.00123565 |
| FLT4 | -5.656313 | 0.00127001 |
| PTPN12 | 2.25791586 | 0.00132189 |
| IGFBP4 | -4.9293598 | 0.00132367 |
| ANK3 | -2.5743565 | 0.00133978 |
| AKR1C4 | 2.29368895 | 0.0013624 |
| RP11-493E1. | 3.62530373 | 0.00136299 |
| SPRY1 | -4.7813225 | 0.00137711 |

| | | |
|---|---|---|
| TCEB1P31 | -2.340077 | 0.00139463 |
| RASGRF2 | -3.7899746 | 0.00139662 |
| RAP1GAP2 | 4.78204299 | 0.00142207 |
| PASK | -2.4900948 | 0.00143096 |
| GNRHR2 | -2.3802079 | 0.00155245 |
| SGPP2 | -4.5801223 | 0.00159124 |
| TMPRSS6 | 3.28225628 | 0.00161535 |
| LINC01358 | 3.61552106 | 0.00168653 |
| TNFSF13B | -3.1159368 | 0.00176931 |
| LRRC16B | 4.34644753 | 0.00181087 |
| TBX21 | 2.2052919 | 0.00182449 |
| TOP2A | -4.0426144 | 0.00182845 |
| TSHZ3 | 5.32311116 | 0.00184249 |
| NEFH | 2.9816944 | 0.00184698 |
| AMPD3 | -4.4145164 | 0.0018563 |
| AGPAT4 | 2.549077 | 0.00190754 |
| KLRB1 | -2.8937912 | 0.00194545 |
| GPAT3 | -5.9450181 | 0.00195377 |
| NCF4 | -3.4935337 | 0.00195841 |
| SLC35F2 | -2.9284755 | 0.00197912 |
| ROM1 | -3.1288493 | 0.00205506 |
| HAUS7 | -2.969126 | 0.00206049 |
| FRMD5 | 3.80732913 | 0.00207625 |
| LINC01146 | -3.823781 | 0.00209399 |
| CDH7 | 3.34395248 | 0.00210471 |
| BCL2L15 | 2.79083678 | 0.00210677 |
| MATN2 | -4.7190922 | 0.0021422 |
| VAV2 | -2.7328885 | 0.00215999 |
| NCR3LG1 | -3.168312 | 0.0021801 |
| ZSWIM1 | 3.94374962 | 0.00222854 |
| BATF3 | -3.3908151 | 0.00224176 |
| THRB | 2.41706164 | 0.0022617 |
| SRPK2P | 2.38345597 | 0.00235043 |
| RP1-244F24. | -4.0155916 | 0.00246879 |
| SPTB | -3.2651265 | 0.00256542 |
| CIART | -4.6912295 | 0.00260443 |
| SYNPO2 | -2.4876271 | 0.00265033 |
| GMPR | -4.0562584 | 0.00265778 |
| OSMR | 3.93330279 | 0.0026691 |
| ZEB1 | -2.5388456 | 0.00275231 |
| MBOAT2 | 3.45527593 | 0.00277145 |
| FXYD2 | -4.293337 | 0.00280034 |
| IL6R | -4.14917 | 0.00281398 |
| LINC01393 | -2.8528745 | 0.00282006 |
| FRMPD3 | 3.75680373 | 0.00283197 |
| RP11-815J21 | 3.4887746 | 0.00291531 |

| | | |
|---|---|---|
| PRCD | -4.3557022 | 0.00291676 |
| ZNF385D | 2.34597977 | 0.00298269 |
| RP11-787I22 | 3.06698383 | 0.0030078 |
| PEX10 | -2.808477 | 0.0030378 |
| IMMP1LP2 | -2.1297667 | 0.00309573 |
| C4orf32 | -2.7495121 | 0.00310405 |
| ME3 | 3.08851732 | 0.00310796 |
| RP11-967K21 | -3.4369082 | 0.00312531 |
| ZFP28 | -5.1889801 | 0.00318558 |
| FURIN | -2.0332024 | 0.00319442 |
| ITPKA | -3.4173359 | 0.00326677 |
| RP11-188C12 | 1.96073434 | 0.00328505 |
| HIVEP3 | 3.09334934 | 0.00328511 |
| OR7E13P | 2.29737006 | 0.00329826 |
| ACTN1 | -2.7467736 | 0.00341665 |
| LGALS3 | -2.7271035 | 0.00344282 |
| RPS6KA6 | 3.33457952 | 0.00344329 |
| PAG1 | -2.5799376 | 0.00346088 |
| CXCR6 | -3.1346509 | 0.00348369 |
| DYX1C1 | -2.7367402 | 0.00350117 |
| TPRN | 2.15602584 | 0.00351636 |
| RP11-81H14. | 6.03722311 | 0.00355757 |
| ZNF814 | -3.9510391 | 0.00363409 |
| RP3-426I6.6 | -3.1260077 | 0.00364339 |
| DENND6B | -3.589839 | 0.00369194 |
| ZNF683 | 4.66190384 | 0.00369611 |
| RP5-902P8.1 | 4.43084803 | 0.0037071 |
| ABLIM2 | 5.03205949 | 0.00376084 |
| RP1-40E16.1 | 3.62144385 | 0.00376267 |
| ZNF793-AS1 | 3.46681327 | 0.00380811 |
| RP11-25I15.1 | 2.19138482 | 0.00382815 |
| RP5-915N17. | -1.8143562 | 0.00383964 |
| CHN1 | -3.0890249 | 0.00386622 |
| MN1 | 3.62901668 | 0.00388252 |
| ATF7IP2 | -2.2547933 | 0.00391348 |
| CLEC18B | 2.91161733 | 0.0039192 |
| KCNQ3 | -3.2705515 | 0.00392854 |
| IL2RA | -3.7122586 | 0.00394799 |
| RAB40B | -2.4491635 | 0.0039865 |
| LINC00330 | 2.94032854 | 0.00398839 |
| RP3-337H4.1 | -4.5276718 | 0.00403653 |
| ZSCAN18 | -3.6775048 | 0.0041089 |
| FMR1-AS1 | 3.73572945 | 0.0041139 |
| GTF2F2P1 | -2.1782393 | 0.00412759 |
| LINC00854 | 2.92933938 | 0.00413771 |
| MME | -3.1401241 | 0.00416943 |

| | | |
|---|---|---|
| HNRNPA3P2 | -3.2571906 | 0.00417667 |
| RGS2 | -2.074225 | 0.0043431 |
| TNFRSF25 | -2.9184072 | 0.0043978 |
| C2CD4A | -3.4137321 | 0.00439823 |
| KCTD11 | 3.30623677 | 0.00444518 |
| CCDC64 | -2.7752299 | 0.00445537 |
| MYC | -5.5815082 | 0.00455119 |
| AC097711.1 | -1.8080436 | 0.00462315 |
| RP11-564D1: | 2.68514885 | 0.00474072 |
| MYO3B | 4.2618944 | 0.00475091 |
| MORN1 | 3.10374187 | 0.00476992 |
| KLRD1 | 2.04432729 | 0.00477868 |
| WDR86-AS1 | -5.4800443 | 0.0047825 |
| AL450992.2 | -3.3020438 | 0.0047939 |
| ZNF584 | -2.7159901 | 0.00480616 |
| CA2 | -7.2128216 | 0.00482297 |
| CPNE2 | -5.6646001 | 0.0048471 |
| RPA4 | 3.74298683 | 0.00487377 |
| JAML | -2.9445514 | 0.00487668 |
| CX3CR1 | 5.65004058 | 0.00495047 |
| RP1-228P16. | 3.01555505 | 0.00497622 |
| B3GALT2 | -4.9518981 | 0.00501693 |
| FCRL6 | 2.53977205 | 0.00502109 |
| ALCAM | -5.1002655 | 0.00507484 |
| RP11-589C21 | -2.4882876 | 0.00516058 |
| SATB1 | -1.7842241 | 0.00525082 |
| PIK3C2B | -2.9495125 | 0.00527623 |
| EFNB1 | 3.3919422 | 0.00533801 |
| SCD5 | 2.06068875 | 0.00534201 |
| USP12PX | -2.5545224 | 0.00534623 |
| CPPED1 | 2.93893131 | 0.00535849 |
| FBXL16 | -4.1504741 | 0.00539131 |
| TSPY26P | -3.6298285 | 0.00539542 |
| SATB2 | 3.62495798 | 0.00549059 |
| ADCY3 | -3.0568591 | 0.00556359 |
| CDC27P2 | 1.68225004 | 0.00560148 |
| GINS3 | -1.7025991 | 0.00571566 |
| SNORD3B-2 | -2.5588354 | 0.00577676 |
| CEP126 | 2.14827559 | 0.00581026 |
| CRIP1P4 | 1.86042211 | 0.00582198 |
| TYMS | -2.50303 | 0.00586755 |
| CTA-212A2.1 | -4.522627 | 0.00589672 |
| ICE2P2 | -1.9469821 | 0.00590755 |
| RPSAP21 | 3.33874249 | 0.00594978 |
| ZNF532 | -3.5320243 | 0.00595244 |
| ZNF93 | -4.3955921 | 0.00598846 |

| | | |
|---|---|---|
| ST3GAL3 | -2.9654238 | 0.00600674 |
| IL4I1 | -3.7734531 | 0.00603098 |
| SLC29A4 | 3.39938854 | 0.00605084 |
| RRS1 | -3.2187656 | 0.00612741 |
| PIANP | -3.4956381 | 0.00617042 |
| RASSF2 | -2.3717416 | 0.00623401 |
| TFP1 | 1.95064959 | 0.00624863 |
| NEDD4L | -3.7418405 | 0.00627979 |
| ENPP1 | -3.9054041 | 0.00631245 |
| TAF4B | -3.4321276 | 0.00636333 |
| TNFSF8 | -4.6312802 | 0.00637213 |
| TMEM164 | -1.5819369 | 0.00639481 |
| AC013264.2 | -4.9711061 | 0.00648803 |
| RP11-574K11 | -3.4316091 | 0.00655549 |
| KRT72 | 4.19004782 | 0.00657348 |
| RP11-817I4.1 | -2.3411667 | 0.00667419 |
| AC007191.4 | 3.06122953 | 0.00668504 |
| AE000662.93 | -2.1012157 | 0.00677538 |
| FOS | -5.1382429 | 0.00679162 |
| RPL23AP96 | 1.80553328 | 0.00682241 |
| TCF4 | 2.53805467 | 0.00682797 |
| TYROBP | 2.74365657 | 0.00683094 |
| RP1-12G14.7 | 1.95418948 | 0.00685732 |
| RAB11FIP5 | 4.27851157 | 0.00709113 |
| ZEB2 | 2.04676035 | 0.00709166 |
| CEP170B | -2.967793 | 0.00715175 |
| CEP164P1 | 2.67198479 | 0.00717181 |
| NKG7 | 1.95534514 | 0.00720465 |
| SLC7A6 | -1.8995468 | 0.0072093 |
| RP11-552M1 | -1.8865334 | 0.00721583 |
| CTC-523E23.4 | 4.00998182 | 0.00723195 |
| C17orf51 | 2.90421378 | 0.00737693 |
| RP11-301G2 | -1.7009936 | 0.00746487 |
| RP11-388C12 | -3.4674134 | 0.00746872 |
| A1BG | -1.8474057 | 0.00750988 |
| CH17-373J23 | -2.4178088 | 0.00751594 |
| CTD-2313F11 | -3.937087 | 0.00752841 |
| LSMEM1 | 2.98948148 | 0.00752994 |
| RP11-302I18 | -1.9242139 | 0.00754208 |
| MTSS1 | 2.19278145 | 0.00759507 |
| RP11-1143G9 | -3.4680537 | 0.0076234 |
| MIR3142HG | 3.55946219 | 0.00762503 |
| TSPAN2 | 3.74620213 | 0.00766378 |
| CTD-2576D5. | -2.910464 | 0.00775893 |
| SPON2 | 1.77491869 | 0.00783662 |
| USP45 | -1.6291106 | 0.00785723 |

| | | |
|---|---|---|
| COL6A1 | -3.6897329 | 0.00791647 |
| MPZL3 | -2.5101362 | 0.00794617 |
| AJ003147.11 | 3.01991407 | 0.00797908 |
| RP11-358B2: | -4.0458024 | 0.0079902 |
| FAM161B | 3.90885874 | 0.00806594 |
| LBX2 | -3.0606945 | 0.00812883 |
| LINC01215 | -3.7839619 | 0.00819436 |
| CST3 | 4.18987955 | 0.00822147 |
| BEX1 | -4.031016 | 0.00822158 |
| ZNF629 | -2.45953 | 0.00824193 |
| CYSLTR1 | -5.3391214 | 0.00824488 |
| ZNF883 | 4.5701447 | 0.00830706 |
| SLC1A7 | 4.08073127 | 0.00830828 |
| RP11-297C4. | 1.87414214 | 0.00838824 |
| GPR135 | 3.39020436 | 0.00840111 |
| AC009784.3 | 3.55104896 | 0.00844124 |
| KCTD21-AS1 | -2.6128015 | 0.00844543 |
| BEX5 | -2.3358143 | 0.00846803 |
| NEK10 | 2.90537106 | 0.00848249 |
| AC009404.2 | 1.73937221 | 0.00849655 |
| TNFRSF13C | -2.9679338 | 0.00855947 |
| TRIM62 | -2.3864216 | 0.00865072 |
| RP11-160H2: | 3.42154666 | 0.00866287 |
| PHBP9 | -2.1648607 | 0.00874044 |
| FGR | 3.26222287 | 0.00877546 |
| PGAP3 | 3.06201168 | 0.00878008 |
| ITGB2-AS1 | 2.08362913 | 0.00880205 |
| AC083862.6 | -1.9407933 | 0.00886363 |
| RP11-1102P: | 2.16053141 | 0.00892997 |
| CTD-2619J13 | 2.78059864 | 0.00895465 |
| KDSR | -2.3683801 | 0.00895923 |
| ZNF165 | -4.3564758 | 0.00902325 |
| LZTS1 | -2.2877424 | 0.00911499 |
| PARP3 | 2.26727431 | 0.00911707 |
| ELOVL4 | -4.1854975 | 0.0092578 |
| PCAT29 | -2.7836273 | 0.00927608 |
| EFNA5 | 4.46933797 | 0.00928734 |
| KLRAP1 | 1.90545838 | 0.0093413 |
| AP000240.9 | -3.284644 | 0.00939413 |
| IL18RAP | -2.020057 | 0.00941867 |
| CD27 | -6.0291806 | 0.00946658 |
| CHRNA5 | -2.9973767 | 0.009523 |
| KIAA0319L | -1.6902763 | 0.00953418 |
| AC007256.5 | 3.4104647 | 0.00958541 |
| GNPDA1 | -2.1760558 | 0.0095995 |
| TNFSF9 | -4.6851506 | 0.00960888 |

| | | |
|---|---|---|
| RLN2 | -2.0262612 | 0.00964284 |
| C9orf24 | 2.30564423 | 0.00966205 |
| AMH | 3.14921926 | 0.0096878 |
| FSD1L | 3.80421976 | 0.00973019 |
| CTD-2301A4. | 2.39193898 | 0.00973553 |
| LIM2 | 3.80818392 | 0.00973972 |
| RP5-864K19. | 2.53101966 | 0.00979053 |
| NPDC1 | -2.989246 | 0.00982789 |
| GNLY | 2.61791211 | 0.00982928 |
| SATB1-AS1 | -2.4348543 | 0.01010033 |
| DUSP19 | 3.26890606 | 0.01017099 |
| XXbac-BPGBI | 2.42963168 | 0.01021572 |
| POU5F2 | 2.53842233 | 0.01021917 |
| PRRG3 | 3.35142932 | 0.01022753 |
| BEX2 | -1.7253744 | 0.01023094 |
| ACTG1P20 | -1.9659736 | 0.01023364 |
| RP11-665C16 | 1.54149845 | 0.01028663 |
| SLC35F6 | -3.7070209 | 0.01031703 |
| HOOK1 | -5.2173412 | 0.01033939 |
| PDGFB | -3.3799464 | 0.01034431 |
| CASK | -2.1564825 | 0.01036205 |
| MICAL2 | -2.596299 | 0.01042025 |
| AC087501.1 | 1.8817847 | 0.01050901 |
| TUBA3FP | 3.0355814 | 0.01056178 |
| TTC39C-AS1 | -2.0557765 | 0.01059898 |
| GALM | -1.8598924 | 0.01062307 |
| RP11-415F23 | -3.3736352 | 0.01080889 |
| SNORA66 | 1.91071938 | 0.01087774 |
| CTD-2537I9.1 | -1.5042772 | 0.01089824 |
| LINC01012 | 2.84439527 | 0.01101823 |
| RP11-521C22 | -4.1798667 | 0.01113673 |
| TGFBR3L | 2.16435315 | 0.01119463 |
| RPL7L1P9 | 3.33823086 | 0.01121218 |
| EPHX4 | 2.2920744 | 0.01130002 |
| PLEKHG2 | -2.6774617 | 0.01133462 |
| LGALS1 | 1.93424679 | 0.01133494 |
| TCF7 | -1.966142 | 0.01135556 |
| RPL21P38 | 1.89415933 | 0.01136253 |
| ATP11A-AS1 | -2.6438185 | 0.01152211 |
| RP11-529G2 | 1.86883743 | 0.01152977 |
| ORC6 | 3.24917209 | 0.01153982 |
| MAP6D1 | -3.6731213 | 0.01154465 |
| DUSP1 | -2.6697002 | 0.01159296 |
| SLAMF1 | -2.2285955 | 0.01162402 |
| ARHGEF39 | -1.5501045 | 0.01165539 |
| AC006116.12 | -1.8890273 | 0.01185514 |

| | | |
|---|---|---|
| COL27A1 | -4.0763379 | 0.0119223 |
| AC004067.5 | -3.8603711 | 0.01192884 |
| PDE7B | -3.610391 | 0.01195776 |
| RN7SL448P | 1.95203089 | 0.01209123 |
| MYO15A | -3.6248181 | 0.01215077 |
| C10orf35 | -4.7460273 | 0.01217113 |
| EFHD2 | 1.41286767 | 0.01217742 |
| RP11-28G8.1 | 3.19420105 | 0.01226285 |
| MIR181A1HG | -1.7660875 | 0.012321 |
| RP11-473O4. | 2.29850511 | 0.01235451 |
| RP11-25K19. | 1.69652417 | 0.01236554 |
| MAST4 | -2.0218566 | 0.01248308 |
| SPATA6 | 3.95180198 | 0.01249149 |
| COTL1 | -1.6854518 | 0.01249241 |
| RP11-996F15 | -4.2957332 | 0.0124978 |
| RP11-617F23 | 3.0414199 | 0.01253654 |
| RP11-295H24 | 1.56680838 | 0.01254205 |
| LANCL1-AS1 | 2.18933849 | 0.01266414 |
| RARB | 4.3235488 | 0.01267426 |
| TRAV8-6 | -3.9458761 | 0.0127096 |
| IL7R | -1.7581975 | 0.01271333 |
| CEACAM8 | 2.40891164 | 0.01274702 |
| AC025750.5 | -2.02583 | 0.01298423 |
| MCM3AP-AS | 1.73712656 | 0.0130123 |
| KANTR | -2.2332072 | 0.01304004 |
| RP11-490B18 | -1.7537764 | 0.01311112 |
| ERI2 | 3.80881688 | 0.01314436 |
| MYOT | 2.33227555 | 0.0132122 |
| SCART1 | -3.1078683 | 0.01328655 |
| FOXO4 | 2.77862794 | 0.01329571 |
| MFAP3L | 3.04723917 | 0.0134514 |
| RP11-5O23.1 | 2.08949103 | 0.01347617 |
| RP11-354M2 | 2.1466854 | 0.01361058 |
| RP11-106D4. | -3.2560666 | 0.01361355 |
| RAB37 | 2.66162518 | 0.01367585 |
| LINC00899 | -2.1339936 | 0.01368994 |
| ZNF850 | 4.11963994 | 0.01373509 |
| RP11-478C1. | 2.19181345 | 0.01373789 |
| CTB-50L17.8 | 2.62380466 | 0.01375955 |
| ST8SIA6 | 1.77230741 | 0.01381283 |
| DUSP16 | -1.8744474 | 0.01383269 |
| RP11-16P6.1 | -2.3648972 | 0.01388656 |
| SKOR1 | 2.0340539 | 0.01399497 |
| CDH2 | -2.7802014 | 0.01405768 |
| FSD1 | -3.3253422 | 0.01406919 |
| LL0XNC01-24 | -1.72032 | 0.01410271 |

| | | |
|---|---|---|
| ZDBF2 | -5.211459 | 0.01410602 |
| FCGR2A | 2.95021798 | 0.01416447 |
| ATP6V0E1P3 | 1.77870282 | 0.01419579 |
| GPR171 | -1.870198 | 0.01420336 |
| SPOCK2 | -1.451072 | 0.01427033 |
| RP11-4O1.2 | 3.67019377 | 0.01430995 |
| FIGNL1 | 1.65171486 | 0.01440192 |
| CELSR1 | -2.9463459 | 0.01444128 |
| RP11-367E12 | 2.20194398 | 0.0144673 |
| CTD-2349P21 | 1.91781083 | 0.01452287 |
| RP11-500G2 | -2.6996688 | 0.01453888 |
| ZGRF1 | 1.78572404 | 0.01457906 |
| SSBP3 | 1.89997545 | 0.01463155 |
| RHEBP2 | 1.68899902 | 0.01464203 |
| DAPP1 | -3.9109923 | 0.01469421 |
| RP11-73G16. | 1.48767209 | 0.01472694 |
| C1orf21 | 1.52416192 | 0.01479505 |
| RP11-363E7. | -2.4559635 | 0.01483814 |
| ACVR1B | -3.6021616 | 0.01484776 |
| RP11-342K2. | 3.18140446 | 0.0149149 |
| SAMD12 | -4.5535456 | 0.01504595 |
| ADCY1 | 2.19800466 | 0.015242 |
| AC006042.8 | 3.0937057 | 0.01525893 |
| CTD-2290C23 | -1.5106182 | 0.01526347 |
| RP11-359K18 | 2.47729633 | 0.01529259 |
| TBC1D31 | -1.5002827 | 0.01532233 |
| NPIPB4 | 1.81765564 | 0.01536603 |
| POC1A | 2.02922451 | 0.01540211 |
| RP11-432I5.8 | 1.61636816 | 0.01543943 |
| MRPL4 | -1.6155363 | 0.01546725 |
| RP3-405J10.3 | 2.85625265 | 0.01554045 |
| SHROOM1 | 2.29841562 | 0.01562039 |
| SNED1 | -4.0314367 | 0.01563917 |
| NUAK2 | 2.16843405 | 0.01573615 |
| RTTN | 1.54749669 | 0.01579089 |
| XBP1 | -1.8615348 | 0.0158768 |
| MS4A1 | -5.6002533 | 0.01591872 |
| FTH1P3 | -1.5375305 | 0.0159224 |
| KIF13A | 2.91448765 | 0.0159309 |
| NLE1 | -2.5569774 | 0.01594003 |

**Table S5. List of top 500 ATAC peaks higher in R1+27-**

| peak_id | chr | start | end | gene | distTSS | log2FoldChar | pvalue |
|---|---|---|---|---|---|---|---|
| Peak69990 | chr4 | 105818202 | 105818420 | CXXC4 | -402260 | -3.0085959 | 8.38E-11 |
| Peak69990 | chr4 | 105818202 | 105818420 | TET2 | -249139 | -3.0085959 | 8.38E-11 |
| Peak24237 | chr12 | 124226838 | 124227082 | DNAH10 | -20082 | -2.3094131 | 3.26E-10 |
| Peak24237 | chr12 | 124226838 | 124227082 | ATP6V0A2 | 30095 | -2.3094131 | 3.26E-10 |
| Peak26364 | chr13 | 76063972 | 76064428 | TBC1D4 | -7950 | -1.3120433 | 6.13E-10 |
| Peak26364 | chr13 | 76063972 | 76064428 | COMMD6 | 47777 | -1.3120433 | 6.13E-10 |
| Peak48430 | chr2 | 38412252 | 38412513 | CYP1B1 | -109060 | -2.391483 | 9.87E-09 |
| Peak48430 | chr2 | 38412252 | 38412513 | ATL2 | 192021 | -2.391483 | 9.87E-09 |
| Peak80990 | chr6 | 102293552 | 102293863 | GRIK2 | 447044 | -1.5400576 | 1.19E-08 |
| Peak64187 | chr3 | 112156527 | 112156705 | BTLA | 61792 | -1.1278744 | 1.24E-08 |
| Peak64187 | chr3 | 112156527 | 112156705 | CD200 | 104613 | -1.1278744 | 1.24E-08 |
| Peak82184 | chr6 | 132933675 | 132933895 | TAAR5 | -22908 | -2.3329349 | 1.54E-08 |
| Peak82184 | chr6 | 132933675 | 132933895 | TAAR2 | 11629 | -2.3329349 | 1.54E-08 |
| Peak34057 | chr16 | 4166424 | 4166694 | ADCY9 | -373 | -1.4312396 | 4.19E-08 |
| Peak68001 | chr4 | 25689424 | 25689744 | SLC34A2 | 32118 | -1.8877738 | 4.73E-08 |
| Peak68001 | chr4 | 25689424 | 25689744 | SEL1L3 | 174997 | -1.8877738 | 4.73E-08 |
| Peak52869 | chr2 | 176072613 | 176072768 | ATP5G3 | -23356 | -2.4567246 | 8.09E-08 |
| Peak69989 | chr4 | 105817905 | 105818076 | CXXC4 | -401940 | -2.8632136 | 1.14E-07 |
| Peak69989 | chr4 | 105817905 | 105818076 | TET2 | -249459 | -2.8632136 | 1.14E-07 |
| Peak47250 | chr2 | 7139356 | 7139525 | RNF144A | 81918 | -2.6924173 | 1.18E-07 |
| Peak2797 | chr1 | 64358676 | 64358991 | UBE2U | -310656 | -2.5119187 | 1.47E-07 |
| Peak2797 | chr1 | 64358676 | 64358991 | ROR1 | 119120 | -2.5119187 | 1.47E-07 |
| Peak94915 | chr9 | 4851731 | 4851943 | JAK2 | -133408 | -2.6713128 | 1.93E-07 |
| Peak94915 | chr9 | 4851731 | 4851943 | RCL1 | 58968 | -2.6713128 | 1.93E-07 |
| Peak39909 | chr17 | 59883480 | 59883783 | NACA2 | -215069 | -1.8779387 | 1.94E-07 |
| Peak39909 | chr17 | 59883480 | 59883783 | BRIP1 | 57250 | -1.8779387 | 1.94E-07 |
| Peak61474 | chr3 | 25502225 | 25502593 | RARB | 32607 | -2.7439556 | 2.08E-07 |
| Peak61474 | chr3 | 25502225 | 25502593 | TOP2B | 203581 | -2.7439556 | 2.08E-07 |
| Peak12943 | chr10 | 97055023 | 97055310 | PDLIM1 | -4386 | -1.4107295 | 2.65E-07 |
| Peak80356 | chr6 | 76236340 | 76236521 | TMEM30A | -241747 | -1.1815861 | 2.65E-07 |
| Peak80356 | chr6 | 76236340 | 76236521 | SENP6 | -75332 | -1.1815861 | 2.65E-07 |
| Peak92009 | chr8 | 60095340 | 60095599 | TOX | -63703 | -2.1862944 | 2.86E-07 |
| Peak3057 | chr1 | 75045137 | 75045474 | TYW3 | -153534 | -2.0336401 | 2.88E-07 |
| Peak3057 | chr1 | 75045137 | 75045474 | LRRC53 | -67008 | -2.0336401 | 2.88E-07 |
| Peak95223 | chr9 | 16884931 | 16885274 | CNTLN | -249877 | -2.6792943 | 3.25E-07 |
| Peak95223 | chr9 | 16884931 | 16885274 | BNC2 | -14399 | -2.6792943 | 3.25E-07 |
| Peak14762 | chr11 | 6249865 | 6250063 | FAM160A2 | 5977 | -1.5426242 | 3.50E-07 |
| Peak14762 | chr11 | 6249865 | 6250063 | OR52W1 | 29588 | -1.5426242 | 3.50E-07 |
| Peak22664 | chr12 | 90130857 | 90131045 | ATP2B1 | -28343 | -2.5298098 | 3.71E-07 |
| Peak51450 | chr2 | 131323841 | 131324040 | ENSG00000 | -4518 | -2.2500938 | 4.77E-07 |
| Peak52879 | chr2 | 176440353 | 176440829 | ATP5G3 | -391256 | -2.6354136 | 4.93E-07 |
| Peak23746 | chr12 | 113286978 | 113287133 | OAS1 | -57532 | -2.4571099 | 4.94E-07 |
| Peak23746 | chr12 | 113286978 | 113287133 | RPH3A | 57596 | -2.4571099 | 4.94E-07 |
| Peak80989 | chr6 | 102293174 | 102293479 | GRIK2 | 446663 | -1.565955 | 4.95E-07 |
| Peak71562 | chr4 | 169828596 | 169828756 | CBR4 | 102733 | -2.159648 | 5.46E-07 |
| Peak71562 | chr4 | 169828596 | 169828756 | PALLD | 410421 | -2.159648 | 5.46E-07 |

| | | | | | | | |
|---|---|---|---|---|---|---|---|
| Peak17605 | chr11 | 86924568 | 86924781 | TMEM135 | 175626 | -1.5211883 | 6.73E-07 |
| Peak17605 | chr11 | 86924568 | 86924781 | RAB38 | 983960 | -1.5211883 | 6.73E-07 |
| Peak42627 | chr18 | 55248562 | 55248781 | FECH | 5214 | -1.6170094 | 6.97E-07 |
| Peak42627 | chr18 | 55248562 | 55248781 | ONECUT2 | 145755 | -1.6170094 | 6.97E-07 |
| Peak28292 | chr14 | 52063262 | 52063542 | FRMD6 | -55263 | -2.2768092 | 7.12E-07 |
| Peak28292 | chr14 | 52063262 | 52063542 | TMX1 | 356516 | -2.2768092 | 7.12E-07 |
| Peak69848 | chr4 | 102256167 | 102256494 | EMCN | -817081 | -1.7026358 | 7.53E-07 |
| Peak69848 | chr4 | 102256167 | 102256494 | PPP3CA | 12306 | -1.7026358 | 7.53E-07 |
| Peak57378 | chr20 | 56884607 | 56884799 | PPP4R1L | -208 | -1.161829 | 8.40E-07 |
| Peak57378 | chr20 | 56884607 | 56884799 | RAB22A | -49 | -1.161829 | 8.40E-07 |
| Peak71550 | chr4 | 169555562 | 169555705 | PALLD | 137379 | -2.7492429 | 8.77E-07 |
| Peak71550 | chr4 | 169555562 | 169555705 | CBR4 | 375775 | -2.7492429 | 8.77E-07 |
| Peak41432 | chr18 | 8601697 | 8602027 | RAB12 | -7581 | -1.7312686 | 8.83E-07 |
| Peak25744 | chr13 | 47300583 | 47300752 | LCP1 | -515185 | -2.0654722 | 8.91E-07 |
| Peak25744 | chr13 | 47300583 | 47300752 | ESD | 70699 | -2.0654722 | 8.91E-07 |
| Peak31254 | chr15 | 35774220 | 35774463 | ZNF770 | -493854 | -1.9225103 | 9.20E-07 |
| Peak31254 | chr15 | 35774220 | 35774463 | DPH6 | 64027 | -1.9225103 | 9.20E-07 |
| Peak95221 | chr9 | 16871767 | 16872130 | BNC2 | -1245 | -2.2807147 | 9.65E-07 |
| Peak62966 | chr3 | 60050469 | 60050804 | NA | NA | -1.0526573 | 1.06E-06 |
| Peak94415 | chr8 | 142163012 | 142163148 | DENND3 | 24360 | -2.207698 | 1.19E-06 |
| Peak94415 | chr8 | 142163012 | 142163148 | SLC45A4 | 75593 | -2.207698 | 1.19E-06 |
| Peak31772 | chr15 | 50295972 | 50296200 | ATP8B4 | 115333 | -1.2910368 | 1.26E-06 |
| Peak31772 | chr15 | 50295972 | 50296200 | FGF7 | 580793 | -1.2910368 | 1.26E-06 |
| Peak82331 | chr6 | 135853193 | 135853485 | PDE7B | -319495 | -1.7809995 | 1.27E-06 |
| Peak82331 | chr6 | 135853193 | 135853485 | AHI1 | -34456 | -1.7809995 | 1.27E-06 |
| Peak46788 | chr19 | 55206501 | 55206701 | KIR3DL3 | -29383 | -2.4643737 | 1.32E-06 |
| Peak46788 | chr19 | 55206501 | 55206701 | LILRB4 | 33022 | -2.4643737 | 1.32E-06 |
| Peak52537 | chr2 | 169235536 | 169235854 | STK39 | -131044 | -1.2122081 | 1.48E-06 |
| Peak52537 | chr2 | 169235536 | 169235854 | CERS6 | -77064 | -1.2122081 | 1.48E-06 |
| Peak12995 | chr10 | 98230028 | 98230383 | OPALIN | -111147 | -1.8610544 | 1.53E-06 |
| Peak12995 | chr10 | 98230028 | 98230383 | TLL2 | 43462 | -1.8610544 | 1.53E-06 |
| Peak32055 | chr15 | 58906917 | 58907239 | ADAM10 | 135099 | -1.7815024 | 1.57E-06 |
| Peak32055 | chr15 | 58906917 | 58907239 | LIPC | 204310 | -1.7815024 | 1.57E-06 |
| Peak64717 | chr3 | 126326052 | 126326509 | TXNRD3 | 47717 | -1.1061925 | 1.70E-06 |
| Peak64717 | chr3 | 126326052 | 126326509 | CHST13 | 83155 | -1.1061925 | 1.70E-06 |
| Peak43150 | chr18 | 74810185 | 74810313 | MBP | 34476 | -2.5229629 | 1.85E-06 |
| Peak43150 | chr18 | 74810185 | 74810313 | ZNF236 | 274133 | -2.5229629 | 1.85E-06 |
| Peak92016 | chr8 | 60293746 | 60294002 | TOX | -262107 | -1.6393396 | 1.87E-06 |
| Peak92016 | chr8 | 60293746 | 60294002 | CA8 | 900097 | -1.6393396 | 1.87E-06 |
| Peak1111 | chr1 | 25020864 | 25021079 | CLIC4 | -50876 | -2.6715534 | 1.91E-06 |
| Peak1111 | chr1 | 25020864 | 25021079 | SRRM1 | 51469 | -2.6715534 | 1.91E-06 |
| Peak70171 | chr4 | 111075751 | 111075994 | ELOVL6 | 43936 | -2.0380669 | 2.01E-06 |
| Peak70171 | chr4 | 111075751 | 111075994 | EGF | 241826 | -2.0380669 | 2.01E-06 |
| Peak88446 | chr7 | 106452460 | 106452727 | NAMPT | -526956 | -1.5972116 | 2.02E-06 |
| Peak88446 | chr7 | 106452460 | 106452727 | PIK3CG | -53330 | -1.5972116 | 2.02E-06 |
| Peak3591 | chr1 | 91999287 | 91999482 | CDC7 | 32720 | -1.5666543 | 2.17E-06 |
| Peak3591 | chr1 | 91999287 | 91999482 | TGFBR3 | 352402 | -1.5666543 | 2.17E-06 |
| Peak73918 | chr5 | 73092344 | 73092513 | ARHGEF28 | 170446 | -2.7009679 | 2.27E-06 |

| | | | | | | | |
|---|---|---|---|---|---|---|---|
| Peak73918 | chr5 | 73092344 | 73092513 | ENC1 | 844820 | -2.7009679 | 2.27E-06 |
| Peak48323 | chr2 | 37025961 | 37026137 | VIT | 102216 | -1.7496455 | 2.37E-06 |
| Peak48323 | chr2 | 37025961 | 37026137 | STRN | 167566 | -1.7496455 | 2.37E-06 |
| Peak94919 | chr9 | 4944387 | 4944614 | JAK2 | -40744 | -1.5532735 | 2.42E-06 |
| Peak94919 | chr9 | 4944387 | 4944614 | RCL1 | 151632 | -1.5532735 | 2.42E-06 |
| Peak92043 | chr8 | 61086431 | 61086655 | CA8 | 107428 | -2.5489994 | 2.55E-06 |
| Peak70741 | chr4 | 129840547 | 129840727 | PHF17 | 109858 | -2.7130819 | 2.70E-06 |
| Peak70741 | chr4 | 129840547 | 129840727 | SCLT1 | 174125 | -2.7130819 | 2.70E-06 |
| Peak26702 | chr13 | 94965188 | 94965448 | DCT | 166618 | -1.5752331 | 2.75E-06 |
| Peak95222 | chr9 | 16879503 | 16879679 | CNTLN | -255389 | -2.6117115 | 2.83E-06 |
| Peak95222 | chr9 | 16879503 | 16879679 | BNC2 | -8887 | -2.6117115 | 2.83E-06 |
| Peak5952 | chr1 | 159770035 | 159770311 | FCRL6 | -2041 | -1.5454363 | 2.88E-06 |
| Peak17785 | chr11 | 95436327 | 95436517 | SESN3 | -472061 | -1.7414477 | 3.26E-06 |
| Peak17785 | chr11 | 95436327 | 95436517 | FAM76B | 86533 | -1.7414477 | 3.26E-06 |
| Peak94995 | chr9 | 5922276 | 5922552 | MLANA | 31612 | -1.9480319 | 3.29E-06 |
| Peak94995 | chr9 | 5922276 | 5922552 | RANBP6 | 93204 | -1.9480319 | 3.29E-06 |
| Peak46784 | chr19 | 55128513 | 55128613 | LILRB1 | -13345 | -2.8477477 | 3.32E-06 |
| Peak46784 | chr19 | 55128513 | 55128613 | LILRA1 | 23450 | -2.8477477 | 3.32E-06 |
| Peak51471 | chr2 | 131730031 | 131730394 | ARHGEF4 | 55989 | -1.4648689 | 3.52E-06 |
| Peak51471 | chr2 | 131730031 | 131730394 | FAM168B | 120788 | -1.4648689 | 3.52E-06 |
| Peak97510 | chr9 | 112919238 | 112919596 | TXN | 99503 | -1.428206 | 3.57E-06 |
| Peak97510 | chr9 | 112919238 | 112919596 | AKAP2 | 376648 | -1.428206 | 3.57E-06 |
| Peak43913 | chr19 | 6753729 | 6753955 | SH2D3A | 13757 | -1.3061796 | 3.70E-06 |
| Peak43913 | chr19 | 6753729 | 6753955 | TRIP10 | 14151 | -1.3061796 | 3.70E-06 |
| Peak2732 | chr1 | 61298074 | 61298348 | C1orf87 | -758769 | -1.8804676 | 3.70E-06 |
| Peak2732 | chr1 | 61298074 | 61298348 | NFIA | -249323 | -1.8804676 | 3.70E-06 |
| Peak83678 | chr6 | 160821521 | 160821642 | SLC22A3 | 52282 | -2.0020366 | 3.92E-06 |
| Peak83678 | chr6 | 160821521 | 160821642 | LPA | 265825 | -2.0020366 | 3.92E-06 |
| Peak14921 | chr11 | 9532459 | 9532755 | WEE1 | -62621 | -1.7454505 | 4.34E-06 |
| Peak14921 | chr11 | 9532459 | 9532755 | ZNF143 | 50095 | -1.7454505 | 4.34E-06 |
| Peak77943 | chr6 | 4915183 | 4915366 | RPP40 | 89006 | -1.9040701 | 4.67E-06 |
| Peak77943 | chr6 | 4915183 | 4915366 | CDYL | 138606 | -1.9040701 | 4.67E-06 |
| Peak69849 | chr4 | 102256555 | 102256647 | EMCN | -817351 | -1.6998203 | 4.67E-06 |
| Peak69849 | chr4 | 102256555 | 102256647 | PPP3CA | 12036 | -1.6998203 | 4.67E-06 |
| Peak31973 | chr15 | 55827734 | 55827967 | DYX1C1 | -27419 | -2.4344446 | 4.98E-06 |
| Peak31973 | chr15 | 55827734 | 55827967 | PYGO1 | 53294 | -2.4344446 | 4.98E-06 |
| Peak40889 | chr17 | 79319940 | 79320252 | ENSG00000 | -53444 | -1.133777 | 5.02E-06 |
| Peak40889 | chr17 | 79319940 | 79320252 | TMEM105 | -15622 | -1.133777 | 5.02E-06 |
| Peak598 | chr1 | 12193114 | 12193332 | TNFRSF1B | -33837 | -1.4998474 | 5.02E-06 |
| Peak598 | chr1 | 12193114 | 12193332 | TNFRSF8 | 69789 | -1.4998474 | 5.02E-06 |
| Peak53338 | chr2 | 191863871 | 191864113 | STAT1 | 14984 | -1.7633553 | 5.07E-06 |
| Peak53338 | chr2 | 191863871 | 191864113 | GLS | 118439 | -1.7633553 | 5.07E-06 |
| Peak14191 | chr10 | 129840524 | 129840841 | MKI67 | 83966 | -1.8388977 | 5.10E-06 |
| Peak14191 | chr10 | 129840524 | 129840841 | PTPRE | 135358 | -1.8388977 | 5.10E-06 |
| Peak65373 | chr3 | 143847177 | 143847391 | C3orf58 | 156644 | -1.1640028 | 5.12E-06 |
| Peak101395 | chrX | 106410464 | 106410628 | RBM41 | -48512 | -2.8923458 | 5.23E-06 |
| Peak101395 | chrX | 106410464 | 106410628 | NUP62CL | 39117 | -2.8923458 | 5.23E-06 |
| Peak50168 | chr2 | 88539299 | 88539650 | TEX37 | -284694 | -1.1706196 | 5.44E-06 |

| | | | | | | | |
|---|---|---|---|---|---|---|---|
| Peak50168 | chr2 | 88539299 | 88539650 | THNSL2 | 68496 | -1.1706196 | 5.44E-06 |
| Peak41297 | chr18 | 3248765 | 3249086 | MYL12B | -13489 | -1.1581295 | 5.44E-06 |
| Peak41297 | chr18 | 3248765 | 3249086 | MYL12A | 1398 | -1.1581295 | 5.44E-06 |
| Peak29956 | chr14 | 92994228 | 92994305 | RIN3 | 14149 | -2.0629704 | 5.64E-06 |
| Peak29956 | chr14 | 92994228 | 92994305 | LGMN | 220757 | -2.0629704 | 5.64E-06 |
| Peak99530 | chrX | 9443573 | 9443792 | TBL1X | 10635 | -1.6344389 | 5.65E-06 |
| Peak99530 | chrX | 9443573 | 9443792 | GPR143 | 290321 | -1.6344389 | 5.65E-06 |
| Peak14922 | chr11 | 9532856 | 9532955 | WEE1 | -62322 | -1.9866229 | 5.76E-06 |
| Peak14922 | chr11 | 9532856 | 9532955 | ZNF143 | 50394 | -1.9866229 | 5.76E-06 |
| Peak48324 | chr2 | 37027548 | 37027868 | VIT | 103875 | -1.250733 | 5.77E-06 |
| Peak48324 | chr2 | 37027548 | 37027868 | STRN | 165907 | -1.250733 | 5.77E-06 |
| Peak62970 | chr3 | 60146244 | 60146527 | NA | NA | -1.8727054 | 5.83E-06 |
| Peak95299 | chr9 | 20242794 | 20243216 | SLC24A2 | -456079 | -1.328191 | 6.05E-06 |
| Peak95299 | chr9 | 20242794 | 20243216 | MLLT3 | 379537 | -1.328191 | 6.05E-06 |
| Peak81838 | chr6 | 126115198 | 126115327 | HINT3 | -162664 | -1.0943347 | 6.16E-06 |
| Peak81838 | chr6 | 126115198 | 126115327 | NCOA7 | 12956 | -1.0943347 | 6.16E-06 |
| Peak22815 | chr12 | 92933889 | 92934080 | PLEKHG7 | -196326 | -1.2883176 | 6.18E-06 |
| Peak22815 | chr12 | 92933889 | 92934080 | CLLU1 | 116250 | -1.2883176 | 6.18E-06 |
| Peak12861 | chr10 | 94082983 | 94083239 | 5-Mar | 32191 | -2.6144433 | 6.18E-06 |
| Peak12861 | chr10 | 94082983 | 94083239 | IDE | 250722 | -2.6144433 | 6.18E-06 |
| Peak53028 | chr2 | 180163426 | 180163780 | SESTD1 | -34086 | -2.8047785 | 6.23E-06 |
| Peak53028 | chr2 | 180163426 | 180163780 | ZNF385B | 562629 | -2.8047785 | 6.23E-06 |
| Peak26536 | chr13 | 81052091 | 81052585 | SPRY2 | -138544 | -1.2567484 | 6.31E-06 |
| Peak42230 | chr18 | 42686325 | 42686709 | SLC14A2 | -508249 | -1.4625244 | 6.41E-06 |
| Peak42230 | chr18 | 42686325 | 42686709 | SETBP1 | 425654 | -1.4625244 | 6.41E-06 |
| Peak71180 | chr4 | 151830410 | 151830537 | LRBA | 106175 | -2.4778296 | 6.43E-06 |
| Peak71180 | chr4 | 151830410 | 151830537 | MAB21L2 | 327397 | -2.4778296 | 6.43E-06 |
| Peak80382 | chr6 | 76540640 | 76541066 | MYO6 | 81804 | -1.688167 | 6.43E-06 |
| Peak80382 | chr6 | 76540640 | 76541066 | IMPG1 | 241542 | -1.688167 | 6.43E-06 |
| Peak13682 | chr10 | 114863009 | 114863140 | HABP2 | -449710 | -2.7516573 | 6.74E-06 |
| Peak13682 | chr10 | 114863009 | 114863140 | TCF7L2 | 153066 | -2.7516573 | 6.74E-06 |
| Peak80955 | chr6 | 100676703 | 100677062 | MCHR2 | -234784 | -1.2883634 | 6.86E-06 |
| Peak80955 | chr6 | 100676703 | 100677062 | SIM1 | 235922 | -1.2883634 | 6.86E-06 |
| Peak35638 | chr16 | 57923445 | 57923634 | KIFC3 | -87040 | -2.7570643 | 6.88E-06 |
| Peak35638 | chr16 | 57923445 | 57923634 | CNGB1 | 81476 | -2.7570643 | 6.88E-06 |
| Peak28236 | chr14 | 50988697 | 50988812 | CDKL1 | -126138 | -1.2179243 | 6.96E-06 |
| Peak28236 | chr14 | 50988697 | 50988812 | MAP4K5 | 10572 | -1.2179243 | 6.96E-06 |
| Peak36602 | chr16 | 86050141 | 86050213 | FOXF1 | -493956 | -2.4881931 | 7.01E-06 |
| Peak36602 | chr16 | 86050141 | 86050213 | IRF8 | 117768 | -2.4881931 | 7.01E-06 |
| Peak87750 | chr7 | 91942455 | 91942797 | GATAD1 | -134141 | -2.1213645 | 7.07E-06 |
| Peak87750 | chr7 | 91942455 | 91942797 | ANKIB1 | 67078 | -2.1213645 | 7.07E-06 |
| Peak12152 | chr10 | 74701556 | 74701800 | PLA2G12B | 12858 | -1.5151559 | 7.23E-06 |
| Peak12152 | chr10 | 74701556 | 74701800 | OIT3 | 48339 | -1.5151559 | 7.23E-06 |
| Peak29178 | chr14 | 71393252 | 71393514 | SIPA1L1 | -602659 | -1.4947841 | 7.38E-06 |
| Peak29178 | chr14 | 71393252 | 71393514 | PCNX | 19261 | -1.4947841 | 7.38E-06 |
| Peak42260 | chr18 | 43469721 | 43469891 | SIGLEC15 | 64329 | -2.0809285 | 7.70E-06 |
| Peak42260 | chr18 | 43469721 | 43469891 | EPG5 | 77434 | -2.0809285 | 7.70E-06 |
| Peak35170 | chr16 | 31200447 | 31200715 | FUS | 9150 | -2.3681878 | 7.71E-06 |

| | | | | | | | |
|---|---|---|---|---|---|---|---|
| Peak35170 | chr16 | 31200447 | 31200715 | PYCARD | 13732 | -2.3681878 | 7.71E-06 |
| Peak74526 | chr5 | 93330789 | 93331055 | POU5F2 | -253579 | -1.1672131 | 7.72E-06 |
| Peak74526 | chr5 | 93330789 | 93331055 | FAM172A | 116418 | -1.1672131 | 7.72E-06 |
| Peak80892 | chr6 | 97742472 | 97742593 | MMS22L | -11481 | -2.7019862 | 7.92E-06 |
| Peak68062 | chr4 | 26724846 | 26725212 | STIM2 | -137434 | -1.3290066 | 7.98E-06 |
| Peak68062 | chr4 | 26724846 | 26725212 | TBC1D19 | 139491 | -1.3290066 | 7.98E-06 |
| Peak80716 | chr6 | 90184479 | 90184614 | ANKRD6 | -87573 | -1.2694643 | 7.99E-06 |
| Peak80716 | chr6 | 90184479 | 90184614 | RRAGD | -62558 | -1.2694643 | 7.99E-06 |
| Peak82924 | chr6 | 145241858 | 145242020 | UTRN | 628974 | -2.5141193 | 8.02E-06 |
| Peak82924 | chr6 | 145241858 | 145242020 | EPM2A | 815221 | -2.5141193 | 8.02E-06 |
| Peak85618 | chr7 | 16385992 | 16386355 | MEOX2 | -659737 | -1.6197888 | 8.13E-06 |
| Peak85618 | chr7 | 16385992 | 16386355 | ISPD | 74773 | -1.6197888 | 8.13E-06 |
| Peak102058 | chrX | 138877438 | 138877636 | MCF2 | -87151 | -1.695071 | 8.14E-06 |
| Peak102058 | chrX | 138877438 | 138877636 | ATP11C | 36910 | -1.695071 | 8.14E-06 |
| Peak15670 | chr11 | 34729043 | 34729273 | EHF | 75147 | -1.8830011 | 8.15E-06 |
| Peak15670 | chr11 | 34729043 | 34729273 | APIP | 208888 | -1.8830011 | 8.15E-06 |
| Peak48532 | chr2 | 41683466 | 41683716 | SLC8A1 | -944090 | -1.9349886 | 8.22E-06 |
| Peak48532 | chr2 | 41683466 | 41683716 | PKDCC | -591569 | -1.9349886 | 8.22E-06 |
| Peak1539 | chr1 | 31255296 | 31255409 | LAPTM5 | -24686 | -1.7055958 | 8.35E-06 |
| Peak1539 | chr1 | 31255296 | 31255409 | SDC3 | 126255 | -1.7055958 | 8.35E-06 |
| Peak57536 | chr20 | 61436045 | 61436252 | OGFR | -38 | -1.0441522 | 8.38E-06 |
| Peak53901 | chr2 | 204042159 | 204042377 | CYP20A1 | -61395 | -1.4628329 | 8.67E-06 |
| Peak53901 | chr2 | 204042159 | 204042377 | CARF | 265151 | -1.4628329 | 8.67E-06 |
| Peak61549 | chr3 | 27882286 | 27882732 | CMC1 | -400577 | -1.9951526 | 8.93E-06 |
| Peak61549 | chr3 | 27882286 | 27882732 | EOMES | -118520 | -1.9951526 | 8.93E-06 |
| Peak91041 | chr8 | 24301883 | 24302087 | NEFM | -468540 | -3.5112326 | 9.02E-06 |
| Peak91041 | chr8 | 24301883 | 24302087 | ADAM7 | 3446 | -3.5112326 | 9.02E-06 |
| Peak6595 | chr1 | 172381245 | 172381355 | PIGC | 31926 | -2.4420573 | 9.29E-06 |
| Peak6595 | chr1 | 172381245 | 172381355 | DNM3 | 570679 | -2.4420573 | 9.29E-06 |
| Peak86902 | chr7 | 55604558 | 55604635 | VOPP1 | 35621 | -2.3588707 | 9.41E-06 |
| Peak86902 | chr7 | 55604558 | 55604635 | LANCL2 | 171456 | -2.3588707 | 9.41E-06 |
| Peak23744 | chr12 | 113189801 | 113189955 | RPH3A | -39582 | -2.1028278 | 9.47E-06 |
| Peak23744 | chr12 | 113189801 | 113189955 | PTPN11 | 333160 | -2.1028278 | 9.47E-06 |
| Peak49021 | chr2 | 58796946 | 58797217 | FANCL | -328597 | -1.8629091 | 9.62E-06 |
| Peak78613 | chr6 | 25010550 | 25010671 | LRRC16A | -269045 | -1.5111851 | 1.00E-05 |
| Peak78613 | chr6 | 25010550 | 25010671 | FAM65B | -99416 | -1.5111851 | 1.00E-05 |
| Peak81443 | chr6 | 112074834 | 112075015 | TRAF3IP2 | -147476 | -1.8270521 | 1.01E-05 |
| Peak81443 | chr6 | 112074834 | 112075015 | FYN | 119730 | -1.8270521 | 1.01E-05 |
| Peak69347 | chr4 | 82718084 | 82718241 | RASGEF1B | -325094 | -1.5679811 | 1.01E-05 |
| Peak69347 | chr4 | 82718084 | 82718241 | HNRNPD | 576946 | -1.5679811 | 1.01E-05 |
| Peak51722 | chr2 | 143623636 | 143623776 | LRP1B | -734436 | -2.0495934 | 1.03E-05 |
| Peak51722 | chr2 | 143623636 | 143623776 | KYNU | -11361 | -2.0495934 | 1.03E-05 |
| Peak83677 | chr6 | 160820093 | 160820188 | SLC22A3 | 50841 | -2.2174937 | 1.03E-05 |
| Peak83677 | chr6 | 160820093 | 160820188 | LPA | 267266 | -2.2174937 | 1.03E-05 |
| Peak58846 | chr21 | 48119777 | 48119890 | PRMT2 | 64272 | -1.0998023 | 1.05E-05 |
| Peak42572 | chr18 | 52496359 | 52496482 | RAB27B | 991 | -1.3403081 | 1.05E-05 |
| Peak86423 | chr7 | 37672702 | 37673019 | ELMO1 | -184009 | -2.5228466 | 1.05E-05 |
| Peak86423 | chr7 | 37672702 | 37673019 | GPR141 | -50614 | -2.5228466 | 1.05E-05 |

| | | | | | | | |
|---|---|---|---|---|---|---|---|
| Peak27532 | chr14 | 22976162 | 22976300 | DAD1 | 81944 | -1.5938038 | 1.06E-05 |
| Peak27532 | chr14 | 22976162 | 22976300 | OR4E2 | 842934 | -1.5938038 | 1.06E-05 |
| Peak2614 | chr1 | 55352793 | 55353118 | DHCR24 | -65 | -1.3572779 | 1.06E-05 |
| Peak3697 | chr1 | 93306256 | 93306521 | RPL5 | 8807 | -1.5763417 | 1.08E-05 |
| Peak3697 | chr1 | 93306256 | 93306521 | FAM69A | 120668 | -1.5763417 | 1.08E-05 |
| Peak92008 | chr8 | 60093282 | 60093372 | TOX | -61560 | -3.0117234 | 1.09E-05 |
| Peak94419 | chr8 | 142183098 | 142183159 | DENND3 | 44409 | -4.0280111 | 1.11E-05 |
| Peak94419 | chr8 | 142183098 | 142183159 | SLC45A4 | 55544 | -4.0280111 | 1.11E-05 |
| Peak50532 | chr2 | 100951548 | 100951734 | LONRF2 | -12446 | -3.2619846 | 1.16E-05 |
| Peak50532 | chr2 | 100951548 | 100951734 | ENSG00000 | 35366 | -3.2619846 | 1.16E-05 |
| Peak11246 | chr10 | 45463993 | 45464085 | RASSF4 | 8799 | -1.4948639 | 1.19E-05 |
| Peak11246 | chr10 | 45463993 | 45464085 | C10orf10 | 10219 | -1.4948639 | 1.19E-05 |
| Peak21616 | chr12 | 56061133 | 56061202 | METTL7B | -14162 | -3.1038085 | 1.21E-05 |
| Peak21616 | chr12 | 56061133 | 56061202 | OR10P1 | 30524 | -3.1038085 | 1.21E-05 |
| Peak75212 | chr5 | 114668994 | 114669157 | PGGT1B | -70507 | -2.5016733 | 1.22E-05 |
| Peak75212 | chr5 | 114668994 | 114669157 | FEM1C | 211515 | -2.5016733 | 1.22E-05 |
| Peak80001 | chr6 | 53177259 | 53177441 | GCM1 | -163723 | -1.2288627 | 1.26E-05 |
| Peak80001 | chr6 | 53177259 | 53177441 | ELOVL5 | 36398 | -1.2288627 | 1.26E-05 |
| Peak71561 | chr4 | 169828416 | 169828494 | CBR4 | 102954 | -2.8932682 | 1.28E-05 |
| Peak71561 | chr4 | 169828416 | 169828494 | PALLD | 410200 | -2.8932682 | 1.28E-05 |
| Peak43308 | chr18 | 77933090 | 77933275 | ADNP2 | 66268 | -1.5231928 | 1.30E-05 |
| Peak43308 | chr18 | 77933090 | 77933275 | PARD6G | 72246 | -1.5231928 | 1.30E-05 |
| Peak78117 | chr6 | 10752059 | 10752319 | ENSG00000 | 4162 | -1.1775321 | 1.32E-05 |
| Peak78117 | chr6 | 10752059 | 10752319 | MAK | 86539 | -1.1775321 | 1.32E-05 |
| Peak57118 | chr20 | 49002597 | 49003200 | PTPN1 | -123992 | -1.1846119 | 1.32E-05 |
| Peak57118 | chr20 | 49002597 | 49003200 | CEBPB | 195523 | -1.1846119 | 1.32E-05 |
| Peak73907 | chr5 | 72921824 | 72922098 | ARHGEF28 | -22 | -1.6211572 | 1.34E-05 |
| Peak34037 | chr16 | 3839748 | 3840019 | TRAP1 | -72286 | -1.1718716 | 1.36E-05 |
| Peak34037 | chr16 | 3839748 | 3840019 | CREBBP | 90843 | -1.1718716 | 1.36E-05 |
| Peak11339 | chr10 | 48133227 | 48133369 | AGAP9 | -82538 | -2.0277096 | 1.37E-05 |
| Peak11339 | chr10 | 48133227 | 48133369 | ASAH2C | -78280 | -2.0277096 | 1.37E-05 |
| Peak98009 | chr9 | 129544199 | 129544307 | ZBTB43 | -23041 | -1.936349 | 1.37E-05 |
| Peak98009 | chr9 | 129544199 | 129544307 | LMX1B | 167531 | -1.936349 | 1.37E-05 |
| Peak45317 | chr19 | 35224988 | 35225176 | ZNF181 | -772 | -1.0200569 | 1.39E-05 |
| Peak75096 | chr5 | 111093133 | 111093975 | STARD4 | -245346 | -1.2298727 | 1.39E-05 |
| Peak75096 | chr5 | 111093133 | 111093975 | NREP | 219074 | -1.2298727 | 1.39E-05 |
| Peak83748 | chr6 | 164088818 | 164089006 | QKI | 253237 | -1.0880487 | 1.40E-05 |
| Peak88959 | chr7 | 128099133 | 128099393 | METTL2B | -17520 | -1.5555693 | 1.40E-05 |
| Peak88959 | chr7 | 128099133 | 128099393 | HILPDA | 3318 | -1.5555693 | 1.40E-05 |
| Peak65379 | chr3 | 144014065 | 144014312 | C3orf58 | 323549 | -2.4764343 | 1.41E-05 |
| Peak92296 | chr8 | 68492396 | 68492645 | ARFGEF1 | -236609 | -1.0403105 | 1.42E-05 |
| Peak92296 | chr8 | 68492396 | 68492645 | CPA6 | 166059 | -1.0403105 | 1.42E-05 |
| Peak80379 | chr6 | 76533934 | 76534290 | MYO6 | 75063 | -3.0832536 | 1.44E-05 |
| Peak80379 | chr6 | 76533934 | 76534290 | IMPG1 | 248283 | -3.0832536 | 1.44E-05 |
| Peak74953 | chr5 | 106855743 | 106856041 | EFNA5 | 150704 | -1.7528211 | 1.50E-05 |
| Peak26155 | chr13 | 61038040 | 61038237 | TDRD3 | 66712 | -1.7432331 | 1.51E-05 |
| Peak26155 | chr13 | 61038040 | 61038237 | PCDH20 | 963843 | -1.7432331 | 1.51E-05 |
| Peak46122 | chr19 | 46389701 | 46389881 | IRF2BP1 | -415 | -1.3233513 | 1.54E-05 |

| | | | | | | | |
|---|---|---|---|---|---|---|---|
| Peak93699 | chr8 | 123501991 | 123502163 | HAS2 | -848447 | -1.5749479 | 1.57E-05 |
| Peak93699 | chr8 | 123501991 | 123502163 | ZHX2 | -291556 | -1.5749479 | 1.57E-05 |
| Peak31339 | chr15 | 39786778 | 39786956 | RASGRP1 | -929860 | -2.12992 | 1.61E-05 |
| Peak31339 | chr15 | 39786778 | 39786956 | THBS1 | -86427 | -2.12992 | 1.61E-05 |
| Peak96618 | chr9 | 80853092 | 80853296 | PSAT1 | -58865 | -1.7827546 | 1.63E-05 |
| Peak96618 | chr9 | 80853092 | 80853296 | CEP78 | 2071 | -1.7827546 | 1.63E-05 |
| Peak76379 | chr5 | 145316148 | 145316288 | SH3RF2 | -1222 | -2.4431855 | 1.68E-05 |
| Peak46512 | chr19 | 50432718 | 50432888 | NUP62 | -42 | -1.0041142 | 1.70E-05 |
| Peak46512 | chr19 | 50432718 | 50432888 | IL4I1 | -7 | -1.0041142 | 1.70E-05 |
| Peak46512 | chr19 | 50432718 | 50432888 | ATF5 | 403 | -1.0041142 | 1.70E-05 |
| Peak24964 | chr13 | 28133894 | 28134272 | MTIF3 | -109382 | -1.1228528 | 1.71E-05 |
| Peak24964 | chr13 | 28133894 | 28134272 | LNX2 | 60458 | -1.1228528 | 1.71E-05 |
| Peak47825 | chr2 | 24962303 | 24962539 | CENPO | -53593 | -1.3834435 | 1.75E-05 |
| Peak47825 | chr2 | 24962303 | 24962539 | NCOA1 | 247620 | -1.3834435 | 1.75E-05 |
| Peak96654 | chr9 | 82245983 | 82246209 | TLE4 | 59408 | -1.477995 | 1.75E-05 |
| Peak50887 | chr2 | 111604245 | 111604551 | BCL2L11 | -274108 | -1.1461208 | 1.79E-05 |
| Peak50887 | chr2 | 111604245 | 111604551 | ACOXL | 114248 | -1.1461208 | 1.79E-05 |
| Peak73912 | chr5 | 72989541 | 72989630 | ARHGEF28 | 67603 | -2.3202252 | 1.79E-05 |
| Peak73912 | chr5 | 72989541 | 72989630 | ENC1 | 947663 | -2.3202252 | 1.79E-05 |
| Peak18238 | chr11 | 110394996 | 110395356 | FDX1 | 94569 | -1.5054587 | 1.80E-05 |
| Peak18238 | chr11 | 110394996 | 110395356 | ARHGAP20 | 188275 | -1.5054587 | 1.80E-05 |
| Peak50038 | chr2 | 86969070 | 86969391 | CHMP3 | -20986 | -1.6446734 | 1.86E-05 |
| Peak50038 | chr2 | 86969070 | 86969391 | CD8A | 66288 | -1.6446734 | 1.86E-05 |
| Peak92035 | chr8 | 60885221 | 60885430 | TOX | -853559 | -1.4588999 | 1.89E-05 |
| Peak92035 | chr8 | 60885221 | 60885430 | CA8 | 308645 | -1.4588999 | 1.89E-05 |
| Peak93517 | chr8 | 117450021 | 117450286 | TRPS1 | -768926 | -1.0544662 | 1.90E-05 |
| Peak93517 | chr8 | 117450021 | 117450286 | EIF3H | 317906 | -1.0544662 | 1.90E-05 |
| Peak52878 | chr2 | 176440239 | 176440324 | ATP5G3 | -390947 | -2.6601779 | 1.94E-05 |
| Peak20833 | chr12 | 41916019 | 41916167 | PDZRN4 | 333843 | -1.8388589 | 1.95E-05 |
| Peak20833 | chr12 | 41916019 | 41916167 | GXYLT1 | 622588 | -1.8388589 | 1.95E-05 |
| Peak88844 | chr7 | 123364778 | 123365057 | WASL | 24203 | -1.965444 | 2.00E-05 |
| Peak88844 | chr7 | 123364778 | 123365057 | LMOD2 | 69057 | -1.965444 | 2.00E-05 |
| Peak72271 | chr5 | 5486319 | 5486596 | ADAMTS16 | 346015 | -1.7518644 | 2.03E-05 |
| Peak72271 | chr5 | 5486319 | 5486596 | MED10 | 892249 | -1.7518644 | 2.03E-05 |
| Peak38000 | chr17 | 20604043 | 20604166 | CDRT15L2 | 121068 | -2.4089664 | 2.03E-05 |
| Peak38000 | chr17 | 20604043 | 20604166 | USP22 | 342247 | -2.4089664 | 2.03E-05 |
| Peak22014 | chr12 | 66320471 | 66320622 | HMGA2 | 102636 | -1.9458315 | 2.08E-05 |
| Peak22014 | chr12 | 66320471 | 66320622 | ENSG00000 | 243218 | -1.9458315 | 2.08E-05 |
| Peak29305 | chr14 | 74107830 | 74107919 | ACOT6 | 24327 | -2.6088748 | 2.12E-05 |
| Peak29305 | chr14 | 74107830 | 74107919 | PNMA1 | 73253 | -2.6088748 | 2.12E-05 |
| Peak82177 | chr6 | 132811703 | 132811839 | MOXD1 | -89087 | -2.2041022 | 2.26E-05 |
| Peak82177 | chr6 | 132811703 | 132811839 | STX7 | 22430 | -2.2041022 | 2.26E-05 |
| Peak41526 | chr18 | 11760836 | 11761082 | CHMP1B | -90436 | -2.1513367 | 2.27E-05 |
| Peak41526 | chr18 | 11760836 | 11761082 | GNAL | 72004 | -2.1513367 | 2.27E-05 |
| Peak81349 | chr6 | 110027815 | 110028049 | GPR6 | -272366 | -2.1328665 | 2.36E-05 |
| Peak81349 | chr6 | 110027815 | 110028049 | FIG4 | 15417 | -2.1328665 | 2.36E-05 |
| Peak7005 | chr1 | 184358194 | 184358278 | TSEN15 | 337425 | -1.6385713 | 2.37E-05 |
| Peak7005 | chr1 | 184358194 | 184358278 | EDEM3 | 365811 | -1.6385713 | 2.37E-05 |

| | | | | | | | |
|---|---|---|---|---|---|---|---|
| Peak76682 | chr5 | 151849863 | 151850013 | NMUR2 | -65098 | -2.1640326 | 2.40E-05 |
| Peak92674 | chr8 | 82709320 | 82709478 | SNX16 | 45122 | -1.8693143 | 2.46E-05 |
| Peak92674 | chr8 | 82709320 | 82709478 | CHMP4C | 64730 | -1.8693143 | 2.46E-05 |
| Peak81541 | chr6 | 115323275 | 115323448 | HS3ST5 | -659153 | -1.2240227 | 2.47E-05 |
| Peak80883 | chr6 | 97394945 | 97395188 | NDUFAF4 | -49310 | -1.1300856 | 2.48E-05 |
| Peak80883 | chr6 | 97394945 | 97395188 | MMS22L | 335985 | -1.1300856 | 2.48E-05 |
| Peak82193 | chr6 | 133072650 | 133072809 | VNN3 | -16899 | -1.2858989 | 2.54E-05 |
| Peak82193 | chr6 | 133072650 | 133072809 | VNN2 | 6417 | -1.2858989 | 2.54E-05 |
| Peak42021 | chr18 | 32303195 | 32303362 | MAPRE2 | -318045 | -2.4245234 | 2.54E-05 |
| Peak42021 | chr18 | 32303195 | 32303362 | DTNA | 129976 | -2.4245234 | 2.54E-05 |
| Peak25060 | chr13 | 30731254 | 30731718 | UBL3 | -306665 | -1.4158518 | 2.55E-05 |
| Peak25060 | chr13 | 30731254 | 30731718 | KATNAL1 | 150135 | -1.4158518 | 2.55E-05 |
| Peak45499 | chr19 | 37862214 | 37862351 | ZNF527 | 220 | -1.2473426 | 2.55E-05 |
| Peak48803 | chr2 | 48455560 | 48455713 | FBXO11 | -322705 | -2.8139547 | 2.56E-05 |
| Peak48803 | chr2 | 48455560 | 48455713 | FOXN2 | -86212 | -2.8139547 | 2.56E-05 |
| Peak5896 | chr1 | 157670626 | 157670865 | FCRL3 | -99 | -1.1891996 | 2.59E-05 |
| Peak85753 | chr7 | 21422921 | 21423162 | SP8 | -596537 | -1.2496911 | 2.59E-05 |
| Peak85753 | chr7 | 21422921 | 21423162 | SP4 | -44610 | -1.2496911 | 2.59E-05 |
| Peak17805 | chr11 | 95808138 | 95808321 | MTMR2 | -150771 | -1.56311 | 2.62E-05 |
| Peak17805 | chr11 | 95808138 | 95808321 | MAML2 | 268114 | -1.56311 | 2.62E-05 |
| Peak22246 | chr12 | 70427161 | 70427394 | CNOT2 | -209499 | -2.5621472 | 2.67E-05 |
| Peak22246 | chr12 | 70427161 | 70427394 | MYRFL | 100962 | -2.5621472 | 2.67E-05 |
| Peak4086 | chr1 | 101797496 | 101797705 | S1PR1 | 95157 | -1.9831389 | 2.69E-05 |
| Peak4086 | chr1 | 101797496 | 101797705 | OLFM3 | 664985 | -1.9831389 | 2.69E-05 |
| Peak41672 | chr18 | 14435619 | 14435774 | ANKRD30B | -312542 | -1.615542 | 2.82E-05 |
| Peak41672 | chr18 | 14435619 | 14435774 | ZNF519 | -303268 | -1.615542 | 2.82E-05 |
| Peak47098 | chr2 | 166474 | 166742 | FAM110C | -120223 | -1.9337527 | 2.90E-05 |
| Peak47098 | chr2 | 166474 | 166742 | SH3YL1 | 97457 | -1.9337527 | 2.90E-05 |
| Peak83709 | chr6 | 161647391 | 161647736 | AGPAT4 | 47529 | -1.3042693 | 2.91E-05 |
| Peak83709 | chr6 | 161647391 | 161647736 | MAP3K4 | 234748 | -1.3042693 | 2.91E-05 |
| Peak92405 | chr8 | 73976090 | 73976261 | SBSPON | 29331 | -2.4556015 | 2.93E-05 |
| Peak92405 | chr8 | 73976090 | 73976261 | TERF1 | 55077 | -2.4556015 | 2.93E-05 |
| Peak89247 | chr7 | 134320879 | 134321061 | BPGM | -10613 | -1.8857937 | 3.05E-05 |
| Peak89247 | chr7 | 134320879 | 134321061 | AKR1B15 | 87082 | -1.8857937 | 3.05E-05 |
| Peak23745 | chr12 | 113190202 | 113190364 | RPH3A | -39177 | -1.5168173 | 3.07E-05 |
| Peak23745 | chr12 | 113190202 | 113190364 | PTPN11 | 333565 | -1.5168173 | 3.07E-05 |
| Peak66804 | chr3 | 188160335 | 188160590 | TPRG1 | -729300 | -1.1395995 | 3.10E-05 |
| Peak66804 | chr3 | 188160335 | 188160590 | LPP | 229742 | -1.1395995 | 3.10E-05 |
| Peak37676 | chr17 | 14233502 | 14233648 | HS3ST3B1 | 29175 | -2.121159 | 3.11E-05 |
| Peak37676 | chr17 | 14233502 | 14233648 | PMP22 | 932331 | -2.121159 | 3.11E-05 |
| Peak13058 | chr10 | 99116333 | 99116623 | FRAT2 | -22020 | -1.6877322 | 3.13E-05 |
| Peak13058 | chr10 | 99116333 | 99116623 | RRP12 | 44622 | -1.6877322 | 3.13E-05 |
| Peak24868 | chr13 | 26312079 | 26312352 | SHISA2 | 312953 | -1.4126925 | 3.14E-05 |
| Peak24868 | chr13 | 26312079 | 26312352 | ATP8A2 | 366007 | -1.4126925 | 3.14E-05 |
| Peak56667 | chr20 | 37405081 | 37405296 | PPP1R16B | -29159 | -1.7208514 | 3.19E-05 |
| Peak56667 | chr20 | 37405081 | 37405296 | ACTR5 | 28104 | -1.7208514 | 3.19E-05 |
| Peak66141 | chr3 | 171813074 | 171813313 | FNDC3B | 54850 | -1.627594 | 3.25E-05 |
| Peak66141 | chr3 | 171813074 | 171813313 | GHSR | 353052 | -1.627594 | 3.25E-05 |

| | | | | | | | |
|---|---|---|---|---|---|---|---|
| Peak61248 | chr3 | 16420460 | 16420752 | OXNAD1 | 113900 | -1.3224016 | 3.29E-05 |
| Peak61248 | chr3 | 16420460 | 16420752 | RFTN1 | 134607 | -1.3224016 | 3.29E-05 |
| Peak92017 | chr8 | 60374985 | 60375146 | TOX | -343299 | -1.4900148 | 3.34E-05 |
| Peak92017 | chr8 | 60374985 | 60375146 | CA8 | 818905 | -1.4900148 | 3.34E-05 |
| Peak27258 | chr13 | 113830664 | 113830868 | PROZ | 17798 | -1.0647586 | 3.35E-05 |
| Peak27258 | chr13 | 113830664 | 113830868 | PCID2 | 32217 | -1.0647586 | 3.35E-05 |
| Peak53483 | chr2 | 196514944 | 196515212 | SLC39A10 | -6393 | -1.0436363 | 3.43E-05 |
| Peak53029 | chr2 | 180167645 | 180167829 | SESTD1 | -38220 | -3.0125721 | 3.47E-05 |
| Peak53029 | chr2 | 180167645 | 180167829 | ZNF385B | 558495 | -3.0125721 | 3.47E-05 |
| Peak65360 | chr3 | 143675731 | 143675979 | SLC9A9 | -108482 | -1.5552693 | 3.48E-05 |
| Peak65360 | chr3 | 143675731 | 143675979 | C3orf58 | -14785 | -1.5552693 | 3.48E-05 |
| Peak42785 | chr18 | 60718310 | 60718491 | BCL2 | 268960 | -1.2027199 | 3.53E-05 |
| Peak42785 | chr18 | 60718310 | 60718491 | PHLPP1 | 335718 | -1.2027199 | 3.53E-05 |
| Peak47826 | chr2 | 24962590 | 24962798 | CENPO | -53320 | -1.3731254 | 3.53E-05 |
| Peak47826 | chr2 | 24962590 | 24962798 | NCOA1 | 247893 | -1.3731254 | 3.53E-05 |
| Peak96261 | chr9 | 71161527 | 71161678 | PIP5K1B | -159013 | -1.8766864 | 3.56E-05 |
| Peak96261 | chr9 | 71161527 | 71161678 | TMEM252 | -5820 | -1.8766864 | 3.56E-05 |
| Peak82187 | chr6 | 132969527 | 132969768 | TAAR1 | -2506 | -1.429458 | 3.60E-05 |
| Peak47570 | chr2 | 12204388 | 12204706 | TRIB2 | -652468 | -1.0742691 | 3.68E-05 |
| Peak47570 | chr2 | 12204388 | 12204706 | LPIN1 | 386826 | -1.0742691 | 3.68E-05 |
| Peak80023 | chr6 | 53499218 | 53499449 | GCLC | -89407 | -1.5153974 | 3.68E-05 |
| Peak80023 | chr6 | 53499218 | 53499449 | KLHL31 | 31172 | -1.5153974 | 3.68E-05 |
| Peak62349 | chr3 | 46997307 | 46997732 | NBEAL2 | -23653 | -1.1914895 | 3.69E-05 |
| Peak62349 | chr3 | 46997307 | 46997732 | PTH1R | 72673 | -1.1914895 | 3.69E-05 |
| Peak19257 | chr11 | 134526402 | 134526972 | B3GAT1 | -264440 | -2.1707864 | 3.74E-05 |
| Peak47887 | chr2 | 25811535 | 25811719 | DNMT3A | -246168 | -2.4705779 | 3.80E-05 |
| Peak47887 | chr2 | 25811535 | 25811719 | DTNB | 84876 | -2.4705779 | 3.80E-05 |
| Peak17572 | chr11 | 86383504 | 86383610 | ME3 | 121 | -1.4741449 | 3.90E-05 |
| Peak96501 | chr9 | 77736936 | 77737034 | PCSK5 | -768575 | -1.7886599 | 3.93E-05 |
| Peak96501 | chr9 | 77736936 | 77737034 | OSTF1 | 33526 | -1.7886599 | 3.93E-05 |
| Peak45614 | chr19 | 39616207 | 39616520 | PAK4 | -90 | -1.0564767 | 3.95E-05 |
| Peak52112 | chr2 | 157520411 | 157520565 | GALNT5 | -593622 | -1.2946337 | 4.04E-05 |
| Peak52112 | chr2 | 157520411 | 157520565 | GPD2 | 228535 | -1.2946337 | 4.04E-05 |
| Peak28033 | chr14 | 39812129 | 39812264 | CTAGE5 | 75869 | -1.7055496 | 4.06E-05 |
| Peak28033 | chr14 | 39812129 | 39812264 | FBXO33 | 89507 | -1.7055496 | 4.06E-05 |
| Peak20202 | chr12 | 18318564 | 18318832 | PIK3C2G | -95776 | -1.3152355 | 4.07E-05 |
| Peak20202 | chr12 | 18318564 | 18318832 | RERGL | -75571 | -1.3152355 | 4.07E-05 |
| Peak64719 | chr3 | 126331448 | 126331611 | TXNRD3 | 42468 | -1.7916824 | 4.12E-05 |
| Peak64719 | chr3 | 126331448 | 126331611 | CHST13 | 88404 | -1.7916824 | 4.12E-05 |
| Peak47650 | chr2 | 15451719 | 15451887 | NBAS | 249651 | -1.3580526 | 4.14E-05 |
| Peak47103 | chr2 | 223566 | 223763 | FAM110C | -177280 | -1.8078253 | 4.15E-05 |
| Peak47103 | chr2 | 223566 | 223763 | SH3YL1 | 40400 | -1.8078253 | 4.15E-05 |
| Peak13820 | chr10 | 119412748 | 119412954 | EMX2 | 110896 | -1.9734381 | 4.16E-05 |
| Peak13820 | chr10 | 119412748 | 119412954 | RAB11FIP2 | 393263 | -1.9734381 | 4.16E-05 |
| Peak67759 | chr4 | 10589371 | 10589579 | ZNF518B | -130491 | -1.5086636 | 4.21E-05 |
| Peak67759 | chr4 | 10589371 | 10589579 | CLNK | 97014 | -1.5086636 | 4.21E-05 |
| Peak92410 | chr8 | 74069139 | 74069271 | SBSPON | -63698 | -2.1703072 | 4.22E-05 |
| Peak92410 | chr8 | 74069139 | 74069271 | RPL7 | 136928 | -2.1703072 | 4.22E-05 |

| | | | | | | | |
|---|---|---|---|---|---|---|---|
| Peak11425 | chr10 | 51026109 | 51026233 | OGDHL | -55802 | -3.3962299 | 4.29E-05 |
| Peak11425 | chr10 | 51026109 | 51026233 | PARG | 104544 | -3.3962299 | 4.29E-05 |
| Peak17720 | chr11 | 94058985 | 94059084 | FOLR4 | 20276 | -2.2002677 | 4.31E-05 |
| Peak17720 | chr11 | 94058985 | 94059084 | GPR83 | 75550 | -2.2002677 | 4.31E-05 |
| Peak44255 | chr19 | 11354105 | 11354267 | C19orf80 | 3891 | -1.3618322 | 4.41E-05 |
| Peak44255 | chr19 | 11354105 | 11354267 | DOCK6 | 18942 | -1.3618322 | 4.41E-05 |
| Peak75483 | chr5 | 126926204 | 126926329 | CTXN3 | -58469 | -1.4028525 | 4.41E-05 |
| Peak75483 | chr5 | 126926204 | 126926329 | PRRC1 | 72966 | -1.4028525 | 4.41E-05 |
| Peak80383 | chr6 | 76545067 | 76545121 | MYO6 | 86045 | -3.0087715 | 4.42E-05 |
| Peak80383 | chr6 | 76545067 | 76545121 | IMPG1 | 237301 | -3.0087715 | 4.42E-05 |
| Peak35310 | chr16 | 47578203 | 47578533 | PHKB | 83130 | -1.3830401 | 4.45E-05 |
| Peak35310 | chr16 | 47578203 | 47578533 | ABCC12 | 602313 | -1.3830401 | 4.45E-05 |
| Peak83432 | chr6 | 157468140 | 157468341 | TMEM242 | 276392 | -1.2525942 | 4.53E-05 |
| Peak83432 | chr6 | 157468140 | 157468341 | ARID1B | 369178 | -1.2525942 | 4.53E-05 |
| Peak661 | chr1 | 15623612 | 15623809 | EFHD2 | -112680 | -1.3192128 | 4.61E-05 |
| Peak661 | chr1 | 15623612 | 15623809 | TMEM51 | 144683 | -1.3192128 | 4.61E-05 |
| Peak71270 | chr4 | 153878658 | 153878920 | TRIM2 | -246829 | -2.0856552 | 4.66E-05 |
| Peak71270 | chr4 | 153878658 | 153878920 | ARFIP1 | 177700 | -2.0856552 | 4.66E-05 |
| Peak46108 | chr19 | 46273843 | 46274160 | SIX5 | -1518 | -1.0539258 | 4.68E-05 |
| Peak55568 | chr20 | 2846890 | 2847118 | GNRH2 | -177264 | -2.2567295 | 4.74E-05 |
| Peak55568 | chr20 | 2846890 | 2847118 | PTPRA | 2174 | -2.2567295 | 4.74E-05 |
| Peak67850 | chr4 | 15773074 | 15773329 | CD38 | -6699 | -1.8025015 | 4.80E-05 |
| Peak67850 | chr4 | 15773074 | 15773329 | BST1 | 68629 | -1.8025015 | 4.80E-05 |
| Peak49626 | chr2 | 73220340 | 73220426 | EMX1 | 75779 | -1.5761003 | 4.82E-05 |
| Peak49626 | chr2 | 73220340 | 73220426 | SFXN5 | 78582 | -1.5761003 | 4.82E-05 |
| Peak13960 | chr10 | 122473224 | 122473402 | WDR11 | -137374 | -1.9484329 | 4.83E-05 |
| Peak13960 | chr10 | 122473224 | 122473402 | PPAPDC1A | 256847 | -1.9484329 | 4.83E-05 |
| Peak37346 | chr17 | 6926991 | 6927295 | BCL6B | 804 | -1.0337122 | 4.88E-05 |
| Peak86122 | chr7 | 29302234 | 29302362 | PRR15 | -301129 | -2.2871704 | 4.93E-05 |
| Peak86122 | chr7 | 29302234 | 29302362 | CPVL | -67406 | -2.2871704 | 4.93E-05 |
| Peak34662 | chr16 | 22584582 | 22584744 | HS3ST2 | -240835 | -1.0595781 | 5.01E-05 |
| Peak34662 | chr16 | 22584582 | 22584744 | NPIPB5 | 59819 | -1.0595781 | 5.01E-05 |
| Peak20566 | chr12 | 28931011 | 28931171 | FAR2 | -445507 | -1.8265602 | 5.01E-05 |
| Peak20566 | chr12 | 28931011 | 28931171 | CCDC91 | 587712 | -1.8265602 | 5.01E-05 |
| Peak58975 | chr22 | 19166285 | 19166554 | SLC25A1 | -77 | -1.1148226 | 5.04E-05 |
| Peak3790 | chr1 | 94486633 | 94486859 | GCLM | -111780 | -1.185364 | 5.07E-05 |
| Peak3790 | chr1 | 94486633 | 94486859 | ABCA4 | 99942 | -1.185364 | 5.07E-05 |
| Peak42979 | chr18 | 67834045 | 67834145 | CD226 | -209863 | -2.3060611 | 5.19E-05 |
| Peak42979 | chr18 | 67834045 | 67834145 | RTTN | 39086 | -2.3060611 | 5.19E-05 |
| Peak52870 | chr2 | 176072853 | 176073118 | ATP5G3 | -23651 | -2.7422857 | 5.21E-05 |
| Peak37288 | chr17 | 5087649 | 5087941 | ZNF594 | -103 | -1.3669706 | 5.22E-05 |
| Peak68491 | chr4 | 46841221 | 46841449 | GABRA2 | -364088 | -1.3938178 | 5.33E-05 |
| Peak68491 | chr4 | 46841221 | 46841449 | COX7B2 | 69889 | -1.3938178 | 5.33E-05 |
| Peak4180 | chr1 | 108486275 | 108486494 | VAV3 | 21381 | -1.3324147 | 5.50E-05 |
| Peak4180 | chr1 | 108486275 | 108486494 | NTNG1 | 802943 | -1.3324147 | 5.50E-05 |
| Peak95190 | chr9 | 15412462 | 15412665 | FREM1 | -501571 | -1.7350001 | 5.53E-05 |
| Peak95190 | chr9 | 15412462 | 15412665 | SNAPC3 | -10138 | -1.7350001 | 5.53E-05 |
| Peak3303 | chr1 | 85488628 | 85488674 | MCOLN2 | -26028 | -2.6261244 | 5.61E-05 |

| | | | | | | | |
|---|---|---|---|---|---|---|---|
| Peak3303 | chr1 | 85488628 | 85488674 | MCOLN3 | 25478 | -2.6261244 | 5.61E-05 |
| Peak83464 | chr6 | 157948510 | 157948651 | SNX9 | -295715 | -1.4352804 | 5.61E-05 |
| Peak83464 | chr6 | 157948510 | 157948651 | ZDHHC14 | 146416 | -1.4352804 | 5.61E-05 |
| Peak23534 | chr12 | 108972382 | 108972441 | ISCU | 16035 | -2.3825476 | 5.62E-05 |
| Peak23534 | chr12 | 108972382 | 108972441 | TMEM119 | 19488 | -2.3825476 | 5.62E-05 |
| Peak60663 | chr3 | 3615163 | 3615269 | CRBN | -393822 | -3.7928399 | 5.62E-05 |
| Peak60663 | chr3 | 3615163 | 3615269 | LRRN1 | -225905 | -3.7928399 | 5.62E-05 |
| Peak82192 | chr6 | 133071871 | 133072016 | VNN3 | -16113 | -1.4050837 | 5.70E-05 |
| Peak82192 | chr6 | 133071871 | 133072016 | VNN2 | 7203 | -1.4050837 | 5.70E-05 |
| Peak82007 | chr6 | 130182063 | 130182579 | TMEM244 | 371 | -1.3140309 | 5.73E-05 |
| Peak61245 | chr3 | 16405957 | 16406104 | OXNAD1 | 99325 | -2.4169525 | 5.74E-05 |
| Peak61245 | chr3 | 16405957 | 16406104 | RFTN1 | 149182 | -2.4169525 | 5.74E-05 |
| Peak47904 | chr2 | 26039250 | 26039385 | DTNB | -142815 | -1.8786067 | 5.80E-05 |
| Peak47904 | chr2 | 26039250 | 26039385 | ASXL2 | 62067 | -1.8786067 | 5.80E-05 |
| Peak5539 | chr1 | 152452648 | 152452781 | CRNN | -65976 | -2.2213942 | 5.80E-05 |
| Peak5539 | chr1 | 152452648 | 152452781 | LCE5A | -30605 | -2.2213942 | 5.80E-05 |
| Peak37620 | chr17 | 10060333 | 10060506 | RCVRN | -251482 | -2.0309523 | 5.80E-05 |
| Peak37620 | chr17 | 10060333 | 10060506 | GAS7 | 41448 | -2.0309523 | 5.80E-05 |
| Peak72603 | chr5 | 32159915 | 32160001 | GOLPH3 | 14498 | -2.7258133 | 5.92E-05 |
| Peak72603 | chr5 | 32159915 | 32160001 | PDZD2 | 520441 | -2.7258133 | 5.92E-05 |
| Peak64179 | chr3 | 112016574 | 112016828 | SLC9C1 | -3596 | -1.6008515 | 5.95E-05 |
| Peak48259 | chr2 | 33722278 | 33722578 | RASGRP3 | 21133 | -1.360768 | 5.98E-05 |

**Table S6. List of top 500 ATAC peaks higher in R1-27+**

| peak_id | chr | start | end | gene | distTSS | log2FoldChar | pvalue |
|---|---|---|---|---|---|---|---|
| Peak66120 | chr3 | 171531802 | 171532204 | PLD1 | -3719 | 1.7045985 | 6.68E-14 |
| Peak100785 | chrX | 69209019 | 69209253 | AWAT2 | 60652 | 4.47779952 | 4.28E-12 |
| Peak100785 | chrX | 69209019 | 69209253 | EDA | 373225 | 4.47779952 | 4.28E-12 |
| Peak93788 | chr8 | 124560582 | 124560866 | FBXO32 | -7278 | 1.46209015 | 7.87E-12 |
| Peak93788 | chr8 | 124560582 | 124560866 | KLHL38 | 104466 | 1.46209015 | 7.87E-12 |
| Peak4336 | chr1 | 110546534 | 110546752 | STRIP1 | -30595 | 2.99514778 | 1.38E-11 |
| Peak4336 | chr1 | 110546534 | 110546752 | AHCYL1 | 19335 | 2.99514778 | 1.38E-11 |
| Peak19989 | chr12 | 12151552 | 12151869 | BCL2L14 | -72162 | 2.89954466 | 1.64E-11 |
| Peak19989 | chr12 | 12151552 | 12151869 | ETV6 | 348923 | 2.89954466 | 1.64E-11 |
| Peak92725 | chr8 | 86413427 | 86413638 | CA2 | 37452 | 3.48748731 | 2.16E-11 |
| Peak92725 | chr8 | 86413427 | 86413638 | REXO1L1 | 162193 | 3.48748731 | 2.16E-11 |
| Peak42815 | chr18 | 60891872 | 60892109 | BCL2 | 95370 | 1.91690813 | 2.39E-11 |
| Peak42815 | chr18 | 60891872 | 60892109 | PHLPP1 | 509308 | 1.91690813 | 2.39E-11 |
| Peak56190 | chr20 | 25218475 | 25218738 | PYGB | -10098 | 1.68447195 | 2.52E-11 |
| Peak56190 | chr20 | 25218475 | 25218738 | ENTPD6 | 42251 | 1.68447195 | 2.52E-11 |
| Peak14890 | chr11 | 9023305 | 9023619 | TMEM9B | -37142 | 1.7424953 | 4.83E-11 |
| Peak14890 | chr11 | 9023305 | 9023619 | NRIP3 | 2134 | 1.7424953 | 4.83E-11 |
| Peak2947 | chr1 | 67637025 | 67637131 | IL12RB2 | -135969 | 1.93032603 | 5.65E-11 |
| Peak2947 | chr1 | 67637025 | 67637131 | IL23R | 4995 | 1.93032603 | 5.65E-11 |
| Peak69480 | chr4 | 87515305 | 87515561 | PTPN13 | -35 | 4.28955967 | 8.05E-11 |
| Peak48454 | chr2 | 38880762 | 38881475 | HNRNPLL | -50948 | 1.07444506 | 1.21E-10 |
| Peak48454 | chr2 | 38880762 | 38881475 | GALM | -11933 | 1.07444506 | 1.21E-10 |
| Peak67768 | chr4 | 11074680 | 11074945 | CLNK | -388324 | 4.06805451 | 1.47E-10 |
| Peak67768 | chr4 | 11074680 | 11074945 | HS3ST1 | 356576 | 4.06805451 | 1.47E-10 |
| Peak75519 | chr5 | 128450614 | 128450767 | ADAMTS19 | -345267 | 4.56295789 | 1.65E-10 |
| Peak75519 | chr5 | 128450614 | 128450767 | ISOC1 | 20247 | 4.56295789 | 1.65E-10 |
| Peak61503 | chr3 | 27387062 | 27387361 | NEK10 | 23700 | 3.93886064 | 1.87E-10 |
| Peak61503 | chr3 | 27387062 | 27387361 | LRRC3B | 722915 | 3.93886064 | 1.87E-10 |
| Peak10570 | chr10 | 24995389 | 24995674 | ARHGAP21 | 17065 | 2.32894327 | 2.05E-10 |
| Peak10570 | chr10 | 24995389 | 24995674 | KIAA1217 | 497439 | 2.32894327 | 2.05E-10 |
| Peak20035 | chr12 | 12642154 | 12642445 | MANSC1 | -138825 | 3.62319872 | 2.18E-10 |
| Peak20035 | chr12 | 12642154 | 12642445 | DUSP16 | 73017 | 3.62319872 | 2.18E-10 |
| Peak70307 | chr4 | 114925733 | 114925982 | UGT8 | -593753 | 2.15160227 | 3.95E-10 |
| Peak70307 | chr4 | 114925733 | 114925982 | ARSJ | -25001 | 2.15160227 | 3.95E-10 |
| Peak51558 | chr2 | 134938855 | 134939114 | MGAT5 | 61232 | 2.26665666 | 5.42E-10 |
| Peak51558 | chr2 | 134938855 | 134939114 | TMEM163 | 537585 | 2.26665666 | 5.42E-10 |
| Peak97149 | chr9 | 99637268 | 99637707 | ZNF782 | -20718 | 1.9857191 | 6.18E-10 |
| Peak97149 | chr9 | 99637268 | 99637707 | HIATL2 | 138374 | 1.9857191 | 6.18E-10 |
| Peak87782 | chr7 | 92300919 | 92301110 | RBM48 | 142928 | 2.68674228 | 6.96E-10 |
| Peak87782 | chr7 | 92300919 | 92301110 | CDK6 | 162216 | 2.68674228 | 6.96E-10 |
| Peak88645 | chr7 | 114871072 | 114871216 | MDFIC | 308935 | 1.91347429 | 8.42E-10 |
| Peak88645 | chr7 | 114871072 | 114871216 | TFEC | 799651 | 1.91347429 | 8.42E-10 |
| Peak7396 | chr1 | 198157252 | 198157736 | NEK7 | 31401 | 1.56906732 | 8.62E-10 |
| Peak7396 | chr1 | 198157252 | 198157736 | ATP6V1G3 | 352581 | 1.56906732 | 8.62E-10 |
| Peak52298 | chr2 | 161339275 | 161339546 | ITGB6 | -282599 | 1.88063231 | 8.67E-10 |
| Peak52298 | chr2 | 161339275 | 161339546 | RBMS1 | 10894 | 1.88063231 | 8.67E-10 |

| | | | | | | | |
|---|---|---|---|---|---|---|---|
| Peak13201 | chr10 | 102266385 | 102266551 | SEC31B | 13123 | 3.43915614 | 9.01E-10 |
| Peak13201 | chr10 | 102266385 | 102266551 | WNT8B | 43670 | 3.43915614 | 9.01E-10 |
| Peak70640 | chr4 | 124521150 | 124521442 | SPRY1 | 200631 | 1.38937141 | 1.05E-09 |
| Peak62113 | chr3 | 43222675 | 43222905 | SNRK | -105242 | 2.06006441 | 1.07E-09 |
| Peak62113 | chr3 | 43222675 | 43222905 | GTDC2 | -75222 | 2.06006441 | 1.07E-09 |
| Peak19867 | chr12 | 10350569 | 10350782 | GABARAPL1 | -14728 | 1.89772894 | 1.15E-09 |
| Peak19867 | chr12 | 10350569 | 10350782 | TMEM52B | 19064 | 1.89772894 | 1.15E-09 |
| Peak49203 | chr2 | 64439533 | 64439754 | LGALSL | -241582 | 1.61810775 | 1.17E-09 |
| Peak49203 | chr2 | 64439533 | 64439754 | PELI1 | -68056 | 1.61810775 | 1.17E-09 |
| Peak10693 | chr10 | 27736581 | 27736709 | RAB18 | -56552 | 3.13114799 | 1.24E-09 |
| Peak10693 | chr10 | 27736581 | 27736709 | PTCHD3 | -33348 | 3.13114799 | 1.24E-09 |
| Peak73304 | chr5 | 54430028 | 54430231 | GPX8 | -25816 | 2.75714434 | 1.28E-09 |
| Peak73304 | chr5 | 54430028 | 54430231 | GZMA | 31654 | 2.75714434 | 1.28E-09 |
| Peak10067 | chr10 | 11917849 | 11918032 | ECHDC3 | 133576 | 4.39011949 | 1.41E-09 |
| Peak10067 | chr10 | 11917849 | 11918032 | UPF2 | 159955 | 4.39011949 | 1.41E-09 |
| Peak28372 | chr14 | 52924673 | 52924974 | TXNDC16 | 94400 | 2.5395602 | 1.41E-09 |
| Peak28372 | chr14 | 52924673 | 52924974 | PTGER2 | 143711 | 2.5395602 | 1.41E-09 |
| Peak72356 | chr5 | 11222324 | 11222600 | DAP | -461078 | 4.63731839 | 1.67E-09 |
| Peak72356 | chr5 | 11222324 | 11222600 | CTNND2 | 681693 | 4.63731839 | 1.67E-09 |
| Peak89963 | chr7 | 151105870 | 151106385 | NUB1 | 67331 | 1.84456534 | 1.67E-09 |
| Peak89963 | chr7 | 151105870 | 151106385 | RHEB | 110882 | 1.84456534 | 1.67E-09 |
| Peak65573 | chr3 | 150994510 | 150994695 | GPR171 | -73624 | 3.11892861 | 1.68E-09 |
| Peak65573 | chr3 | 150994510 | 150994695 | P2RY14 | 1652 | 3.11892861 | 1.68E-09 |
| Peak15227 | chr11 | 16830182 | 16830386 | C11orf58 | 70336 | 5.04607794 | 1.71E-09 |
| Peak15227 | chr11 | 16830182 | 16830386 | PLEKHA7 | 205675 | 5.04607794 | 1.71E-09 |
| Peak66375 | chr3 | 179457698 | 179457897 | USP13 | 87255 | 3.88039471 | 1.80E-09 |
| Peak66375 | chr3 | 179457698 | 179457897 | PEX5L | 296920 | 3.88039471 | 1.80E-09 |
| Peak51951 | chr2 | 149638200 | 149638496 | LYPD6B | -256633 | 4.12083624 | 1.93E-09 |
| Peak51951 | chr2 | 149638200 | 149638496 | KIF5C | 5529 | 4.12083624 | 1.93E-09 |
| Peak77125 | chr5 | 169681051 | 169681425 | LCP2 | 43993 | 1.23425091 | 2.53E-09 |
| Peak77125 | chr5 | 169681051 | 169681425 | FOXI1 | 148337 | 1.23425091 | 2.53E-09 |
| Peak80771 | chr6 | 90966798 | 90966990 | BACH2 | 39567 | 2.5052281 | 2.85E-09 |
| Peak80771 | chr6 | 90966798 | 90966990 | GJA10 | 362706 | 2.5052281 | 2.85E-09 |
| Peak72555 | chr5 | 24272906 | 24273100 | CDH10 | 372084 | 3.0121854 | 3.56E-09 |
| Peak72555 | chr5 | 24272906 | 24273100 | PRDM9 | 765279 | 3.0121854 | 3.56E-09 |
| Peak70631 | chr4 | 124399075 | 124399333 | SPRY1 | 78539 | 2.14915568 | 3.82E-09 |
| Peak22397 | chr12 | 76311593 | 76311796 | KRR1 | -406294 | 2.05274092 | 4.11E-09 |
| Peak22397 | chr12 | 76311593 | 76311796 | PHLDA1 | 116017 | 2.05274092 | 4.11E-09 |
| Peak91797 | chr8 | 53676918 | 53677179 | NPBWR1 | -173942 | 1.3787661 | 4.15E-09 |
| Peak91797 | chr8 | 53676918 | 53677179 | RB1CC1 | -50057 | 1.3787661 | 4.15E-09 |
| Peak87170 | chr7 | 69772775 | 69773023 | WBSCR17 | -824256 | 2.93853719 | 4.24E-09 |
| Peak87170 | chr7 | 69772775 | 69773023 | AUTS2 | 708580 | 2.93853719 | 4.24E-09 |
| Peak14155 | chr10 | 127731635 | 127731872 | FANK1 | 146646 | 3.89586929 | 4.62E-09 |
| Peak14155 | chr10 | 127731635 | 127731872 | ADAM12 | 345270 | 3.89586929 | 4.62E-09 |
| Peak83394 | chr6 | 156717433 | 156717885 | NOX3 | -940622 | 2.40352069 | 5.29E-09 |
| Peak83394 | chr6 | 156717433 | 156717885 | ARID1B | -381404 | 2.40352069 | 5.29E-09 |
| Peak88646 | chr7 | 114871280 | 114871506 | MDFIC | 309184 | 1.50243459 | 5.45E-09 |
| Peak88646 | chr7 | 114871280 | 114871506 | TFEC | 799402 | 1.50243459 | 5.45E-09 |

| | | | | | | | |
|---|---|---|---|---|---|---|---|
| Peak94804 | chr9 | 2570687 | 2571062 | VLDLR | -50959 | 1.25056322 | 5.63E-09 |
| Peak94804 | chr9 | 2570687 | 2571062 | SMARCA2 | 548930 | 1.25056322 | 5.63E-09 |
| Peak83569 | chr6 | 159229922 | 159230315 | EZR | 10325 | 2.1350971 | 5.84E-09 |
| Peak83569 | chr6 | 159229922 | 159230315 | SYTL3 | 147768 | 2.1350971 | 5.84E-09 |
| Peak34466 | chr16 | 17684967 | 17685342 | XYLT1 | -120417 | 1.22379903 | 5.92E-09 |
| Peak34466 | chr16 | 17684967 | 17685342 | NOMO2 | 888273 | 1.22379903 | 5.92E-09 |
| Peak63209 | chr3 | 71116088 | 71116231 | FOXP1 | 63828 | 1.81282923 | 5.99E-09 |
| Peak78521 | chr6 | 20837520 | 20837739 | SOX4 | -756342 | 2.06585363 | 6.45E-09 |
| Peak78521 | chr6 | 20837520 | 20837739 | CDKAL1 | 302942 | 2.06585363 | 6.45E-09 |
| Peak93743 | chr8 | 124096322 | 124096788 | TBC1D31 | 11635 | 1.88242052 | 6.47E-09 |
| Peak93743 | chr8 | 124096322 | 124096788 | ZHX1 | 190180 | 1.88242052 | 6.47E-09 |
| Peak95108 | chr9 | 7967516 | 7967719 | C9orf123 | -167551 | 2.49547324 | 6.61E-09 |
| Peak24952 | chr13 | 28006787 | 28006998 | GTF3A | 8212 | 3.4241505 | 6.72E-09 |
| Peak24952 | chr13 | 28006787 | 28006998 | MTIF3 | 17808 | 3.4241505 | 6.72E-09 |
| Peak55669 | chr20 | 4678534 | 4678801 | PRND | -23888 | 2.97692363 | 7.47E-09 |
| Peak55669 | chr20 | 4678534 | 4678801 | PRNP | 11786 | 2.97692363 | 7.47E-09 |
| Peak18134 | chr11 | 107454613 | 107454796 | ALKBH8 | -18233 | 2.60647646 | 7.61E-09 |
| Peak18134 | chr11 | 107454613 | 107454796 | ELMOD1 | -7251 | 2.60647646 | 7.61E-09 |
| Peak28412 | chr14 | 53427415 | 53427647 | FERMT2 | -9716 | 2.45823764 | 8.15E-09 |
| Peak28412 | chr14 | 53427415 | 53427647 | DDHD1 | 192285 | 2.45823764 | 8.15E-09 |
| Peak75453 | chr5 | 126147597 | 126147920 | LMNB1 | 34919 | 1.15055438 | 8.21E-09 |
| Peak75453 | chr5 | 126147597 | 126147920 | 3-Mar | 218741 | 1.15055438 | 8.21E-09 |
| Peak44859 | chr19 | 18625724 | 18626009 | ISYNA1 | -76756 | 2.59146802 | 8.22E-09 |
| Peak44859 | chr19 | 18625724 | 18626009 | ELL | 7070 | 2.59146802 | 8.22E-09 |
| Peak9550 | chr10 | 310317 | 311260 | ZMYND11 | 130365 | 1.27411249 | 8.42E-09 |
| Peak9550 | chr10 | 310317 | 311260 | DIP2C | 424817 | 1.27411249 | 8.42E-09 |
| Peak28389 | chr14 | 53173125 | 53173269 | PSMC6 | -693 | 1.97587263 | 8.47E-09 |
| Peak86338 | chr7 | 36248696 | 36249124 | ANLN | -180505 | 4.18921056 | 8.47E-09 |
| Peak86338 | chr7 | 36248696 | 36249124 | EEPD1 | 56152 | 4.18921056 | 8.47E-09 |
| Peak12344 | chr10 | 79602668 | 79602915 | KCNMA1 | -205392 | 3.12185909 | 8.61E-09 |
| Peak12344 | chr10 | 79602668 | 79602915 | DLG5 | 83492 | 3.12185909 | 8.61E-09 |
| Peak15441 | chr11 | 23710473 | 23710794 | SVIP | -859223 | 2.7427324 | 8.73E-09 |
| Peak15441 | chr11 | 23710473 | 23710794 | LUZP2 | -808090 | 2.7427324 | 8.73E-09 |
| Peak16370 | chr11 | 61792917 | 61793542 | INCENP | -98215 | 1.15589726 | 9.07E-09 |
| Peak16370 | chr11 | 61792917 | 61793542 | FTH1 | -58098 | 1.15589726 | 9.07E-09 |
| Peak99826 | chrX | 19645775 | 19645975 | MAP3K15 | -112496 | 2.29656977 | 9.42E-09 |
| Peak99826 | chrX | 19645775 | 19645975 | SH3KBP1 | 259844 | 2.29656977 | 9.42E-09 |
| Peak25325 | chr13 | 40670542 | 40670771 | COG6 | 440843 | 1.41396085 | 9.43E-09 |
| Peak25325 | chr13 | 40670542 | 40670771 | FOXO1 | 570077 | 1.41396085 | 9.43E-09 |
| Peak80697 | chr6 | 90007757 | 90007961 | GABRR1 | -80707 | 1.80031068 | 9.86E-09 |
| Peak80697 | chr6 | 90007757 | 90007961 | GABRR2 | 17159 | 1.80031068 | 9.86E-09 |
| Peak21025 | chr12 | 47605917 | 47606059 | AMIGO2 | -132340 | 1.26407326 | 9.97E-09 |
| Peak21025 | chr12 | 47605917 | 47606059 | RPAP3 | 493856 | 1.26407326 | 9.97E-09 |
| Peak87346 | chr7 | 74561066 | 74561302 | TRIM73 | -463719 | 1.90346116 | 1.07E-08 |
| Peak87346 | chr7 | 74561066 | 74561302 | GTF2IRD2B | 52797 | 1.90346116 | 1.07E-08 |
| Peak92617 | chr8 | 81938641 | 81938873 | ZNF704 | -151741 | 3.22356892 | 1.11E-08 |
| Peak92617 | chr8 | 81938641 | 81938873 | PAG1 | 85546 | 3.22356892 | 1.11E-08 |
| Peak88643 | chr7 | 114869551 | 114869700 | MDFIC | 307417 | 2.50387942 | 1.20E-08 |

| | | | | | | | |
|---|---|---|---|---|---|---|---|
| Peak88643 | chr7 | 114869551 | 114869700 | TFEC | 801169 | 2.50387942 | 1.20E-08 |
| Peak50360 | chr2 | 97403663 | 97403900 | KANSL3 | -99736 | 3.30378105 | 1.25E-08 |
| Peak50360 | chr2 | 97403663 | 97403900 | LMAN2L | 2019 | 3.30378105 | 1.25E-08 |
| Peak56343 | chr20 | 31097469 | 31097808 | ASXL1 | 151484 | 1.90033171 | 1.38E-08 |
| Peak56343 | chr20 | 31097469 | 31097808 | COMMD7 | 234164 | 1.90033171 | 1.38E-08 |
| Peak77223 | chr5 | 172137796 | 172137991 | DUSP1 | 60304 | 5.48322418 | 1.46E-08 |
| Peak77223 | chr5 | 172137796 | 172137991 | NEURL1B | 69625 | 5.48322418 | 1.46E-08 |
| Peak8003 | chr1 | 207644315 | 207644550 | CR1 | -25180 | 2.92705862 | 1.52E-08 |
| Peak8003 | chr1 | 207644315 | 207644550 | CR2 | 16858 | 2.92705862 | 1.52E-08 |
| Peak48307 | chr2 | 36688517 | 36688747 | CRIM1 | 105563 | 2.03347516 | 1.58E-08 |
| Peak48307 | chr2 | 36688517 | 36688747 | FEZ2 | 136702 | 2.03347516 | 1.58E-08 |
| Peak52546 | chr2 | 169346544 | 169346742 | NOSTRIN | -308018 | 2.54095528 | 1.59E-08 |
| Peak52546 | chr2 | 169346544 | 169346742 | CERS6 | 33884 | 2.54095528 | 1.59E-08 |
| Peak79783 | chr6 | 45123882 | 45124233 | RUNX2 | -172048 | 2.08779088 | 1.59E-08 |
| Peak79783 | chr6 | 45123882 | 45124233 | CDC5L | 768796 | 2.08779088 | 1.59E-08 |
| Peak22643 | chr12 | 89888593 | 89889015 | DUSP6 | -141756 | 1.11759038 | 1.62E-08 |
| Peak22643 | chr12 | 89888593 | 89889015 | GALNT4 | 29779 | 1.11759038 | 1.62E-08 |
| Peak80761 | chr6 | 90866670 | 90866870 | BACH2 | 139691 | 2.65973955 | 1.76E-08 |
| Peak80761 | chr6 | 90866670 | 90866870 | GJA10 | 262582 | 2.65973955 | 1.76E-08 |
| Peak26589 | chr13 | 86556390 | 86556577 | SLITRK6 | -182861 | 3.08472297 | 1.80E-08 |
| Peak23445 | chr12 | 107141654 | 107141990 | RIC8B | -26596 | 2.32215462 | 1.82E-08 |
| Peak23445 | chr12 | 107141654 | 107141990 | RFX4 | 146907 | 2.32215462 | 1.82E-08 |
| Peak43092 | chr18 | 73120910 | 73121132 | SMIM21 | 18637 | 1.75145807 | 1.93E-08 |
| Peak43092 | chr18 | 73120910 | 73121132 | TSHZ1 | 198311 | 1.75145807 | 1.93E-08 |
| Peak7194 | chr1 | 192873647 | 192873962 | RGS2 | 95634 | 2.14964031 | 1.94E-08 |
| Peak7194 | chr1 | 192873647 | 192873962 | UCHL5 | 154821 | 2.14964031 | 1.94E-08 |
| Peak57808 | chr21 | 16583891 | 16584106 | USP25 | -518345 | 4.4826315 | 1.95E-08 |
| Peak57808 | chr21 | 16583891 | 16584106 | NRIP1 | -146678 | 4.4826315 | 1.95E-08 |
| Peak15119 | chr11 | 14057177 | 14057367 | RRAS2 | 323458 | 3.86094968 | 2.02E-08 |
| Peak15119 | chr11 | 14057177 | 14057367 | FAR1 | 367055 | 3.86094968 | 2.02E-08 |
| Peak41842 | chr18 | 21552399 | 21552677 | CABYR | -166404 | 1.9838336 | 2.03E-08 |
| Peak41842 | chr18 | 21552399 | 21552677 | LAMA3 | 283131 | 1.9838336 | 2.03E-08 |
| Peak76814 | chr5 | 156599421 | 156599775 | ITK | -8239 | 1.57638755 | 2.14E-08 |
| Peak76814 | chr5 | 156599421 | 156599775 | FAM71B | -6323 | 1.57638755 | 2.14E-08 |
| Peak61560 | chr3 | 27947186 | 27947512 | CMC1 | -335737 | 1.39735313 | 2.27E-08 |
| Peak61560 | chr3 | 27947186 | 27947512 | EOMES | -183360 | 1.39735313 | 2.27E-08 |
| Peak101889 | chrX | 131756307 | 131756537 | MBNL3 | -182704 | 1.39541268 | 2.32E-08 |
| Peak101889 | chrX | 131756307 | 131756537 | HS6ST2 | 339001 | 1.39541268 | 2.32E-08 |
| Peak12982 | chr10 | 97833419 | 97833879 | CCNJ | 30309 | 1.91414027 | 2.33E-08 |
| Peak12982 | chr10 | 97833419 | 97833879 | BLNK | 197648 | 1.91414027 | 2.33E-08 |
| Peak61548 | chr3 | 27869452 | 27869669 | CMC1 | -413525 | 3.06385135 | 2.34E-08 |
| Peak61548 | chr3 | 27869452 | 27869669 | EOMES | -105572 | 3.06385135 | 2.34E-08 |
| Peak19042 | chr11 | 128346640 | 128346965 | ETS1 | 110634 | 1.50839755 | 2.37E-08 |
| Peak35323 | chr16 | 47766488 | 47766684 | PHKB | 271348 | 2.82525854 | 2.40E-08 |
| Peak35323 | chr16 | 47766488 | 47766684 | ABCC12 | 414095 | 2.82525854 | 2.40E-08 |
| Peak72350 | chr5 | 10962292 | 10962513 | DAP | -201019 | 2.95578818 | 2.52E-08 |
| Peak72350 | chr5 | 10962292 | 10962513 | CTNND2 | 941752 | 2.95578818 | 2.52E-08 |
| Peak15639 | chr11 | 34183247 | 34183693 | NAT10 | 56321 | 1.81207276 | 2.55E-08 |

| | | | | | | | |
|---|---|---|---|---|---|---|---|
| Peak15639 | chr11 | 34183247 | 34183693 | ABTB2 | 196085 | 1.81207276 | 2.55E-08 |
| Peak80433 | chr6 | 80492571 | 80492833 | SH3BGRL2 | 151702 | 2.27999336 | 2.60E-08 |
| Peak80433 | chr6 | 80492571 | 80492833 | ELOVL4 | 164595 | 2.27999336 | 2.60E-08 |
| Peak19969 | chr12 | 11699052 | 11699271 | PRB2 | -150663 | 1.38303114 | 2.61E-08 |
| Peak19969 | chr12 | 11699052 | 11699271 | ETV6 | -103626 | 1.38303114 | 2.61E-08 |
| Peak8930 | chr1 | 232109625 | 232109934 | DISC1 | 347219 | 2.04279652 | 2.67E-08 |
| Peak8930 | chr1 | 232109625 | 232109934 | SIPA1L2 | 587524 | 2.04279652 | 2.67E-08 |
| Peak75424 | chr5 | 125839431 | 125839782 | ALDH7A1 | 91503 | 4.34151725 | 2.69E-08 |
| Peak75424 | chr5 | 125839431 | 125839782 | GRAMD3 | 143783 | 4.34151725 | 2.69E-08 |
| Peak65245 | chr3 | 141271925 | 141272264 | RNF7 | -184951 | 1.32871327 | 2.77E-08 |
| Peak65245 | chr3 | 141271925 | 141272264 | RASA2 | 66204 | 1.32871327 | 2.77E-08 |
| Peak52183 | chr2 | 159950875 | 159951046 | TANC1 | 125778 | 3.20190878 | 2.77E-08 |
| Peak52183 | chr2 | 159950875 | 159951046 | WDSUB1 | 192142 | 3.20190878 | 2.77E-08 |
| Peak36821 | chr16 | 89600931 | 89601141 | RPL13 | -26083 | 3.15558364 | 2.77E-08 |
| Peak36821 | chr16 | 89600931 | 89601141 | SPG7 | 26225 | 3.15558364 | 2.77E-08 |
| Peak75495 | chr5 | 127156707 | 127157064 | SLC12A2 | -262572 | 2.2900442 | 2.78E-08 |
| Peak75495 | chr5 | 127156707 | 127157064 | CTXN3 | 172150 | 2.2900442 | 2.78E-08 |
| Peak78480 | chr6 | 20202250 | 20202428 | MBOAT1 | 10291 | 1.99021983 | 2.85E-08 |
| Peak78480 | chr6 | 20202250 | 20202428 | ID4 | 364722 | 1.99021983 | 2.85E-08 |
| Peak57883 | chr21 | 19171842 | 19172098 | CHODL | -445180 | 1.25676265 | 2.86E-08 |
| Peak57883 | chr21 | 19171842 | 19172098 | BTG3 | -186808 | 1.25676265 | 2.86E-08 |
| Peak68715 | chr4 | 55808576 | 55808680 | KDR | 183128 | 2.97538324 | 2.87E-08 |
| Peak68715 | chr4 | 55808576 | 55808680 | KIT | 284543 | 2.97538324 | 2.87E-08 |
| Peak86027 | chr7 | 27175623 | 27175782 | HOXA4 | -5285 | 2.65178048 | 2.89E-08 |
| Peak86027 | chr7 | 27175623 | 27175782 | HOXA5 | 7584 | 2.65178048 | 2.89E-08 |
| Peak66774 | chr3 | 187907722 | 187907969 | BCL6 | -444331 | 2.61346821 | 2.91E-08 |
| Peak66774 | chr3 | 187907722 | 187907969 | LPP | -22875 | 2.61346821 | 2.91E-08 |
| Peak32677 | chr15 | 73256930 | 73257173 | ADPGK | -180926 | 2.65492199 | 2.93E-08 |
| Peak32677 | chr15 | 73256930 | 73257173 | NEO1 | -86999 | 2.65492199 | 2.93E-08 |
| Peak10318 | chr10 | 16990361 | 16990728 | RSU1 | -131163 | 1.4955089 | 2.98E-08 |
| Peak10318 | chr10 | 16990361 | 16990728 | CUBN | 181285 | 1.4955089 | 2.98E-08 |
| Peak49304 | chr2 | 65547163 | 65547543 | ACTR2 | 92382 | 1.14881124 | 3.01E-08 |
| Peak49304 | chr2 | 65547163 | 65547543 | SPRED2 | 111958 | 1.14881124 | 3.01E-08 |
| Peak37677 | chr17 | 14262633 | 14263015 | HS3ST3B1 | 58424 | 1.13480337 | 3.04E-08 |
| Peak37677 | chr17 | 14262633 | 14263015 | PMP22 | 903082 | 1.13480337 | 3.04E-08 |
| Peak77901 | chr6 | 4011124 | 4011330 | PRPF4B | -10333 | 1.81154584 | 3.06E-08 |
| Peak77901 | chr6 | 4011124 | 4011330 | ENSG00000 | 28318 | 1.81154584 | 3.06E-08 |
| Peak72648 | chr5 | 33236032 | 33236224 | TARS | -204943 | 3.25390225 | 3.12E-08 |
| Peak72648 | chr5 | 33236032 | 33236224 | NPR3 | 524588 | 3.25390225 | 3.12E-08 |
| Peak32027 | chr15 | 58501376 | 58501727 | LIPC | -201216 | 1.81445976 | 3.27E-08 |
| Peak32027 | chr15 | 58501376 | 58501727 | AQP9 | 71157 | 1.81445976 | 3.27E-08 |
| Peak93171 | chr8 | 101527435 | 101527859 | RNF19A | -212160 | 1.7526167 | 3.38E-08 |
| Peak93171 | chr8 | 101527435 | 101527859 | ANKRD46 | 44323 | 1.7526167 | 3.38E-08 |
| Peak10307 | chr10 | 16877341 | 16877535 | RSU1 | -18056 | 2.05730257 | 3.53E-08 |
| Peak10307 | chr10 | 16877341 | 16877535 | CUBN | 294392 | 2.05730257 | 3.53E-08 |
| Peak26390 | chr13 | 76444178 | 76444444 | LMO7 | 109514 | 2.05275762 | 3.58E-08 |
| Peak20699 | chr12 | 32287438 | 32287800 | FGD4 | -367475 | 1.20349152 | 3.61E-08 |
| Peak20699 | chr12 | 32287438 | 32287800 | BICD1 | 27456 | 1.20349152 | 3.61E-08 |

| | | | | | | | |
|---|---|---|---|---|---|---|---|
| Peak11006 | chr10 | 33537587 | 33537884 | ITGB1 | -290914 | 1.22293454 | 3.67E-08 |
| Peak11006 | chr10 | 33537587 | 33537884 | NRP1 | 87454 | 1.22293454 | 3.67E-08 |
| Peak20215 | chr12 | 19205845 | 19205964 | PLEKHA5 | -76797 | 3.25260201 | 3.83E-08 |
| Peak20215 | chr12 | 19205845 | 19205964 | CAPZA3 | 314860 | 3.25260201 | 3.83E-08 |
| Peak35543 | chr16 | 56713849 | 56714053 | MT1X | -2385 | 2.51586816 | 3.83E-08 |
| Peak4981 | chr1 | 144891805 | 144892034 | NBPF9 | 80172 | 2.15095879 | 3.86E-08 |
| Peak4981 | chr1 | 144891805 | 144892034 | PDE4DIP | 103102 | 2.15095879 | 3.86E-08 |
| Peak34568 | chr16 | 21577238 | 21577542 | NPIPB3 | -146312 | 2.11741552 | 3.88E-08 |
| Peak34568 | chr16 | 21577238 | 21577542 | METTL9 | -33407 | 2.11741552 | 3.88E-08 |
| Peak15646 | chr11 | 34272639 | 34272978 | ABTB2 | 106746 | 1.05192469 | 3.91E-08 |
| Peak15646 | chr11 | 34272639 | 34272978 | NAT10 | 145660 | 1.05192469 | 3.91E-08 |
| Peak89437 | chr7 | 139467396 | 139467556 | HIPK2 | 10041 | 2.12594742 | 4.27E-08 |
| Peak89437 | chr7 | 139467396 | 139467556 | CLEC2L | 258874 | 2.12594742 | 4.27E-08 |
| Peak18227 | chr11 | 110216495 | 110216755 | FDX1 | -83982 | 2.57497708 | 4.34E-08 |
| Peak18227 | chr11 | 110216495 | 110216755 | RDX | -49188 | 2.57497708 | 4.34E-08 |
| Peak70620 | chr4 | 124317801 | 124318017 | SPRY1 | -2756 | 1.32186114 | 4.35E-08 |
| Peak93978 | chr8 | 128209154 | 128209432 | FAM84B | -638655 | 2.02502559 | 4.42E-08 |
| Peak93978 | chr8 | 128209154 | 128209432 | POU5F1B | -217242 | 2.02502559 | 4.42E-08 |
| Peak63995 | chr3 | 108061843 | 108061967 | HHLA2 | 40573 | 3.68262048 | 4.47E-08 |
| Peak63995 | chr3 | 108061843 | 108061967 | MYH15 | 186264 | 3.68262048 | 4.47E-08 |
| Peak52717 | chr2 | 173117993 | 173118225 | ITGA6 | -174408 | 2.22009149 | 4.53E-08 |
| Peak52717 | chr2 | 173117993 | 173118225 | DLX2 | -150481 | 2.22009149 | 4.53E-08 |
| Peak51950 | chr2 | 149634434 | 149634607 | LYPD6B | -260460 | 2.08598646 | 4.54E-08 |
| Peak51950 | chr2 | 149634434 | 149634607 | KIF5C | 1702 | 2.08598646 | 4.54E-08 |
| Peak70280 | chr4 | 114623873 | 114624060 | CAMK2D | 58257 | 2.8849941 | 4.57E-08 |
| Peak70280 | chr4 | 114623873 | 114624060 | ANK2 | 653135 | 2.8849941 | 4.57E-08 |
| Peak97401 | chr9 | 107769448 | 107769706 | SLC44A1 | -237326 | 2.2841331 | 4.66E-08 |
| Peak97401 | chr9 | 107769448 | 107769706 | ABCA1 | -79059 | 2.2841331 | 4.66E-08 |
| Peak64247 | chr3 | 112721623 | 112721870 | GTPBP8 | 11982 | 1.22549905 | 4.67E-08 |
| Peak64247 | chr3 | 112721623 | 112721870 | C3orf17 | 16939 | 1.22549905 | 4.67E-08 |
| Peak53715 | chr2 | 201090791 | 201090968 | TYW5 | -270421 | 3.74706675 | 4.70E-08 |
| Peak53715 | chr2 | 201090791 | 201090968 | SPATS2L | -79994 | 3.74706675 | 4.70E-08 |
| Peak64451 | chr3 | 119281462 | 119281767 | CD80 | -3166 | 1.66098897 | 4.75E-08 |
| Peak82629 | chr6 | 139700070 | 139700184 | CITED2 | -4370 | 1.9299825 | 4.87E-08 |
| Peak80652 | chr6 | 88442163 | 88442285 | SPACA1 | -315283 | 3.17744791 | 4.92E-08 |
| Peak80652 | chr6 | 88442163 | 88442285 | AKIRIN2 | -30297 | 3.17744791 | 4.92E-08 |
| Peak95104 | chr9 | 7879598 | 7879864 | C9orf123 | -79664 | 1.56401511 | 4.93E-08 |
| Peak18263 | chr11 | 111410181 | 111410380 | LAYN | -1103 | 3.25757472 | 5.04E-08 |
| Peak80227 | chr6 | 71938166 | 71938765 | B3GAT2 | -271725 | 1.83216029 | 5.15E-08 |
| Peak80227 | chr6 | 71938166 | 71938765 | OGFRL1 | -60040 | 1.83216029 | 5.15E-08 |
| Peak26602 | chr13 | 87234069 | 87234211 | SLITRK6 | -860517 | 2.99959152 | 5.17E-08 |
| Peak35087 | chr16 | 30578588 | 30578838 | ENSG00000[illegible] | -9113 | 1.76488013 | 5.21E-08 |
| Peak35087 | chr16 | 30578588 | 30578838 | ZNF688 | 5342 | 1.76488013 | 5.21E-08 |
| Peak53657 | chr2 | 198734347 | 198734636 | PLCL1 | 65066 | 2.80997989 | 5.21E-08 |
| Peak48329 | chr2 | 37116954 | 37117327 | STRN | 76474 | 1.49013602 | 5.57E-08 |
| Peak48329 | chr2 | 37116954 | 37117327 | VIT | 193308 | 1.49013602 | 5.57E-08 |
| Peak2967 | chr1 | 67788884 | 67789065 | IL12RB2 | 15928 | 2.52352727 | 5.60E-08 |
| Peak2967 | chr1 | 67788884 | 67789065 | SERBP1 | 107094 | 2.52352727 | 5.60E-08 |

| | | | | | | | |
|---|---|---|---|---|---|---|---|
| Peak92569 | chr8 | 81083667 | 81083984 | ZBTB10 | -314076 | 1.22727014 | 5.62E-08 |
| Peak92569 | chr8 | 81083667 | 81083984 | TPD52 | -90775 | 1.22727014 | 5.62E-08 |
| Peak33182 | chr15 | 85567985 | 85568197 | AKAP13 | -355869 | 2.79342758 | 5.87E-08 |
| Peak33182 | chr15 | 85567985 | 85568197 | PDE8A | 44420 | 2.79342758 | 5.87E-08 |
| Peak2943 | chr1 | 67625028 | 67625199 | SLC35D1 | -105332 | 2.09142977 | 5.90E-08 |
| Peak2943 | chr1 | 67625028 | 67625199 | IL23R | -6969 | 2.09142977 | 5.90E-08 |
| Peak47337 | chr2 | 8595870 | 8596014 | ID2 | -223033 | 2.38959583 | 6.00E-08 |
| Peak69085 | chr4 | 76608219 | 76608333 | USO1 | -41728 | 2.33874505 | 6.03E-08 |
| Peak69085 | chr4 | 76608219 | 76608333 | G3BP2 | -9123 | 2.33874505 | 6.03E-08 |
| Peak8583 | chr1 | 226053805 | 226054061 | TMEM63A | 16136 | 1.74807222 | 6.11E-08 |
| Peak8583 | chr1 | 226053805 | 226054061 | EPHX1 | 40878 | 1.74807222 | 6.11E-08 |
| Peak53998 | chr2 | 205889866 | 205890151 | NRP2 | -657215 | 1.7840105 | 6.23E-08 |
| Peak53998 | chr2 | 205889866 | 205890151 | PARD3B | 479286 | 1.7840105 | 6.23E-08 |
| Peak73387 | chr5 | 56413449 | 56413819 | MIER3 | -165682 | 2.6173284 | 6.29E-08 |
| Peak73387 | chr5 | 56413449 | 56413819 | GPBP1 | -96314 | 2.6173284 | 6.29E-08 |
| Peak94636 | chr8 | 145317445 | 145317612 | SCXB | -3988 | 3.24236192 | 6.63E-08 |
| Peak52141 | chr2 | 158320787 | 158321077 | CYTIP | -20278 | 1.54665795 | 6.67E-08 |
| Peak52141 | chr2 | 158320787 | 158321077 | ACVR1C | 164585 | 1.54665795 | 6.67E-08 |
| Peak69927 | chr4 | 103551889 | 103552151 | NFKB1 | 129534 | 2.05289634 | 6.67E-08 |
| Peak69927 | chr4 | 103551889 | 103552151 | MANBA | 130131 | 2.05289634 | 6.67E-08 |
| Peak64056 | chr3 | 109504567 | 109504780 | DPPA4 | -448255 | 2.81993855 | 6.79E-08 |
| Peak80544 | chr6 | 84139096 | 84139213 | PGM3 | -235522 | 3.70389592 | 6.79E-08 |
| Peak80544 | chr6 | 84139096 | 84139213 | ME1 | 1635 | 3.70389592 | 6.79E-08 |
| Peak51670 | chr2 | 136970381 | 136970739 | THSD7B | -669189 | 1.4719342 | 7.07E-08 |
| Peak51670 | chr2 | 136970381 | 136970739 | CXCR4 | -96747 | 1.4719342 | 7.07E-08 |
| Peak63451 | chr3 | 87255289 | 87255596 | VGLL3 | -215186 | 2.15144247 | 7.14E-08 |
| Peak63451 | chr3 | 87255289 | 87255596 | CHMP2B | -20992 | 2.15144247 | 7.14E-08 |
| Peak61664 | chr3 | 31503558 | 31504098 | GADL1 | -567571 | 1.33661127 | 7.17E-08 |
| Peak61664 | chr3 | 31503558 | 31504098 | STT3B | -70454 | 1.33661127 | 7.17E-08 |
| Peak40118 | chr17 | 63019239 | 63019523 | LRRC37A3 | -104478 | 1.93385898 | 7.19E-08 |
| Peak40118 | chr17 | 63019239 | 63019523 | GNA13 | 33576 | 1.93385898 | 7.19E-08 |
| Peak4980 | chr1 | 144887665 | 144887916 | NBPF9 | 76043 | 3.2612362 | 7.25E-08 |
| Peak4980 | chr1 | 144887665 | 144887916 | PDE4DIP | 107231 | 3.2612362 | 7.25E-08 |
| Peak62259 | chr3 | 46029973 | 46030142 | FYCO1 | 7249 | 2.99740595 | 7.31E-08 |
| Peak62259 | chr3 | 46029973 | 46030142 | CXCR6 | 43547 | 2.99740595 | 7.31E-08 |
| Peak62878 | chr3 | 58200553 | 58200762 | DNASE1L3 | -260 | 2.15691807 | 7.49E-08 |
| Peak79802 | chr6 | 45401888 | 45402067 | SUPT3H | -56308 | 2.88582139 | 8.16E-08 |
| Peak79802 | chr6 | 45401888 | 45402067 | CLIC5 | 646154 | 2.88582139 | 8.16E-08 |
| Peak11398 | chr10 | 50488166 | 50488428 | C10orf128 | -91893 | 1.92505256 | 8.41E-08 |
| Peak11398 | chr10 | 50488166 | 50488428 | DRGX | 111610 | 1.92505256 | 8.41E-08 |
| Peak77052 | chr5 | 163344874 | 163345064 | MAT2B | 412415 | 3.04512108 | 8.60E-08 |
| Peak28184 | chr14 | 50449528 | 50449674 | ARF6 | 89791 | 2.98733668 | 8.61E-08 |
| Peak28184 | chr14 | 50449528 | 50449674 | METTL21D | 133674 | 2.98733668 | 8.61E-08 |
| Peak82649 | chr6 | 140363167 | 140363459 | CITED2 | -667556 | 3.83288057 | 8.61E-08 |
| Peak96403 | chr9 | 75201352 | 75201967 | TMC1 | 64943 | 2.06236377 | 8.62E-08 |
| Peak96403 | chr9 | 75201352 | 75201967 | ALDH1A1 | 366311 | 2.06236377 | 8.62E-08 |
| Peak48610 | chr2 | 43411594 | 43411822 | HAAO | -391976 | 1.70334218 | 8.65E-08 |
| Peak48610 | chr2 | 43411594 | 43411822 | ZFP36L2 | 42040 | 1.70334218 | 8.65E-08 |

| | | | | | | | |
|---|---|---|---|---|---|---|---|
| Peak48889 | chr2 | 54268327 | 54268726 | ACYP2 | -74114 | 1.20560616 | 8.73E-08 |
| Peak48889 | chr2 | 54268327 | 54268726 | PSME4 | -70550 | 1.20560616 | 8.73E-08 |
| Peak15642 | chr11 | 34236497 | 34236741 | NAT10 | 109470 | 2.12427292 | 8.77E-08 |
| Peak15642 | chr11 | 34236497 | 34236741 | ABTB2 | 142936 | 2.12427292 | 8.77E-08 |
| Peak23032 | chr12 | 96190573 | 96190772 | SNRPF | -62033 | 2.91859358 | 8.90E-08 |
| Peak23032 | chr12 | 96190573 | 96190772 | NTN4 | -6137 | 2.91859358 | 8.90E-08 |
| Peak42093 | chr18 | 33186066 | 33186314 | INO80C | -108283 | 1.6775462 | 9.19E-08 |
| Peak42093 | chr18 | 33186066 | 33186314 | GALNT1 | -48334 | 1.6775462 | 9.19E-08 |
| Peak10797 | chr10 | 30101122 | 30101376 | SVIL | -76519 | 2.57819371 | 9.29E-08 |
| Peak10797 | chr10 | 30101122 | 30101376 | KIAA1462 | 247204 | 2.57819371 | 9.29E-08 |
| Peak42309 | chr18 | 43895167 | 43895404 | RNF165 | -18901 | 2.96480041 | 9.31E-08 |
| Peak42309 | chr18 | 43895167 | 43895404 | C18orf25 | 141293 | 2.96480041 | 9.31E-08 |
| Peak41627 | chr18 | 13436732 | 13436899 | LDLRAD4 | 218030 | 3.20823826 | 9.57E-08 |
| Peak41627 | chr18 | 13436732 | 13436899 | FAM210A | 289743 | 3.20823826 | 9.57E-08 |
| Peak92233 | chr8 | 67426140 | 67426559 | ADHFE1 | 81629 | 3.2990444 | 9.88E-08 |
| Peak92233 | chr8 | 67426140 | 67426559 | MYBL1 | 99134 | 3.2990444 | 9.88E-08 |
| Peak70632 | chr4 | 124401123 | 124401417 | SPRY1 | 80605 | 3.26711866 | 1.01E-07 |
| Peak80553 | chr6 | 84256412 | 84256667 | PRSS35 | 34346 | 2.63696662 | 1.01E-07 |
| Peak80553 | chr6 | 84256412 | 84256667 | SNAP91 | 162587 | 2.63696662 | 1.01E-07 |
| Peak3956 | chr1 | 100828549 | 100828729 | GPR88 | -175054 | 3.96069517 | 1.08E-07 |
| Peak3956 | chr1 | 100828549 | 100828729 | CDC14A | 10592 | 3.96069517 | 1.08E-07 |
| Peak28753 | chr14 | 62036674 | 62036843 | HIF1A | -127581 | 3.01780642 | 1.09E-07 |
| Peak28753 | chr14 | 62036674 | 62036843 | PRKCH | 248324 | 3.01780642 | 1.09E-07 |
| Peak92632 | chr8 | 82009486 | 82010026 | ZNF704 | -222740 | 1.6493223 | 1.09E-07 |
| Peak92632 | chr8 | 82009486 | 82010026 | PAG1 | 14547 | 1.6493223 | 1.09E-07 |
| Peak94088 | chr8 | 130900533 | 130900794 | GSDMC | -101530 | 1.42907741 | 1.10E-07 |
| Peak94088 | chr8 | 130900533 | 130900794 | FAM49B | 51414 | 1.42907741 | 1.10E-07 |
| Peak53208 | chr2 | 190126217 | 190126607 | WDR75 | -179747 | 3.20474547 | 1.11E-07 |
| Peak53208 | chr2 | 190126217 | 190126607 | COL5A2 | -81807 | 3.20474547 | 1.11E-07 |
| Peak82524 | chr6 | 138303520 | 138303970 | TNFAIP3 | 115164 | 1.50775829 | 1.14E-07 |
| Peak82524 | chr6 | 138303520 | 138303970 | PERP | 124903 | 1.50775829 | 1.14E-07 |
| Peak71215 | chr4 | 153021471 | 153021721 | PET112 | -339421 | 1.53869234 | 1.16E-07 |
| Peak71215 | chr4 | 153021471 | 153021721 | FBXW7 | 435657 | 1.53869234 | 1.16E-07 |
| Peak68714 | chr4 | 55808196 | 55808462 | KDR | 183427 | 2.56943034 | 1.17E-07 |
| Peak68714 | chr4 | 55808196 | 55808462 | KIT | 284244 | 2.56943034 | 1.17E-07 |
| Peak26010 | chr13 | 52220629 | 52220733 | WDFY2 | 62037 | 3.06761395 | 1.20E-07 |
| Peak26010 | chr13 | 52220629 | 52220733 | DHRS12 | 157578 | 3.06761395 | 1.20E-07 |
| Peak62789 | chr3 | 56702028 | 56702501 | CCDC66 | 111064 | 3.44469472 | 1.22E-07 |
| Peak62789 | chr3 | 56702028 | 56702501 | ARHGEF3 | 411071 | 3.44469472 | 1.22E-07 |
| Peak95653 | chr9 | 33417145 | 33417289 | AQP7 | -14700 | 2.79563488 | 1.23E-07 |
| Peak95653 | chr9 | 33417145 | 33417289 | AQP3 | 30392 | 2.79563488 | 1.23E-07 |
| Peak60896 | chr3 | 10265449 | 10265687 | TATDN2 | -24139 | 1.25395815 | 1.23E-07 |
| Peak60896 | chr3 | 10265449 | 10265687 | IRAK2 | 59019 | 1.25395815 | 1.23E-07 |
| Peak53001 | chr2 | 179379428 | 179379805 | PLEKHA3 | 34422 | 2.69808511 | 1.24E-07 |
| Peak53001 | chr2 | 179379428 | 179379805 | TTN | 292533 | 2.69808511 | 1.24E-07 |
| Peak95654 | chr9 | 33426753 | 33426811 | AQP7 | -24265 | 4.88623771 | 1.24E-07 |
| Peak95654 | chr9 | 33426753 | 33426811 | AQP3 | 20827 | 4.88623771 | 1.24E-07 |
| Peak13587 | chr10 | 112295916 | 112296253 | SMC3 | -31364 | 1.34196941 | 1.32E-07 |

| | | | | | | | |
|---|---|---|---|---|---|---|---|
| Peak13587 | chr10 | 112295916 | 112296253 | DUSP5 | 38489 | 1.34196941 | 1.32E-07 |
| Peak68755 | chr4 | 56732558 | 56732806 | CEP135 | -82465 | 1.14011784 | 1.32E-07 |
| Peak68755 | chr4 | 56732558 | 56732806 | EXOC1 | 12900 | 1.14011784 | 1.32E-07 |
| Peak7062 | chr1 | 185256685 | 185256871 | TRMT1L | -130665 | 3.0129935 | 1.32E-07 |
| Peak7062 | chr1 | 185256685 | 185256871 | IVNS1ABP | 29683 | 3.0129935 | 1.32E-07 |
| Peak3341 | chr1 | 86805139 | 86805426 | COL24A1 | -182837 | 2.27639481 | 1.36E-07 |
| Peak3341 | chr1 | 86805139 | 86805426 | ODF2L | 56662 | 2.27639481 | 1.36E-07 |
| Peak61567 | chr3 | 28092392 | 28092710 | EOMES | -328562 | 1.38361594 | 1.38E-07 |
| Peak61567 | chr3 | 28092392 | 28092710 | CMC1 | -190535 | 1.38361594 | 1.38E-07 |
| Peak68699 | chr4 | 55524727 | 55524812 | KIT | 685 | 4.43681189 | 1.42E-07 |
| Peak13500 | chr10 | 111773491 | 111773725 | MXI1 | -193755 | 1.70503473 | 1.49E-07 |
| Peak13500 | chr10 | 111773491 | 111773725 | ADD3 | 5888 | 1.70503473 | 1.49E-07 |
| Peak21978 | chr12 | 65078858 | 65079111 | GNS | 74242 | 2.42154217 | 1.49E-07 |
| Peak21978 | chr12 | 65078858 | 65079111 | RASSF3 | 74692 | 2.42154217 | 1.49E-07 |
| Peak88622 | chr7 | 114571839 | 114572036 | MDFIC | 9729 | 1.30103405 | 1.49E-07 |
| Peak73723 | chr5 | 67678594 | 67678813 | SLC30A5 | -710769 | 2.08448521 | 1.54E-07 |
| Peak73723 | chr5 | 67678594 | 67678813 | PIK3R1 | 167156 | 2.08448521 | 1.54E-07 |
| Peak10581 | chr10 | 25154041 | 25154157 | ARHGAP21 | -141502 | 2.79694522 | 1.58E-07 |
| Peak10581 | chr10 | 25154041 | 25154157 | PRTFDC1 | 87434 | 2.79694522 | 1.58E-07 |
| Peak47598 | chr2 | 12696839 | 12697208 | TRIB2 | -159991 | 1.67044193 | 1.59E-07 |
| Peak47598 | chr2 | 12696839 | 12697208 | LPIN1 | 879303 | 1.67044193 | 1.59E-07 |
| Peak33838 | chr16 | 1758960 | 1759118 | MAPK8IP3 | 2855 | 2.61955765 | 1.60E-07 |
| Peak33838 | chr16 | 1758960 | 1759118 | NME3 | 62692 | 2.61955765 | 1.60E-07 |
| Peak53727 | chr2 | 201241298 | 201241408 | SPATS2L | 70479 | 2.72686197 | 1.61E-07 |
| Peak53727 | chr2 | 201241298 | 201241408 | KCTD18 | 133433 | 2.72686197 | 1.61E-07 |
| Peak102007 | chrX | 135709617 | 135709875 | CD40LG | -20606 | 1.766424 | 1.62E-07 |
| Peak102007 | chrX | 135709617 | 135709875 | VGLL1 | 95435 | 1.766424 | 1.62E-07 |
| Peak74777 | chr5 | 99886335 | 99886616 | FAM174A | 15467 | 2.6210097 | 1.65E-07 |
| Peak74777 | chr5 | 99886335 | 99886616 | ST8SIA4 | 352494 | 2.6210097 | 1.65E-07 |
| Peak22569 | chr12 | 85388280 | 85388566 | SLC6A15 | -81768 | 2.69035775 | 1.65E-07 |
| Peak22569 | chr12 | 85388280 | 85388566 | TSPAN19 | 41612 | 2.69035775 | 1.65E-07 |
| Peak7061 | chr1 | 185250373 | 185250769 | TRMT1L | -124458 | 2.38547521 | 1.66E-07 |
| Peak7061 | chr1 | 185250373 | 185250769 | IVNS1ABP | 35890 | 2.38547521 | 1.66E-07 |
| Peak11472 | chr10 | 52263745 | 52264003 | ASAH2 | -255504 | 2.50657264 | 1.68E-07 |
| Peak11472 | chr10 | 52263745 | 52264003 | SGMS1 | 119869 | 2.50657264 | 1.68E-07 |
| Peak73074 | chr5 | 43193755 | 43193979 | ENSG00000[illegible] | 913 | 1.68285074 | 1.69E-07 |
| Peak92263 | chr8 | 67638484 | 67638649 | SGK3 | -48895 | 2.75811732 | 1.70E-07 |
| Peak92263 | chr8 | 67638484 | 67638649 | C8orf44-SGK[illegible] | 58716 | 2.75811732 | 1.70E-07 |
| Peak82246 | chr6 | 134499411 | 134499678 | SLC2A12 | -125771 | 1.42863279 | 1.70E-07 |
| Peak82246 | chr6 | 134499411 | 134499678 | SGK1 | 139651 | 1.42863279 | 1.70E-07 |
| Peak95655 | chr9 | 33426881 | 33427326 | AQP7 | -24587 | 3.30537751 | 1.70E-07 |
| Peak95655 | chr9 | 33426881 | 33427326 | AQP3 | 20505 | 3.30537751 | 1.70E-07 |
| Peak92884 | chr8 | 94371221 | 94371477 | TRIQK | -392976 | 3.6894538 | 1.72E-07 |
| Peak92884 | chr8 | 94371221 | 94371477 | RBM12B | 381896 | 3.6894538 | 1.72E-07 |
| Peak94642 | chr8 | 145486420 | 145486625 | SCXA | -4026 | 3.6681779 | 1.73E-07 |
| Peak89369 | chr7 | 138482560 | 138482752 | TMEM213 | -41 | 3.01925962 | 1.75E-07 |
| Peak89369 | chr7 | 138482560 | 138482752 | ATP6V0A4 | 285 | 3.01925962 | 1.75E-07 |
| Peak78085 | chr6 | 9624431 | 9624630 | OFCC1 | 315021 | 3.06399041 | 1.76E-07 |

| | | | | | | | |
|---|---|---|---|---|---|---|---|
| Peak28489 | chr14 | 55538710 | 55539016 | LGALS3 | -56899 | 1.51064802 | 1.80E-07 |
| Peak28489 | chr14 | 55538710 | 55539016 | SOCS4 | 44915 | 1.51064802 | 1.80E-07 |
| Peak14960 | chr11 | 10477447 | 10477678 | AMPD3 | 5335 | 1.09204603 | 1.81E-07 |
| Peak14960 | chr11 | 10477447 | 10477678 | MTRNR2L8 | 53160 | 1.09204603 | 1.81E-07 |
| Peak100731 | chrX | 65086035 | 65086345 | VSIG4 | 173759 | 1.89561053 | 1.83E-07 |
| Peak100731 | chrX | 65086035 | 65086345 | MSN | 198653 | 1.89561053 | 1.83E-07 |
| Peak28819 | chr14 | 64330773 | 64330977 | SYNE2 | 11192 | 1.13281175 | 1.85E-07 |
| Peak28819 | chr14 | 64330773 | 64330977 | ESR2 | 430203 | 1.13281175 | 1.85E-07 |
| Peak42829 | chr18 | 60976834 | 60976984 | BCL2 | 10452 | 2.11359914 | 1.85E-07 |
| Peak42829 | chr18 | 60976834 | 60976984 | PHLPP1 | 594226 | 2.11359914 | 1.85E-07 |
| Peak11835 | chr10 | 65467075 | 65467411 | REEP3 | 186120 | 1.46969683 | 1.85E-07 |
| Peak23533 | chr12 | 108962664 | 108962938 | ISCU | 6424 | 2.2259293 | 1.88E-07 |
| Peak23533 | chr12 | 108962664 | 108962938 | TMEM119 | 29099 | 2.2259293 | 1.88E-07 |
| Peak70643 | chr4 | 124538136 | 124538387 | SPRY1 | 217597 | 1.98196954 | 1.89E-07 |
| Peak43022 | chr18 | 71950493 | 71950680 | CYB5A | 8664 | 3.3593078 | 1.93E-07 |
| Peak43022 | chr18 | 71950493 | 71950680 | TIMM21 | 134841 | 3.3593078 | 1.93E-07 |
| Peak53938 | chr2 | 204584862 | 204585063 | CTLA4 | -147546 | 2.17396279 | 1.95E-07 |
| Peak53938 | chr2 | 204584862 | 204585063 | CD28 | 13692 | 2.17396279 | 1.95E-07 |
| Peak12518 | chr10 | 85916602 | 85916877 | C10orf99 | -16754 | 1.47674358 | 1.96E-07 |
| Peak12518 | chr10 | 85916602 | 85916877 | GHITM | 17544 | 1.47674358 | 1.96E-07 |
| Peak86460 | chr7 | 38269652 | 38269759 | STARD3NL | 51882 | 1.47768385 | 1.98E-07 |
| Peak86460 | chr7 | 38269652 | 38269759 | AMPH | 401461 | 1.47768385 | 1.98E-07 |
| Peak74674 | chr5 | 96130481 | 96130936 | ERAP1 | 13094 | 1.01685639 | 2.01E-07 |
| Peak74674 | chr5 | 96130481 | 96130936 | CAST | 132768 | 1.01685639 | 2.01E-07 |
| Peak26691 | chr13 | 94410931 | 94411163 | GPC6 | 531952 | 2.95251345 | 2.04E-07 |
| Peak26691 | chr13 | 94410931 | 94411163 | DCT | 720889 | 2.95251345 | 2.04E-07 |
| Peak42119 | chr18 | 34000224 | 34000463 | FHOD3 | 122667 | 3.03176501 | 2.04E-07 |
| Peak42119 | chr18 | 34000224 | 34000463 | TPGS2 | 408814 | 3.03176501 | 2.04E-07 |
| Peak48576 | chr2 | 42648074 | 42648225 | COX7A2L | -52000 | 2.64446003 | 2.07E-07 |
| Peak48576 | chr2 | 42648074 | 42648225 | KCNG3 | 73087 | 2.64446003 | 2.07E-07 |
| Peak48849 | chr2 | 53163236 | 53163441 | CHAC2 | -831590 | 2.83995142 | 2.07E-07 |
| Peak11966 | chr10 | 71119451 | 71119785 | TACR2 | 57005 | 2.52734093 | 2.09E-07 |
| Peak11966 | chr10 | 71119451 | 71119785 | HK1 | 71118 | 2.52734093 | 2.09E-07 |
| Peak7277 | chr1 | 193603676 | 193603874 | B3GALT2 | -447991 | 1.86721738 | 2.15E-07 |
| Peak2948 | chr1 | 67637748 | 67638038 | IL12RB2 | -135154 | 1.46090888 | 2.17E-07 |
| Peak2948 | chr1 | 67637748 | 67638038 | IL23R | 5810 | 1.46090888 | 2.17E-07 |
| Peak92798 | chr8 | 90914140 | 90914352 | OSGIN2 | -516 | 1.07377306 | 2.17E-07 |
| Peak53659 | chr2 | 198756732 | 198756902 | PLCL1 | 87391 | 3.09053217 | 2.18E-07 |
| Peak70864 | chr4 | 140578920 | 140579190 | SETD7 | -101127 | 1.77887265 | 2.18E-07 |
| Peak70864 | chr4 | 140578920 | 140579190 | MGST2 | -7867 | 1.77887265 | 2.18E-07 |
| Peak62877 | chr3 | 58192492 | 58192638 | DNASE1L3 | 7833 | 3.43682003 | 2.18E-07 |
| Peak62877 | chr3 | 58192492 | 58192638 | FLNB | 198438 | 3.43682003 | 2.18E-07 |
| Peak33568 | chr15 | 99496425 | 99496721 | PGPEP1L | 52218 | 2.22041246 | 2.20E-07 |
| Peak33568 | chr15 | 99496425 | 99496721 | IGF1R | 304373 | 2.22041246 | 2.20E-07 |
| Peak26608 | chr13 | 87417750 | 87418045 | SLITRK5 | -906972 | 3.61365563 | 2.22E-07 |
| Peak75238 | chr5 | 115367821 | 115368120 | COMMD10 | -52717 | 1.89232227 | 2.27E-07 |
| Peak75238 | chr5 | 115367821 | 115368120 | ENSG00000: | 69780 | 1.89232227 | 2.27E-07 |
| Peak79769 | chr6 | 44893224 | 44893495 | RUNX2 | -402746 | 1.41108568 | 2.28E-07 |

| | | | | | | | |
|---|---|---|---|---|---|---|---|
| Peak79769 | chr6 | 44893224 | 44893495 | CDC5L | 538098 | 1.41108568 | 2.28E-07 |
| Peak15726 | chr11 | 35298040 | 35298281 | CD44 | 137433 | 2.85836255 | 2.30E-07 |
| Peak15726 | chr11 | 35298040 | 35298281 | SLC1A2 | 142635 | 2.85836255 | 2.30E-07 |
| Peak91002 | chr8 | 23100988 | 23101184 | CHMP7 | -64 | 1.93616177 | 2.32E-07 |
| Peak33396 | chr15 | 91388509 | 91388778 | FURIN | -23178 | 2.91257695 | 2.37E-07 |
| Peak33396 | chr15 | 91388509 | 91388778 | BLM | 128086 | 2.91257695 | 2.37E-07 |
| Peak96302 | chr9 | 72164225 | 72164547 | APBA1 | 122836 | 2.34824892 | 2.39E-07 |
| Peak96302 | chr9 | 72164225 | 72164547 | FAM189A2 | 220145 | 2.34824892 | 2.39E-07 |
| Peak64103 | chr3 | 111291000 | 111291225 | ZBED2 | 23177 | 1.55714932 | 2.42E-07 |
| Peak64103 | chr3 | 111291000 | 111291225 | CD96 | 30148 | 1.55714932 | 2.42E-07 |
| Peak100064 | chrX | 37705535 | 37705721 | DYNLT3 | 1262 | 1.52000991 | 2.42E-07 |
| Peak100064 | chrX | 37705535 | 37705721 | CYBB | 66364 | 1.52000991 | 2.42E-07 |
| Peak85514 | chr7 | 12296050 | 12296132 | TMEM106B | 45165 | 2.62908874 | 2.50E-07 |
| Peak85514 | chr7 | 12296050 | 12296132 | VWDE | 147440 | 2.62908874 | 2.50E-07 |
| Peak6524 | chr1 | 169877853 | 169878122 | SCYL3 | -14912 | 1.75946408 | 2.52E-07 |
| Peak6524 | chr1 | 169877853 | 169878122 | KIFAP3 | 165848 | 1.75946408 | 2.52E-07 |
| Peak93188 | chr8 | 101821750 | 101822166 | PABPC1 | -87018 | 1.61249957 | 2.54E-07 |
| Peak93188 | chr8 | 101821750 | 101822166 | YWHAZ | 143253 | 1.61249957 | 2.54E-07 |
| Peak82150 | chr6 | 132110393 | 132110702 | CTAGE9 | -78342 | 3.59529844 | 2.54E-07 |
| Peak82150 | chr6 | 132110393 | 132110702 | ENPP1 | -18608 | 3.59529844 | 2.54E-07 |
| Peak22689 | chr12 | 90470692 | 90470920 | ATP2B1 | -368198 | 2.16067112 | 2.55E-07 |
| Peak22689 | chr12 | 90470692 | 90470920 | EPYC | 927997 | 2.16067112 | 2.55E-07 |
